# Learning mutational signatures and their multidimensional genomic properties with TensorSignatures

**DOI:** 10.1101/850453

**Authors:** Harald Vöhringer, Arne van Hoeck, Edwin Cuppen, Moritz Gerstung

## Abstract

Mutational signature analysis is an essential part of the cancer genome analysis toolkit. Conventionally, mutational signature analysis extracts patterns of different mutation types across many cancer genomes. Here we present TensorSignatures, an algorithm to learn mutational signatures jointly across all variant categories and their genomic context. The analysis of 2,778 primary and 3,824 metastatic cancer genomes of the PCAWG consortium and the HMF cohort shows that practically all signatures operate dynamically in response to various genomic and epigenomic states. The analysis pins differential spectra of UV mutagenesis found in active and inactive chromatin to global genome nucleotide excision repair. TensorSignatures accurately characterises transcription-associated mutagenesis, which is detected in 7 different cancer types. The analysis also unmasks replication- and double strand break repair-driven APOBEC mutagenesis, which manifests with differential numbers and length of mutation clusters indicating a differential processivity of the two triggers. As a fourth example, TensorSignatures detects a signature of somatic hypermutation generating highly clustered variants around the transcription start sites of active genes in lymphoid leukaemia, distinct from a more general and less clustered signature of Polη-driven translesion synthesis found in a broad range of cancer types.

**Key findings:**

- Simultaneous inference of mutational signatures across mutation types and genomic features refines signature spectra and defines their genomic determinants.
- Analysis of 6,602 cancer genomes reveals pervasive intra-genomic variation of mutational processes.
- Distinct mutational signatures found in quiescent and active regions of the genome reveal differential repair and mutagenicity of UV- and tobacco-induced DNA damage.
- APOBEC mutagenesis produces two signatures reflecting highly clustered, double strand break repair-initiated and lowly clustered replication-driven mutagenesis, respectively.
- Somatic hypermutation in lymphoid cancers produces a strongly clustered mutational signature localised to transcription start sites, which is distinct from a weakly clustered translesion synthesis signature found in multiple tumour types.

## Introduction

Cancer arises through the accumulation of mutations caused by multiple processes that leave behind distinct patterns of mutations on the DNA. A number of studies have analysed cancer genomes to extract such mutational signatures using computational pattern recognition algorithms such as non-negative matrix factorization (NMF) over catalogues of single nucleotide variants (SNVs) and other mutation types^1–8^. So far, mutational signature analysis has provided more than 50 different single base substitution patterns, indicative of a range of endogenous mutational processes, as well as genetically acquired hypermutation and exogenous mutagen exposures^9^.

Mutational signature analysis via computational pattern recognition draws its strength from detecting recurrent patterns of mutations across catalogues of cancer genomes. As many mutational processes also generate characteristic multi nucleotide variants (MNVs)^10,11^, insertion and deletions (indels)^12–14^, and structural variants (SVs)^6,15–17^ it appears valuable to jointly deconvolve broader mutational catalogues to further understand the multifaceted nature of mutagenesis.

Moreover, it has also been reported that mutagenesis depends on a range of additional genomic properties, such as the transcriptional orientation and the direction of replication^18–20^, and sometimes manifests as local hypermutation (kataegis)^1^. Additionally, epigenetic and local genomic properties can also influence mutation rates and spectra^21–23^. In fact, these phenomena may help to more precisely characterize the underlying mutational processes, but the large number of possible combinations makes the resulting multidimensional data structure unamenable to conventional matrix factorisation methods.

To overcome these challenges, we developed TensorSignatures, a multidimensional tensor factorisation framework, incorporating the aforementioned features for a more comprehensive and robust extraction of mutational signatures using an overdispersed statistical model. We tested the algorithm using simulations and applied it to a dataset comprising 2,778 whole genomes from the International Cancer Genome Consortium (ICGC) Pan Cancer Analysis of Whole Genomes (PCAWG) consortium^24^ spanning 39 cancer types.

The resulting tensor signatures add considerable detail to known mutational signatures in terms of their genomic determinants and broader mutational context. Almost all signatures have contributions from mutation types beyond SNVs and display dynamic activity across the genome. Strikingly, some signatures are being further subdivided based on additional genomic properties, illustrating the differential manifestation of the same mutational process in different parts of the genome. This includes UV-associated mutagenesis in skin cancer, which yields different spectra in regions of active and quiescent chromatin, and possibly also a currently unknown mutational process causing transcription-associated mutagenesis. On the other hand, APOBEC mutations manifest differentially either as predominantly unclustered, replication associated mutations, or highly clustered SV-associated base substitutions. Similarly, mutations caused by polymerase η-driven somatic hypermutation localise into TSS-proximal clusters in lymphoid neoplasms with spectrum distinct from a mostly unclustered, genome-wide pattern found in a range of other cancer types.

Finally, we validate our findings by applying our algorithm to an additional 3,824 metastatic cancer whole genomes from the Hartwig Medical Foundation (HMF), which substantiates the evidence for epigenetic modulation of mutagenesis, and underlines the importance of incorporating additional variant types in mutational signature analysis to uncover complex mutagenic events.

Taken together, TensorSignatures adds great detail and refines mutational signature analysis by jointly learning mutation patterns and their genomic determinants. This sheds light on the manifold influences that underlie mutagenesis and helps to pinpoint mutagenic influences which cannot easily be distinguished based on the mutation spectra alone. TensorSignatures is implemented using the powerful TensorFlow^25^ backend and therefore benefits from GPU acceleration, and can be flexibly tailored. The accompanying code for this work can be found on https://github.com/gerstung-lab/tensorsignatures. or conveniently installed via the Python Package Index (PyPI). TensorSignatures is also available as a web application (http://193.62.55.163/ for the purpose of review please use these anonymous login data: Username: *review,* Password: *test123*) enabling users to fit the set of mutational signatures to user provided samples, and to explore the results of this paper in depth.

## Results

### TensorSignatures jointly decomposes mutation spectra and genomic localisation

#### Multiple mutation types contribute to mutagenesis

Here we analyzed the somatic mutational catalogue of the PCAWG cohort comprising 2,778 curated whole-genomes from 37 different cancer types containing a total of 48,329,388 SNVs, 384,892 MNVs, 2,813,127 deletions, 1,157,263 insertions and 157,371 SVs. We adopted the convention of classifying single base substitutions by expressing the mutated base pair in terms of its pyrimidine equivalent (C>A, C>G, C>T, T>A, T>C and T>G) plus the flanking 5’ and 3’ bases. We categorized other mutation types into 91MNV classes, 62 indel classes, and used the classification of SVs provided by the PCAWG Structural Variants Working Group^17^.

#### Multidimensional genomic features produce a data tensor

Matrix-based mutational signature analysis proved to be powerful in deconvolving mutational spectra into mutational signatures, yet it is limited in characterizing them with regard to their genomic properties. This is because individual mutations cannot always be unambiguously assigned *post hoc* to a given mutational process, which reduces the accuracy of measuring the genomic variation of closely related mutational processes. To overcome this limitation, we use 5 different genomic annotations – transcription and replication strand orientation, nucleosomal occupancy, epigenetic states as well as local hypermutation – and generate 96-dimensional base substitution spectra for each possible combination of these genomic states separately and for each sample. Partitioning variants creates a seven-dimensional count tensor (a multidimensional array), owing to the multitude of possible combinations of different genomic features **(Fig. 1a).**

**Figure 1:**
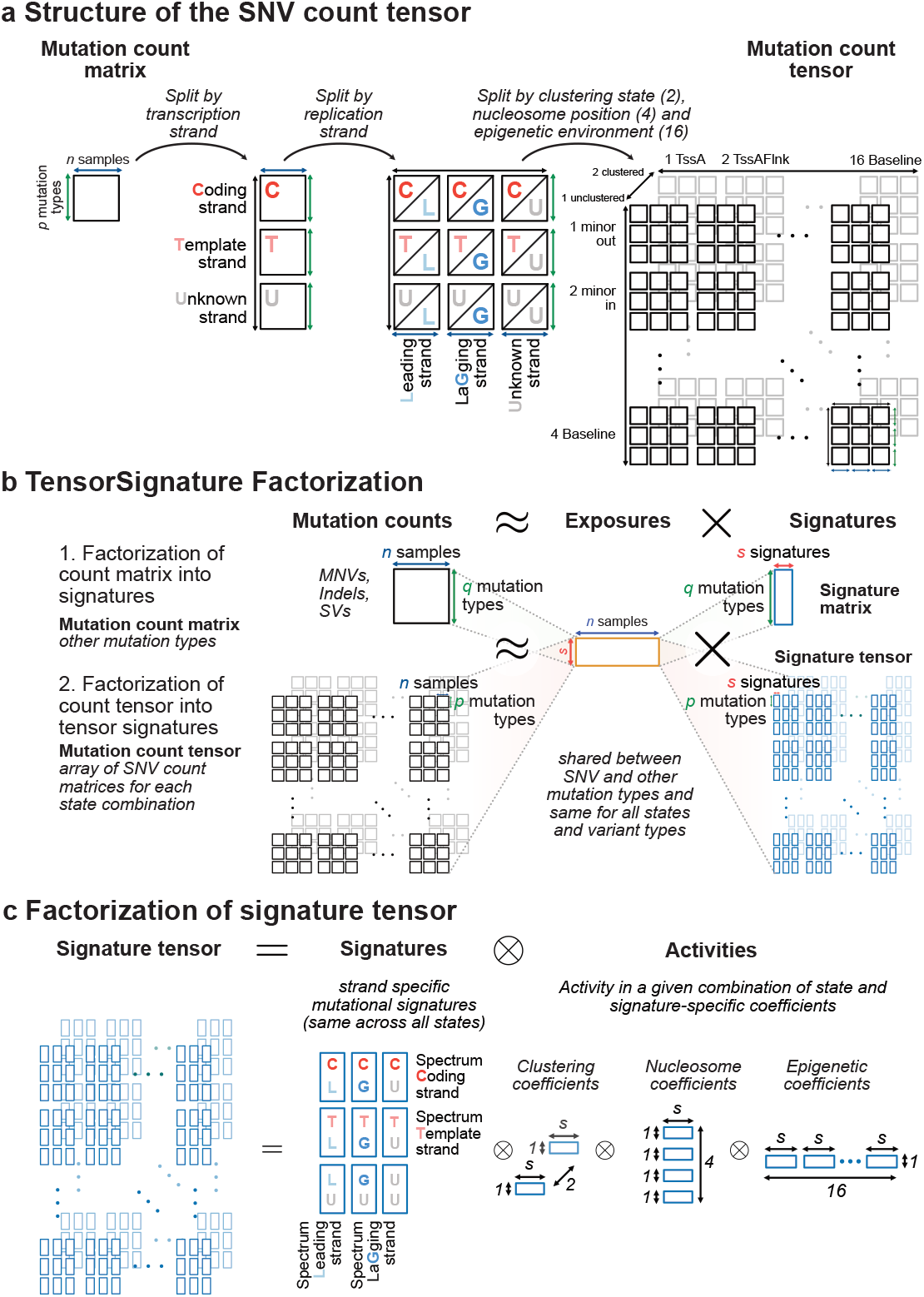
A multidimensional tensor factorization framework to extract mutational signatures, **a,** Splitting variants by transcriptional and replicational strand, and genomic states creates an array of count matrices, a multidimensional tensor, in which each matrix harbours the mutation counts for each possible combination of genomic states, **b,** TensorSignatures factorizes a mutation count tensor (SNVs) into an exposure matrix and signature tensor. Simultaneously, other mutation types (MNVs, indels, SVs), represented as a conventional count matrix are factorised using the same exposure matrix, **c,** The signature tensor has itself a lower dimensional structure, defined by the product of strand-specific signatures, and coefficients reflecting the activity of the mutational process in a given genomic state combination.

#### Directional effects

Mutation rates may differ between template and coding strand, because RNA polymerase II recruits transcription-coupled nucleotide excision repair (TC-NER) upon lesion recognition on transcribed DNA only. Thus, TC-NER leads to lower mutation rates on the template strand, which is best illustrated by UV-induced mutations found in skin cancers^10^. TC-NER usually decreases the number of mutations in highly transcribed genes, but also the opposite effect – transcription associated mutagenesis (TAM) – occurs^18,26^.

Similar to transcriptional strand asymmetries, mutation rates and spectra may differ between leading and lagging strand replication^18,20^. This may be related to the fact that the leading strand is continuously synthesised by DNA polymerase ε, while lagging strand DNA synthesis is conducted by DNA polymerase δ, and is discontinuous due to formation of Okazaki fragments. Therefore, deficiencies in components involved in, or mutational processes interfering with DNA replication may lead to differential mutagenesis on leading or lagging strand.

Since not all mutations can be oriented either due to absent or bidirectional transcription, or because of unknown preferred replication direction far from a replication origin, this creates a total of 3×3 = (template, coding, unknown) x (leading, lagging, unknown) combinations of orientation states in the count tensor **(Fig. 1a).**

#### (Epi-)genomic Localisation factors

Numerous studies found a strong influence of chromatin features on regional mutation rates. Strikingly, these effects range from the 10 bp periodicity on nucleosomes^23^ to the scale of kilo to mega bases caused by the epigenetic state of the genome^21^. To understand how mutational processes manifest on histone-bound DNA, we computed the number of variants on minor groove DNA facing away from and towards histone proteins, and linker DNA between two consecutive nucleosomes. Additionally, we utilized ChromHMM annotations from 127 cell-lines^27^ to annotate genomic regions with epigenetic consensus states, which we used to assign SNVs to epigenetic contexts. Together this adds two dimensions of size 4 and 16 to the count tensor **(Fig. 1a).**

Finally, there are mutational processes capable of introducing large numbers of clustered mutations within confined genomic regions. This phenomenon is termed kataegis^1^ and is thought to be caused by multiple mutational processes^28^. To detect such mutations, we developed a hidden markov model (HMM) to assign the states clustered and unclustered to each mutation based on the inter-mutation distance between consecutive mutations. Separating clustered from unclustered mutations adds the final dimension in the mutation count tensor, which has a total of 6 dimensions with 2 × 576 = 1,152 combinations of states **(Fig. 1a).**

### TensorSignatures learns signatures based on mutation spectra and genomic properties

#### Each sample is modelled as superposition of TensorSignatures

At its core, mutational signature analysis amounts to finding a finite set of prototypical mutation patterns and expressing each sample as a sum of these signatures with different weights reflecting the variable exposures in each sample. Mathematically, this process can be modelled by non-negative matrix factorisation into lower dimensional exposure and signature matrices. TensorSignatures generalises this framework by expressing the (expected value of the) count tensor as a product of an exposure matrix and a signature tensor **(Fig. 1b; Methods).** The key innovation is that the signature tensor itself has a lower dimensional structure, reflecting the effects of different genomic features **(Fig. 1c).** This enables the model to simultaneously learn mutational patterns and their genomic properties by drawing information from the whole dataset even when the number of combinations of genomic states becomes high (1,152), thus yielding a more accurate inference in comparison to conventional NMF relying on a 96-trinucleotide channel decomposition only and subsequent assessment of signature properties, or post-hoc posterior probability calculations **(Fig. S2, S3, Methods).** In this parametrization each signature is represented as a set of 2×2 strand-specific mutation spectra and a set of defined coefficients, measuring its activity in a given genomic state of a given dimension. TensorSignatures incorporates the effect of other variants (MNVs, indels, SVs), which remain unoriented and are expressed as a conventional count matrix, by sharing the same exposure matrix as SNVs, thus enabling to jointly learn mutational processes across different variant classes more robustly in comparison to approaches which rely on (post-hoc) matching mutational spectra **(Fig. S4, Methods).** TensorSignatures models mutation counts with an overdispersed negative binomial distribution, which we tested extensively on simulated data sets **(Fig. S1a-e),** and enables to choose the number of signatures with established statistical model selection criteria, such as the Bayesian Information Criterion (BIC, **Fig. S1f).**

### Mutational signatures are composed of a multitude of mutation types and vary across the genome

#### Discovery analysis of 2,778 genomes produces 20 TensorSignatures

Applying TensorSignatures to the PCAWG dataset and using the conservative BIC **(Fig. S5)** produced 20 tensor signatures (TS) encompassing mutational spectra for SNVs and other mutation types **(Fig. 2a),** and associated genomic properties **(Fig. 2b).** Reassuringly, we extracted a number of signatures with SNV spectra highly similar to the well curated catalogue of COSMIC signatures^9,29^. Interestingly, our analysis revealed a series of signatures that have similar SNV spectra in common, but differ with regard to their genomic properties or mutational composition. These signature splits indicate how mutational processes change across the genome and will be discussed in further detail below. In the following, we refer to signatures via their predominant mutation pattern and associated genomic properties. Of the 20 signatures, 4 were observed in nearly every cancer type: TS01, characterised by C>T mutations in a CpG context, most likely due to spontaneous deamination of 5meC, similar to COSMIC SBS1, TS02 of unknown aetiology, and two signatures with relatively uniform base substitution spectra, TS03 (unknown/quiet chromatin), and TS04 (unknown/active chromatin), which loosely correspond to SBS40 and SBS5.

**Figure 2:**
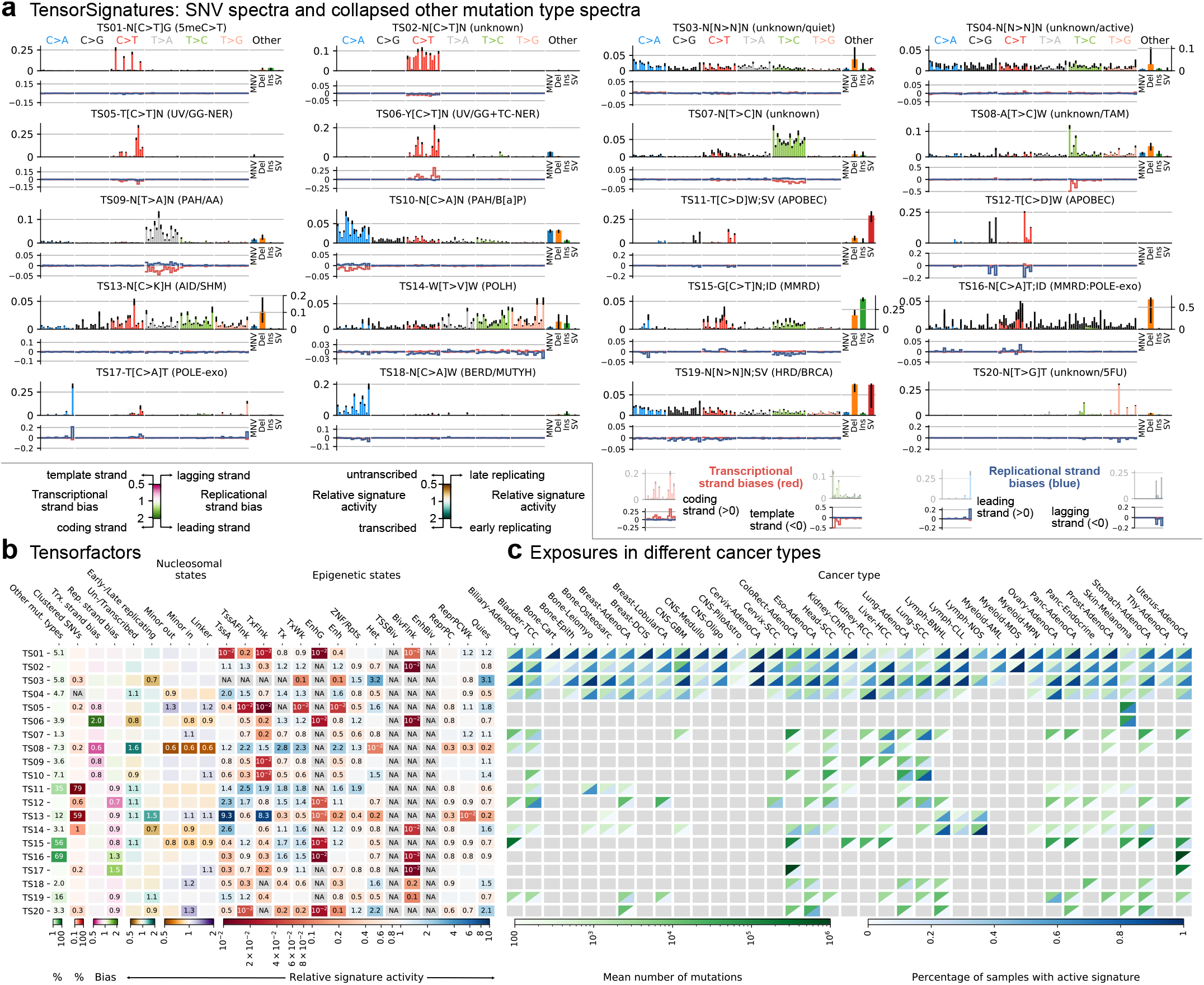
Applying TensorSignatures on 2778 whole genomes from the ICCG PCAWG consortium revealed 20 tensor signatures and their genomic properties, **a,** Upper panels depict SNV spectra, and a summarized representation of associated other mutation types. SNV mutations are shown according to the conventional 96 single base substitution classification based on mutation type in a pyrimidine context (color) and 5’ and 3’ flanking bases (in alphabetical order). The panel under each SNV spectrum indicates transcriptional (red), and replicational strand biases (blue) for each mutation type, in which negative deviations indicate a higher probability for template or lagging strand pyrimidine mutations, and positive amplitudes a larger likelihood for coding or lagging strand pyrimidine mutations (and vice versa for purine mutations), **b,** Heatmap visualization of extracted tensor factors describing the genomic properties of each tensor signature. *Proportions of other mutation types and clustered SNVs* are indicated in percentages. *Transcriptional and replicational strand biases* indicate shifts in the distribution of pyrimidine mutations on coding/template and leading/lagging strand. Coefficients < 1 (pink) indicate signature enrichment on template or lagging strand DNA, and conversely values > 1 (green), a larger mutational burden on coding or leading strand (a value of 1 indicates no transcriptional or replicational bias). *Relative signature activities in transcribed/untranscribed and early/late replicating regions.* Coefficients > 1 (turquoise) indicate enrichment in transcribed and early replicating regions, while values < 1 (brown) indicate a stronger activity of the mutational process in untranscribed or late replicating regions. *Relative signature activities on nucleosomes and linker regions, and across epigenetic states as defined by consensus chromHMM states.* Scores indicate relative signature activity in comparison to genomic baseline activity. A value of 1 means no increase or decrease of a signature’s activity in the particular genomic state, while values > 1 indicate a higher, and values < 1 imply a decreased activity, **c,** *Signature activity in different cancer types (Exposures).* Upper triangles (green) indicate the mean number of mutations contributed by each signature, lower triangles show the percentage of samples with a detectable signal of signature defined as the number of mutations attributed to the signature falling into a signature-specific typical range **(Methods).** Greyed boxes indicate cancer types for which a signature was not found to contribute meaningfully.

#### Signatures are defined by diverse mutation types

While the most prevalent mutations are single base substitutions, there are 16/20 signatures with measurable contributions from other mutation types (> 1%; **Fig. 2b**). The most notable cases are TS15, which is similar to a compound of COSMIC signatures SBS6/15/26 + ID1/2 and characterised by C>T transversions in a GCN context and frequent mononucleotide repeat indels indicative of mismatch repair deficiency (MMRD). Similarly, TS16, likely to reflect concurrent MMRD and POLE exonuclease deficiency, exhibits large probabilities for deletions and a base substitution pattern similar to SBS14. Large proportions of SVs (~25 %) were found in TS11, which reflects SV-associated APOBEC mutagenesis caused by double strand break repair with a base substitution spectrum similar to SBS2/13. Furthermore, TS19 apparently reflects a pattern of homologous recombination deficiency (HRD), characterised by a relatively uniform base substitution pattern similar to SBS3, but a high frequency of SVs, in particular tandem duplications **(Supplementary note Fig. 93).**

9/20 signatures displayed a measurable propensity to generate clustered mutations (>0.1%; **Fig. 2b).** The proportions of clustered mutations produced by each mutational process were highest in signatures associated with APOBEC and activation-induced deaminase (AID) activity: Up to 79% and 0.6% of SNVs attributed to TS11 and TS12, respectively, were clustered, with otherwise indistinguishable base substitution spectra. A similar phenomenon was observed in two signatures reflecting Polη driven somatic hypermutation (SHM). While both TS13 and TS14 have only mildly diverging base substitution spectra, with TS14 being similar to SBS9, they dramatically differ in the rates at which they generate clustered mutations, which are 59% and 1%, respectively **(Fig. 2b).**

#### Replication and Transcription strand biases

5/20 signatures exhibit substantial transcriptional strand bias (TSB ≥ 10%; **Fig. 2b).** This is strongest in the UV-associated signature TS06, similar to SBS7b, where the rate of C>T substitutions on the template strand was half of the corresponding value on the coding strand, highly indicative for active TC-NER. In contrast, TS08, similar to SBS16, shows largest activities in liver cancers and preferably produces T>C transitions on template strand DNA. In line with a transcription-coupled role, the activity of TS08 shows a noteworthy elevation in transcribed regions. Both signatures will be discussed in more detail later on.

Analysis of pyrimidine/purine shifts in relation to the direction of replication indicated 9/20 signatures with replication strand biases (RSB ≥ 10%). In accordance with previous studies, TS12 asserts a higher prevalence of APOBEC-associated C>D mutations, consistent with cytosine deamination, on lagging strand DNA which is thought to be exposed for longer periods as opposed to more processively synthesized leading strand DNA. Conversely, TS17, associated with POLE exonuclease variants (SBS10a/b), displays a pyrimidine bias towards the leading strand^18^ **(Fig. 2b).** Since DNA polymerase ε performs leading strand synthesis, the strand bias indicates that C>A (G>T) mutations arise on a template C, presumably through C·dT misincorporation^30^. Further examples with replication strand biases include the MMRD-associated signatures TS15 and TS16 discussed above. Of note, the two SHM-associated signatures TS13 and TS14 displayed opposing patterns with respect to their activity in oriented (early) and unoriented (late) replicating regions **(Fig. 2b).**

#### Genomic properties modulate signatures, with epigenetic states having the greatest influence

To understand how mutational processes manifest on nucleosomal DNA, we estimated signature activities on minor groove DNA facing away from and towards histone proteins, and linker DNA between two consecutive nucleosomes **(Fig. 2b).** Almost all signatures showed either an increase or a decrease of mutational rates across all nucleosomal states. The only exception to this rule is TS20 (SBS17a,b), which showed a slight decrease in the outward facing minor groove, while the inwards facing showed elevated mutation rates^23^.

Considering the activities of mutational processes across epigenetic domains, our analysis indicates that there is not a single mutational process which is acting uniformly on the genome **(Fig. 2b).** However, our results suggest that mutational processes may be categorized into two broad groups: Those that are elevated in active (TssA, TssAFlnk, TxFlnk, Tx and TxWk) and depleted in quiescent regions (Het, Quies), and vice versa. This phenomenon includes the two omnipresent signatures with relatively uniform spectra TS03 and TS04, suggesting a mechanism associated with the chromatin state behind their differential manifestation **(Fig. 2a).** This also applies to two signatures associated with UV exposure, TS05 and TS06, and also two signatures of unknown aetiology, most prominently found in Liver cancers, TS07, similar to SBS12, and TS08, which we will discuss in detail in the following section.

#### Validation of TensorSignatures

The aforementioned observations were replicated in a fully independent second cohort of whole genomes from the Hartwig Medical Foundation with 3,824 samples from 31 cancers encompassing 95,531,862 SNVs, 1,628,116 MNVs, 9,228,261 deletions, 5,408,915 insertions and 1,001,433 structural variants^31^. Applying TensorSignatures to this data set produced 27 tensor signatures **(Fig. S6a).** Of these 10 closely resembled (cosine distance < 0.2) signatures of the discovery analysis with closely matching genomic activity coefficients **(Fig. 3d, S6b).** These include the signatures of spontaneous deamination TS01, the two signatures of UV mutagenesis TS05/06, SV-associated APOBEC mutagenesis TS11, as well as signatures of MMRD TS16, *POLE^exo^* mutations TS17, as well as MUTYH deficiency TS18, HRD TS19 and TS20.

**Figure 3:**
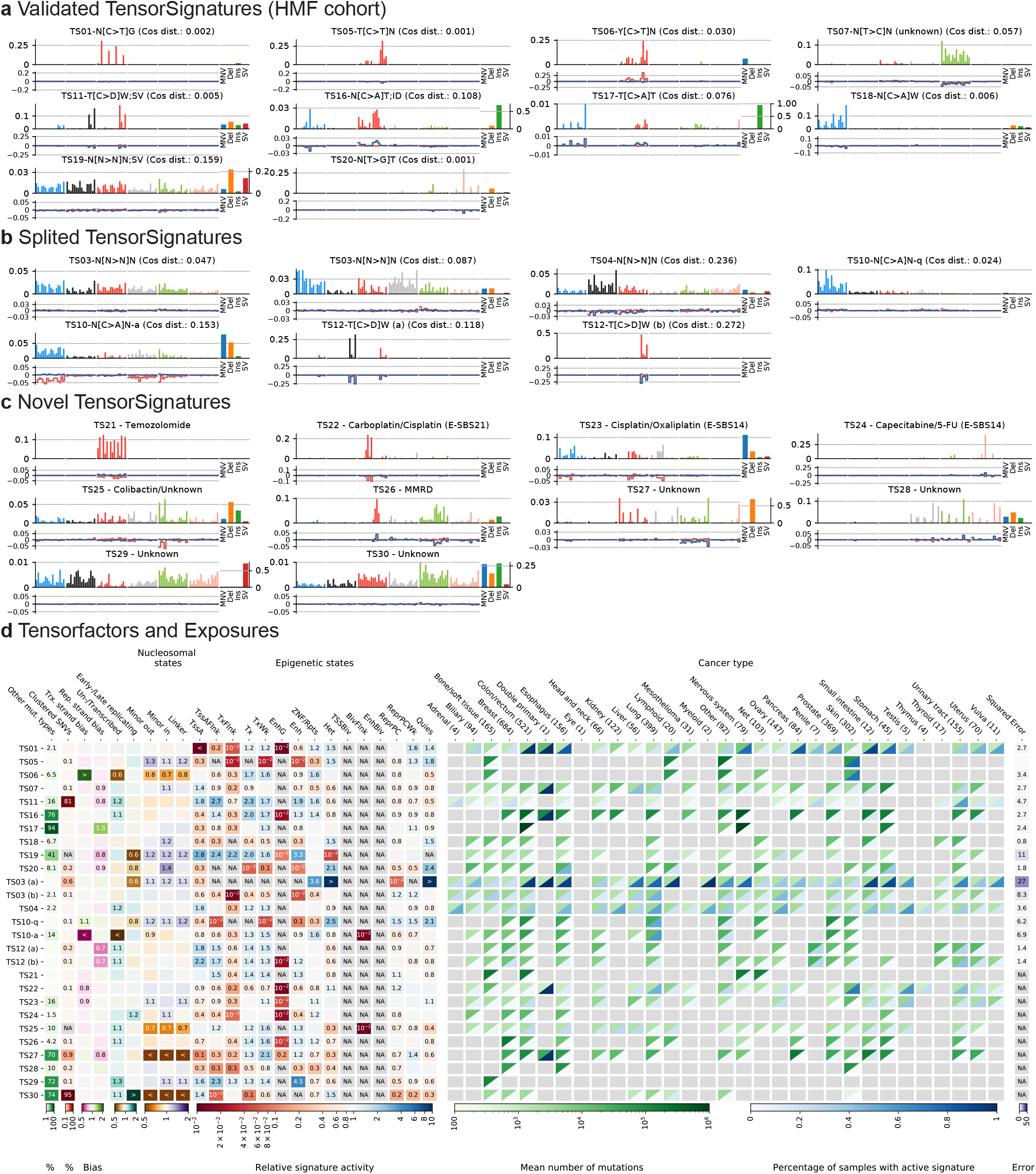
Tensor signatures of the HMF cohort, **a,** Validated tensor signatures with high similarity (indicated as cosine distance) to the mutational processes extracted in our discovery analysis using PCAWG data, **b,** TensorSignature splits that seemingly represent derivatives of tensor signature TS03, TS04, TS10 and TS12. **c,** Novel tensor signatures of the HMF cohort, **d,** Extracted tensor factors, exposures and summed squared errors of tensor factors from the discovery and validation analysis.

A further 7 signatures seemingly constitute splits of tensor signatures from the PCAWG cohort **(Fig. 3b).** A complex three-way split appeared to occur for TS03 and TS04, which were found in a broad range of cancer types. One of the derivative signatures resembles the mutation spectrum of SBS8 from the COSMIC catalogue, however without measurable transcriptional strand bias. A second derived signature is similar to SBS39; our analysis reveals replication strand bias for C>G variants and a potentially wider range of cancer types for both signatures. Further, signature TS12, resembling replication associated APOBEC mutagenesis, split into two signatures with base substitution spectra similar to SBS2 (C>T) and SBS13 (C>G), but preserving the strong replication strand bias. Lastly, a split of TS10, likely attributed to mutagens included in tobacco smoke, was observed.

Finally a set of 10 novel signatures without close match to those in the PCAWG cohort was found **(Fig 3c).** This includes five spectra linked to cancer therapies, illustrating the additional insights on preceding therapies provided by the HMF metastatic cancer cohort. TS21 is characteristic of treatment with the methylating agent temozolomide (SBS11); the observed transcriptional strand bias reflects a higher rate of G>A mutations on the coding strand (equivalent to higher rates of C>T on the template strand), consistent with methyl guanine being removed by TC-NER in the absence of *MGMT.* TS22 and TS23 have been previously associated with cisplatin (termed E-SBS21 and E-SBS14)^32,33^. While both signatures exhibit mild transcriptional strand biases, only TS23 shows a strong association with MNVs going in line with the propensity of cisplatin/oxaliplatin to form intrastrand DNA adducts **(Fig. S6c).** TS24 displays the characteristics of treatment with 5-FU, which inhibits thymine synthesis and has been proposed to be mutagenic via genomic fluorouracil incorporation^33^. TS28, with similarity to SBS41, was only found in two samples, possibly due to treatment with the experimental drug SYD985, which consists of a duocarmycin-based HER2-targeting antibody-drug conjugate^31^.

Further, TensorSignatures detected a signature of colibactin, TS25, which has been previously characterised^32,34^. TS25 displays contributions of MNVs and short indels, activity in active genomic regions and concomitant transcriptional strand bias of T>C mutations **(Fig. 3c,d).** TS26’s indels and similarity to SBS15 suggests an association with MMRD; TS27 has an unknown aetiology and displays strong replicational strand bias. The large proportion of structural variants and the flat SNV spectrum of TS29 may represent non-specific mutagenesis at SVs. TS30 was found in lymphoid and other cancers and had a high proportion of clustered mutations, similar, but not identical to TS14 **(Fig. 3d).**

### Differential mutagenesis in quiet and active chromatin

Mutational signatures are an abstraction of the observed mutational spectra casting light on the underlying processes. As TensorSignatures provides additional clues about the aetiology of diverse processes, we explore some of the genomic activity patterns and properties of selected TensorSignatures in further detail. An emerging feature was differential mutagenesis in active and quiet areas of the genome.

#### The spectrum of UV mutagenesis changes from closed to open chromatin, reflecting GG- and TC-NER

Two signatures, TS05 and TS06, were exclusively occurring in Skin cancers of both cohorts and displayed almost perfect correlation (Spearman *R*^2^=0.98, **Fig. S7a**) of attributed mutations, strongly suggesting UV mutagenesis as their common cause. Both signatures share a very similar SNV spectrum, only differing in the relative extent of C[C>T]N and T[C>T]N mutations, which is more balanced in TS06 **(Fig. 2a).** However, they strongly diverge in their activities for epigenetic contexts and transcriptional strand biases: TS05 is enriched in quiescent regions, and shows no transcriptional strand bias, while the opposite is true for TS06, which is mostly operating in active chromatin **(Fig. 2b).** Of note, the spectra of these signatures closely resemble that of COSMIC SBS7a and SBS7b, which have been suggested to be linked to different classes of UV damage^35^. However, as our genomically informed TensorSignature inference and further analysis show, the cause for the signature divergence may be found in the epigenetic context, which seemingly not only determines mutation rates, but also the resulting mutational spectra.

A characteristic difference between the two signatures is the presence of a strong transcriptional strand bias in signature TS06, which is almost entirely absent in signature TS05 **(Fig. 4a).** To verify that this signature inference is correct, and the observed bias and spectra are genuinely reflecting the differences between active and quiescent chromatin, we pooled C>T variants from Skin-Melanoma samples which revealed that the data closely resembled predicted spectra **(Fig. 4b).** In addition, quiescent chromatin also displays a predominant T[C>T]N substitution spectrum (5’C/5’T=0.3), while the spectrum in active chromatin is closer to Y[C>T]N (5’C/5’T=0.58), as predicted by the signature inference **(Fig. 4a).** This difference does not appear to be related to the genomic composition, and holds true even when adjusting for the heptanucleotide context **(Fig. S7b).**

**Figure 4:**
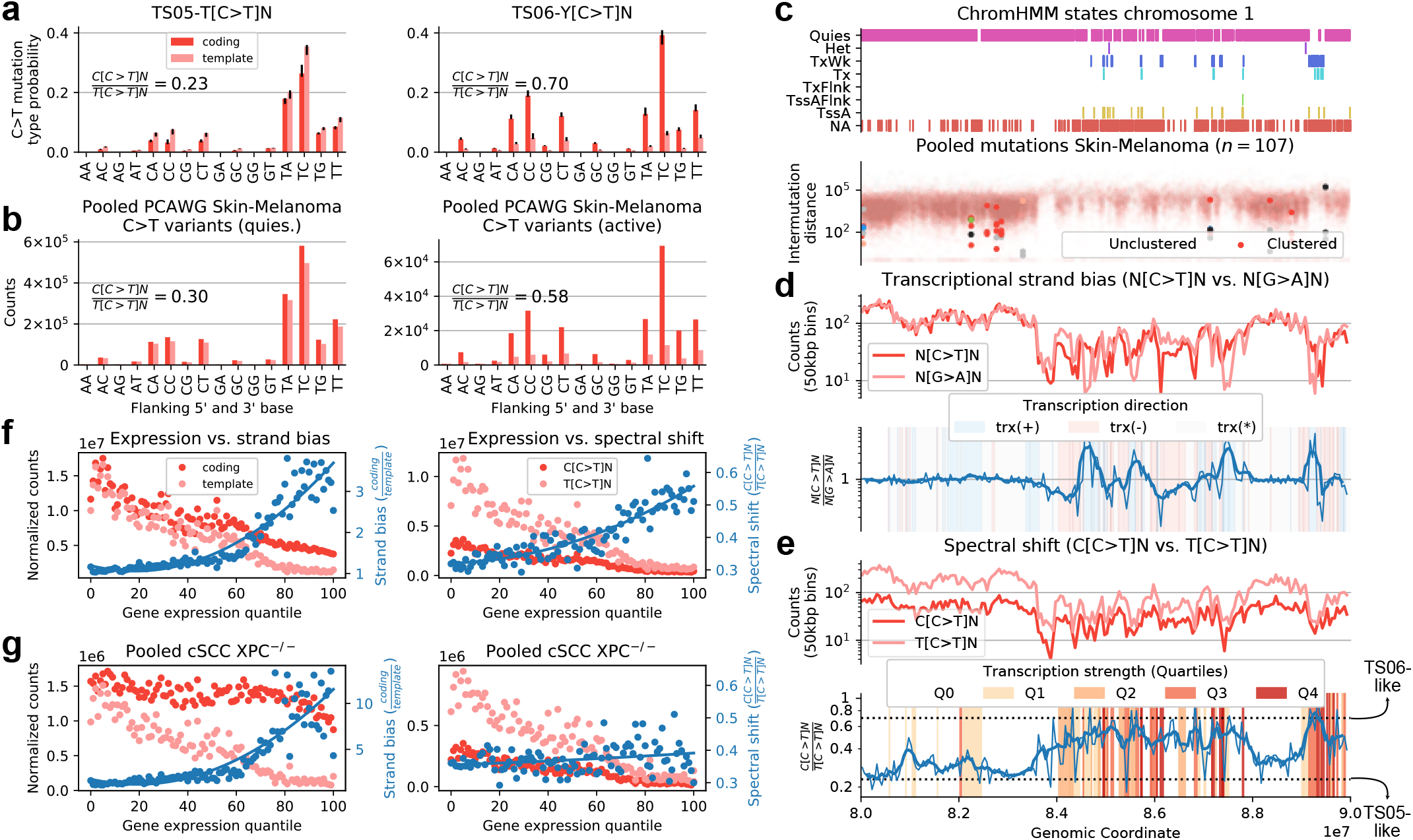
The spectrum of UV mutagenesis changes from open to closed chromatin, **a,** C>T mutation probabilities of TensorSignatures TS05 and TS06 for coding and template strand DNA. **b,** Pooled PCAWG Skin-Melanoma C>T variant counts from coding and template strand DNA in epigenetically active (TssA, TssAFlnk, TxFlnk, Tx and TxWk, right) and quiescent regions (Het and Quies, left), **c,** Consensus ChromHMM states from a representative 10 Mbp region on chromosome 1, and the corresponding mutational density of pooled Skin-Melanoma samples, **d,** N[C>T]N and N[G>A]N counts in 50kbp bins, and their respective ratios (thin blue line: ratio; thick blue line: rolling average over 5 consecutive bins) illustrate the transcriptional strand bias of C>T mutations in quiescent and active regions of the genome, **e,** Relationship between expression strength and the spectral shift of C>T mutations in terms of binned C>T variant counts in TpC and CpC context and their respective ratios (thin blue line) as well as a rolling average (thick blue line), **f,** Gene expression strength vs. transcriptional strand bias (measured by the ratio normalized C>T variants in Skin-Melanoma on coding and template strand), and gene expression strength vs. C[C>T]/T[C>T] spectral shift (indicated as the ratio of normalize C>T mutations in 5’C and 5’T context), **g,** Transcriptional strand bias and C[C>T]/T[C>T] spectral shift in GG-NER deficient XPC^-/-^ cSCC genomes. Blue curves: quadratic fit.

To illustrate how the mutation spectrum changes dynamically along the genome in response to the epigenetic context, we selected a representative 10 Mbp region from chromosome 1 comprising a quiescent and active genomic region as judged by consensus ChromHMM states, and the varying mutational density from pooled Skin-Melanoma samples **(Fig. 4c).** As expected, actively transcribed regions display a strong transcriptional strand bias **(Fig. 4d).** Further, this change is also accompanied by a change of the mutation spectrum from a T[C>T]N pattern to a Y[C>T]N pattern with the ratios indicated by our TensorSignature inference **(Fig. 4e).**

These observations are further corroborated by RNA-seq data available for a subset of samples **(n=11):** The transcriptional strand bias is most pronounced in expression percentiles greater than 50 leading to an increased ratio of coding to template strand mutations **(Fig. 4f**). Again, the decline is accompanied by a shift in the mutation spectrum: While both C[C>T]N and T[C>T]N variant counts decline steadily as gene expression increases, the reduction of C[C>T]N mutations is larger in comparison to T[C>T]N mutations, which manifests as an increasing C[C>T]N and T[C>T]N ratio, reaching a ratio of approximately 0.5 in the highest expression quantiles **(Fig. 4f**).

The diverging activity in relation to the chromatin state suggests an underlying differential repair activity. Global genome nucleotide excision repair (GG-NER) clears the vast majority of UV-lesions in quiescent and active regions of the genome and is triggered by different damage-sensing proteins. Conversely, TC-NER is activated by template strand DNA lesions of actively transcribed genes. As TS05 is found in quiescent parts of the genome, it appears likely that it reflects the mutation spectrum of UV damage as repaired by GG-NER. Based on the activity of TS06 in actively transcribed regions and its transcriptional strand bias, it seemingly reflects the effects of a combination of GG- and TC-NER, which are both operating in active chromatin. This joint activity also explains the fact that the spectrum of TS06 is found on *both* template and coding strands.

This attribution is further supported by data from *n*=13 cutaneous squamous cell carcinomas (cSCCs) of *n*=5 patients with Xeroderma Pigmentosum, group C, who are deficient of GG-NER and п=8 sporadic cases which are GG-NER proficient^36^. XPC/GG-NER deficiency leads to an absence of TS05 in quiescent chromatin and to a mutation spectrum that is nearly identical in active and quiescent regions of the genome **(Fig. S7C).** Furthermore, the UV mutation spectrum of XPC/GG-NER deficiency, which is thought to be compensated by TC-NER, differs from that of TS06, reinforcing the notion that TS06 is a joint product of GG- and TC-NER. This is further supported by the observation that XPC/GG-NER deficiency leads to a near constant coding strand mutation rate, independent of transcription strength^36^ **(Fig. F4g**), indicating that the transcriptional dependence of coding strand mutations in GG-NER proficient melanomas and cSCCs is due to transcriptionally facilitated GG-NER.

While the activity patterns of TS05 and TS06 appear to be well aligned with GG-NER and GG/TC-NER, these observations, however, do not explain the observed differences in mutation spectra. The fact that the rates of C[C>T]N and T[C>T]N mutations change between active and quiescent chromatin – and the fact the these differences vanish under XPC/GG-NER deficiency – suggests that DNA damage recognition of CC and TC cyclobutane pyrimidine dimers by GG-NER differs between active and quiescent chromatin, with relatively lower efficiency of TC repair in quiescent genomic regions, as evidenced by TS05.

#### TC-NER changes the mutation spectrum of tobacco-associated mutations

A similar split of an exogenous mutational signature into quiet and active chromatin was observed in lung cancers of the HMF cohort where TS10 splits into two signatures, HMF TS10-q, which shows largest activity in heterochromatin, while HMF TS10-a is enriched in actively transcribed regions, and exhibits a strong transcriptional strand bias with lower rates of C>A changes on the coding strand, equivalent to G>T transversions on the template strand **(Fig. 3b, 3d, 5a).** This strand bias has been attributed to TC-NER removing benzo[a]pyrene derived adducts on guanines from the template strand^37^.

**Figure 5:**
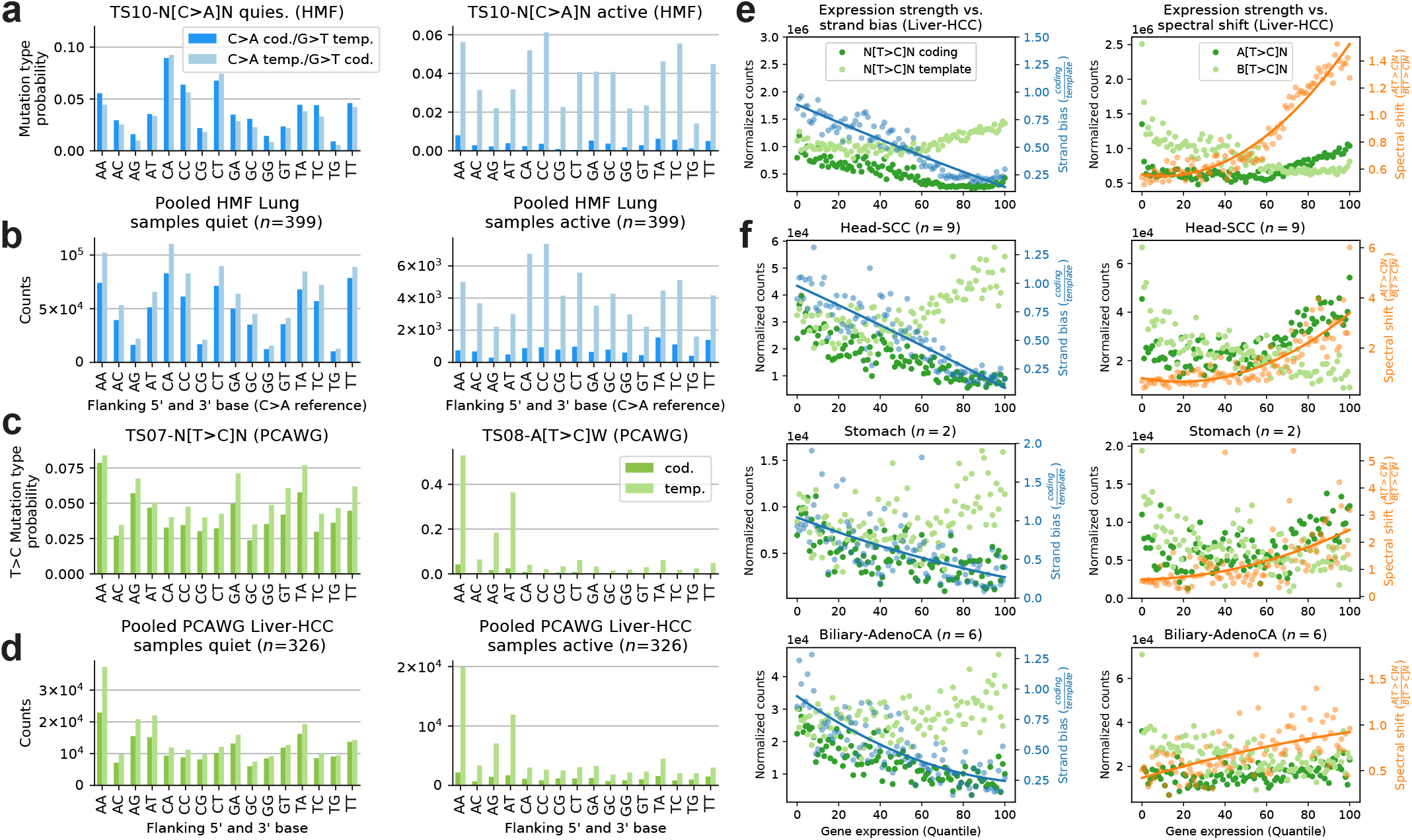
Genomically dependent T>C mutagenesis in Liver-HCC and other cancer types, **a,** C>A mutation type probabilities of HMF TS10-q and TS10-a for coding and template strand DNA. **b** Pooled HMF lung samples C>A variant counts from coding and template strand DNA in quiescent (Het and Quies, left) and epigenetically active regions (TssA, TssAFlnk, TxFlnk, Tx and TxWk, right), **c,** T>C mutation type probabilities of PCAWG TS07 and TS08 for coding and template strand DNA. **d,** Pooled PCAWG Liver-HCC T>C variant counts for coding and template strand DNA in epigenetically active and quiescent regions, **e** and **f,** Transcriptional strand bias and A[T>C]/B[T>C] spectral shift in samples from different cancers with TS07 and TS08 contributions. Lines correspond to quadratic fits.

The emergence of two mutational signatures indicates that this repair process also changes the mutation spectrum. The suggested split is also evident in pooled mutations from HMF lung cancers in quiescent and active genomic regions, respectively, revealing that predicted spectra coincide with corresponding tensor signatures HMF TS 10-q and TS10-a **(Fig. 5b)**.

The C>A (G>T) mutation spectrum observed in quiescent regions, extracted by HMF TS10-q, displays highest rates of mutations in a CCN (NGG) context **(Fig. 5a)**. Interestingly, the same pattern is also observed in actively transcribed regions for C>A on the template strand, equivalent to G>T mutations on the coding strand. This is in contrast to the C>A coding strand pattern, and HMF TS10-a, for which this difference is largely eroded. These observations reflect how TC-NER removes genotoxic guanine adducts from the template strand, which leads to lower mutation rates and also a more homogeneous base context of G>T mutations. The differentential mutation spectrum indicates that either the efficiency of TC-NER – or the mutagenicity of residual genomic alterations – differs depending on the base context, analogous to observations in UV-induced mutagenesis. The result being that the mutation types and rates caused by tobacco-associated carcinogens differ between coding and template strand in transcribed regions and also to different mutation spectra in quiescent and active genomic regions.

#### Transcription-associated mutagenesis manifests in an ApT context in highly transcribed genes

A third split of mutational signatures between active and quiet regions was observed in Liver and other cancer types **(Fig. 2b,c),** driven by differential activity of TS07 and TS08, which closely resemble COSMIC signatures SBS12 and SBS16, respectively. In line with previous findings, there was a strong transcriptional bias of TS08, introducing 1.6× more T>C variants on the template strand **(Fig. 2b).** While both signatures are most frequently found in Liver cancers, where they are strongly correlated (*R*^2^=0.68, **Fig. S8a),** they are also observed in a range of other cancers, indicating that they are reflecting endogenous mutagenic processes.

The most prominent difference between these signatures is the depletion of mutation types in 5’-B context on coding strand DNA in TS08 **(Fig. 5c;** B = C, G, or T). Signature TS08 displays a strong transcriptional strand bias, as previously noted for SBS16^26^, and is confirmed here by a direct investigation of variant counts **(Fig. 5d).** A further defining feature of TS08 are indels >2bp **(Fig. 2a, Supplementary Note Fig. 38),** which were reported to frequently occur in highly expressed lineage-specific genes in cancer^12^, consistent with experimental data of transcription-replication collisions^38^. In line with this, mutation rates showed a dynamic relation to transcriptional strength **(Fig. 5e).** Normalized counts of T>C mutations on coding and template strand initially decline for low transcription. Yet this trend only continues on the coding strand for transcription quantiles (>50), but reverses on the template strand, producing more N[T>C]N, and most commonly A[T>C]N, mutations the higher the transcription, in line with previous reports of TAM^18^.

While this effect is most common in Liver-HCC samples, where it has been described in detail, it has been observed that SBS5, one of three broadly active signatures, displays signs of potential contamination by SBS16/TS08 in the absence of further intra-genomic stratification. Accordingly, a genomically informed analysis by TensorSignatures also discovers this signature in highly transcribed genes of Head-SCC, Stomach-AdenoCa and Biliary-AdenoCa **(Fig. 5f, S8b),** showing that A[T>C]WTAM and N[T>C]N mutagenesis in heterochromatic regions occur in a broad range of cancers.

### Replication- and DSBR-driven APOBEC mutagenesis

In the following, we turn our focus to TS11 and TS12, which both share a base substitution spectrum attributed to APOBEC mutagenesis, but differ greatly with regard to their replicational strand bias, broader mutational composition, and clustering properties. While TS12 is dominated by SNVs (99%) with strong replicational strand bias, SNVs in TS11 make up only 64% of the overall spectrum and are highly clustered. The rest of the spectrum is mostly dominated by structural variants **(Fig. 2a, 6a, Supplementary note Fig. 53).** This signature split reveals two independent triggers of APOBEC mutagenesis, which is thought to require single stranded DNA as a substrate, present either during lagging strand replication, or double strand break repair (DSBR). In the following, we will further assess the genomic properties of these two different modes of action.

**Figure 6:**
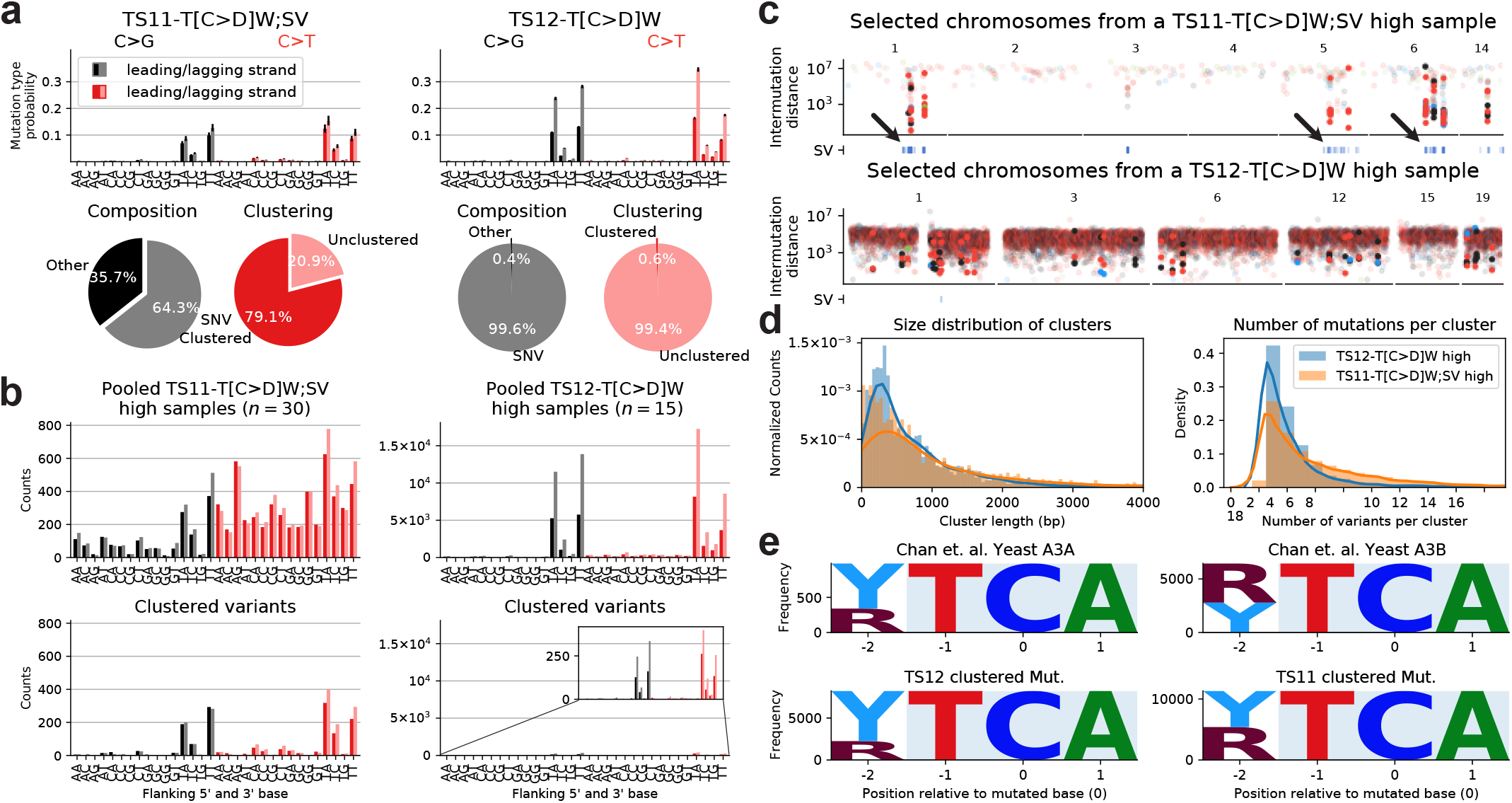
Double-strand break and replication induced APOBEC mutagenesis, **a,** C>G and C>T spectra of TS11 and TS12 for leading and lagging strand DNA. Pie charts underneath indicate percentages of clustered mutations and the contribution of other mutation types in TS11 and TS12. **b,** Observed unclustered (top) and clustered variants (bottom) in TS11 and TS12 high samples, **c,** Rainfall plots with SV annotations from a typical sample with high TS11 (top) and TS12 contributions (bottom). **d,** Size distribution of mutation clusters (consecutive clustered mutations), and the distribution of number of variants per mutation cluster in TS11 and TS12 high samples respectively. Curves depict corresponding kernel density estimates, **e,** Motif logo plots of the tetranucleotide context at mutated TCA sites in yeast cells exposed to APOBEC3A/3B mutagenesis respectively (Chan et. al.), and similar motif logo plots extracted at clustered mutations from samples with high TS11 or TS12 exposures.

To verify the split, we pooled C>G and C>T variants from 30 and 15 samples with high TS11 and TS12 exposures, respectively (TS11 and TS12 contributions > 10 % and 70 % respectively, **Fig. 6b).** We noticed that the spectrum in TS12-high samples was clearly dominated by T[C>D]N mutations, whereas the distribution in TS11-high samples was cross-contaminated by other mutational processes. However, assessment of replicational strand biases revealed that lagging strand mutations were twice as large as leading strand mutations in TS12-high samples, but not in TS11-high samples. Moreover, the proportion of clustered variants in TS12-high samples was much lower than in TS12-high in line with the signature inference **(Fig. 6b).**

The association of TS11 with structural variants suggests clustered APOBEC mutagenesis at sites of DNA double strand break events. This is confirmed by the spatial co-occurrence of SVs and clustered mutations (a feature not directly measured by TensorSignatures; **Fig. 6c).** Furthermore, SV-proximal clustered variants do not display a replicational strand bias, adding further weight to the notion that these arise in a DSBR-driven, replication-independent manner **(Fig. S9a).** Interestingly, SV-distal clusters displayed, on average, only a very weak replicational strand bias, indicating that the majority of these foci arose in a replication-independent fashion, presumably during successful DSBR, which did not create SVs.

Next, we assessed whether differences exist in the characteristics of clustered variants, beyond the fact that these are much more frequent in DSBR driven mutagenesis. To this end, we pooled clustered variants from TS11/ 12-high samples and computed their size distribution, which revealed that the length of mutation clusters tend to be larger at SVs (Median 717 vs. 490bp, **Fig. 6d).** This goes in line with the observation that clustered mutations at DSBRs tend to have more mutations per cluster (Median 5 vs 4 variants; **Fig. 6d).**

Differential size distributions of TSıı/12 mutation clusters raise the question if an APOBEC subtype-specific activity is underlying their distinct genomic manifestation. Previous studies linked the motifs YT[C>T]A and RT[C>T]A to APOBEC3A and APOBEC3B mutagenesis respectively^39^. To test whether TS11 or TS12 clusters may be linked one or the other isoform, we extracted the tetranucleotide context at clustered T[C>T]A sites from samples with high TS11 or TS12 contributions. Clustered TS12 mutations comprise only a small fraction of purines, while this proportion increases to approximately 50 % in TS11 samples. These findings may indicate larger contributions of APOBEC3A and 3B in TS12 and TS11 samples respectively **(Fig. 6e, S9b).** To confirm this, we assessed samples that harboured a germline copy number polymorphism that effectively deletes APOBEC3B^40^. This analysis was unfortunately inconclusive, as the subset of PCAWG samples for which this annotation was available, did not show any activity for TS11.

Finally we note that, using data from the HMF cohort, TS12, was further split into C>T and C>G akin to mutational signatures SBS2 and 13 from the COSMIC catalogue **(Fig. 3b).**

These two signatures were attributed to differential activity of OGG/UNG-driven base excision repair of uracil created by APOBEC-induced cytosine deamination^41^. The observation that this split occurs for TS12 rather than TS11, suggests that repair of replication-driven APOBEC deamination, possibly by APOBEC3A, is subject to higher variation in downstream BER than DSBR-driven APOBEC mutagenesis.

Taken together, these results indicate that there are two distinct triggers of APOBEC mutagenesis, induced by DSBR or replication. Higher rates, longer stretches and larger proportions of RT[C>T]A APOBEC mutation clusters in the vicinity of SVs, as evidenced by TS11, suggests that DSBR leads to larger and possibly longer exposed stretches of single-stranded DNA possibly due to A3B. Conversely, lower rates, shorter stretches and a high fraction of YT[C>T]A mutation clusters of TS12 in conjunction with a strong replicational strand bias indicate A3A mutagenesis during lagging strand synthesis, which is more processive than DSBR, allowing for fewer and shorter mutation clusters only.

### Clustered somatic hypermutation at TSS and dispersed SHM

Two other TensorSignatures produced substantial amounts of clustered variants with, but different epigenomic localisation. TS13 showed largest activities in lymphoid cancers and produced **60%** clustered variants **(Fig. 2b).** The SNV spectrum resembles the c-AID signature reported previously^7^, suggesting an association with activation-induced cytidine deaminases (AID), which initiates somatic hypermutation in immunoglobulin genes of germinal center B cells. Like its homolog APOBEC, AID deaminates cytosines within single stranded DNA, although it targets temporarily unwound DNA in actively transcribed genes, rather than lagging strand DNA or DSBRs^42,43^.

TensorSignatures analysis reveals that TS13 activity is 9x and 8x enriched at active transcription start sites (TssA) and flanking transcription sites (TxFlnk, **Fig. 2b),** respectively. To illustrate this, we pooled single base substitutions from Lymph-BHNL samples and identified mutational hotspots by counting mutations in 10 kbp bins **(Fig. 7a, b).** Inspection of hotspots confirmed that clustered mutations often fell accurately into genomic regions assigned as TssA **(Fig. 7c).** The aggregated clustered mutation spectrum in TssA/TxFlnk regions across lymphoid neoplasms (Lymph-BNHL/CLL/NOS, п=202) indeed showed high similarity to TS13, possibly with an even more pronounced rate of C>K (K=G or T) variants similar to SBS84^9^ **(Fig. 7d).** Conversely, the clustered mutational spectrum from all other epigenetic regions was characterized by a larger proportion of T>C and T>G mutations, similar to TS14, which only produces about 1% clustered mutations and closely resembles SBS9, attributed to Polη-driven translesion synthesis (TLS) during somatic hypermutation.

**Figure 7:**
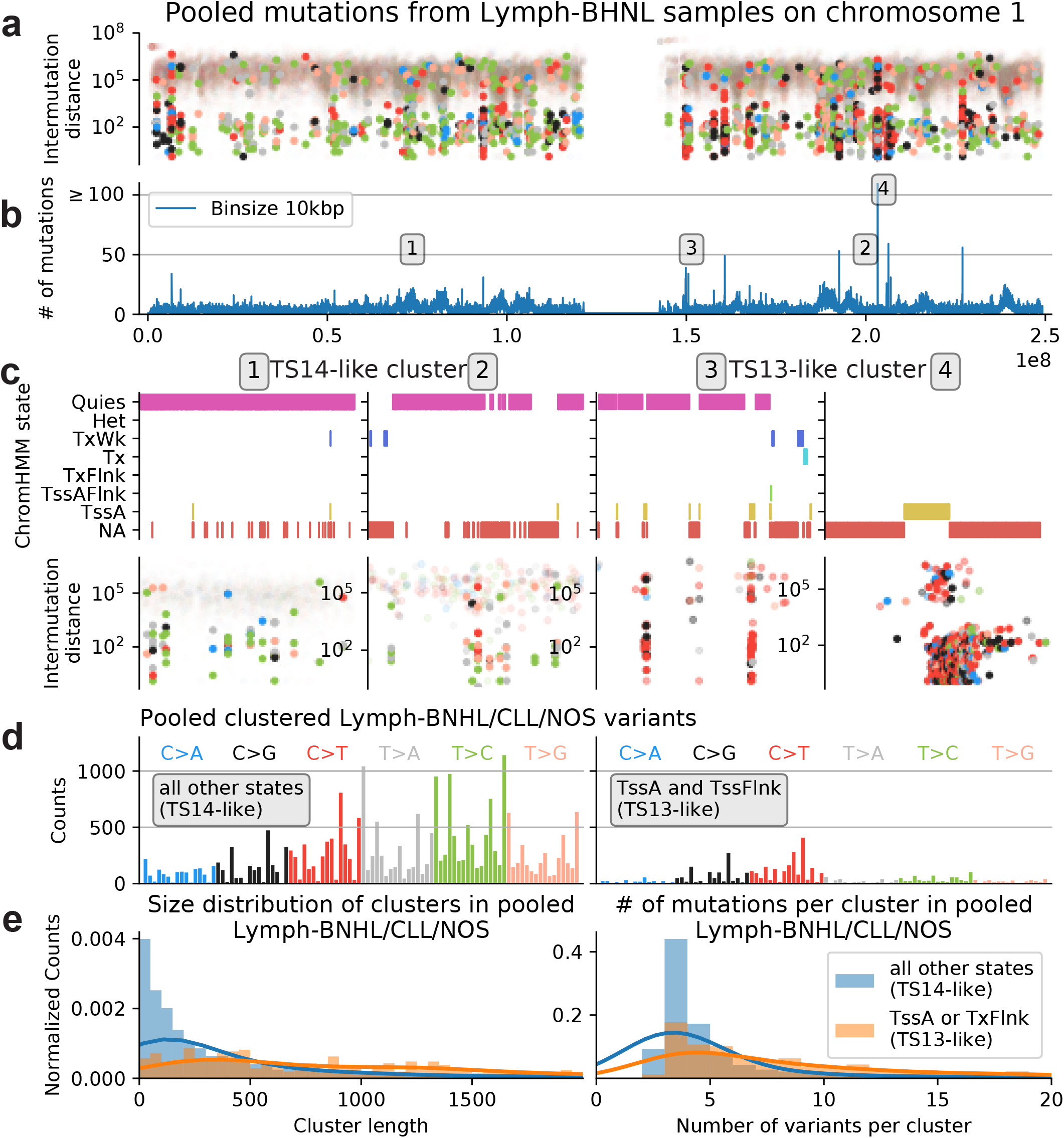
Identification of a highly clustered mutational signature at active TSS. **a,** Rainfall plot of pooled variants from Lymph-BHNL samples on chromosome 1 (highlighted dots indicate clustered mutations), **b,** Binned (10 kb) SNV counts of chromosome 1. Numbers 1-4 indicate mutation hotspots, **c,** Consensus ChromHMM states and rainfall plots at mutation hotspots, **d,** Pooled clustered variants from PCAWG Lymph-BHNL/CLL/NOS samples from TssA or TxFlnk (TS13-like), and all other epigenetic states (TS14-like). **e,** Size distribution of mutation clusters (consecutive clustered mutations), and the distribution of number of variants per mutation cluster in TS13 and TS14 high samples respectively.

While TS13 and TS14 are strongly correlated (*R*^2^=0.88, **Fig. S10**), the diverging localisation pattern and SNV spectrum, characterised by higher rates of C>K mutations in TS13, indicates that a related, but different mutational process drives TSS hypermutation, seemingly linked to AID. The differential mechanism behind TS13 also manifests as longer clusters (Median: 1,068 vs. 183bp), which contain more variants per cluster (Median: 8 vs. 3 mutations) in comparison to TS14 **(Fig. 7e).**

As a further distinction, the weakly clustered TLS signature TS14 can be found in more than 15 cancer types, suggesting a broad involvement of this mutagenic process in resolving endogenous and exogenous DNA alterations^28^. Polη has also been described to compete with lagging strand DNA synthesis^44^, which is further corroborated by the fact that TS14 displays a mild replicational strand bias (RSB=0.9; **Fig. 2b).** Interestingly, TS14 is found to be predominantly active in the regions without replication orientation (*a*_RS_=0.7), which are usually far from the origin of replication **(Fig. 2b).** Conversely, TS13 is mostly found in oriented, early replicating regions, but does not display a measurable replication strand bias **(Fig. 2b),** indicating different modes of activation.

Finally, a third mutational signature of somatic hypermutation, TS30, was found in lymphoid and other cancers of the HMF cohort. This signature displayed a large proportion of clustered mutations and an enrichment in early replicating regions similar to TS13, combined with an SNV spectrum that was closer to TS14 (Cosine distance *d=*0.13 vs. 0.25), suggesting that TS30 may represent a combination of TS13 and TS14.

## Discussion

We presented TensorSignatures, a novel framework for learning mutational signatures jointly from their mutation spectra and genomic properties to better understand the underlying mutational processes. We illustrated the capabilities of this algorithm by presenting a set of 20 mutational signatures extracted from 2,778 cancer genomes of the PCWAG consortium, and validated our analysis on additional 3,824 metastatic samples from the HMF cohort. The number of signatures was deliberately kept low for the signatures to be interpretable. The analysis demonstrated that the majority of mutational signatures comprised different variant types, and that no single mutational signature acted uniformly along the genome. Measuring how mutational spectra are influenced by their associated genomic features sheds light on the mechanisms underlying mutagenesis. A joint inference also helps to dissect mutational processes in situations where mutation spectra are very similar, such that genomic associations cannot be unambiguously attributed based on the mutation spectrum alone.

Studying the resulting signatures revealed that the SNV spectra of TS05 and TS06 show high similarity to signatures SBS7a and SBS7b of the COSMIC catalogue of mutational signatures. Due to the high similarity of the mutational spectra, it is difficult to unambiguously attribute individual mutations to these signatures and measure their genomic activity and transcriptional strand biases based on the mutation spectra alone. TensorSignature analysis reveals that the two processes are strongly differing with respect to their epigenetic context and transcriptional strand bias pointing towards differentially active GG-NER to be the underlying cause of the regional signature, which is confirmed by analysing cSCCs from GG-NER deficient XPC patients.

A similar change of the mutation spectrum was observed in Liver-HCC and other cancer types, reflected by the diverging activity of TS07 and TS08. The activity of TS08 is most prominent in highly transcribed genes, indicative of transcription-associated mutagenesis^12, 18^. TensorSignatures unifies the overarching mutational spectrum of this process and sheds light on its genomic determinants. Furthermore, its ability to detect mutational signatures in specific genomic regions also increases the sensitivity to detect signature activity, which may only contribute low levels of mutation at a genome wide scale. Here, we find TS08 also in Bladder-TCC, ColoRectal-AdenoCa, Lung-AdenoCa, Prostate-AdenoCa and Stomach-AdenoCa in addition to Billiary-AdenoCa, Head-SCC, and Liver-HCC, where it has been previously found^9^.

TensorSignatures’ capability to detect signatures with a confined regional context was also highlighted by detecting a highly localised signature associated with AID, TS13, which specifically manifests in and around transcription start sites in lymphoid neoplasms^7^. This signature has a base substitution spectrum similar to TS14 (SBS9), which does not display the tight localisation to TSS and is found in a range of cancer types, likely reflecting Polη-driven TLS during replication.

Inclusion of other mutation types led to the discovery of two APOBEC-associated signatures representative for mutagenesis during replication and at DSBRs, which differ with regard to their replicational strand bias and clustering propensity. Specifically, APOBEC-mediated mutagenesis at SVs lacks any preference for leading or lagging strand and is up to 80% clustered, suggesting that the formation of single stranded DNA during DSBR may trigger APOBEC activity. While an association of rearrangement events was reported earlier^1^, our study adds that DSBR- and replication-driven APOBEC mutations can be discerned by replication strand bias, clustering rate and size of clusters, indicating differential processivity of these two processes enabling different rates of mutation.

Validation of TensorSignature’s predictions was achieved by applying the algorithm to a second cohort of whole genomes from the Hartwig Medical Foundation. Reassuringly, the algorithm confirmed spectra and genomic properties from our primary analysis, thereby demonstrating the robustness of this approach, and the value of assessing additional genomic dimensions and other mutation types to gain more in depth insights to mutagenesis.

This analysis maps out the regional activity of mutational processes across the genome and pinpoints their various genomic determinants. Further improvements may include incorporation of more genomic features, potentially so in quantitative ways and ideally matched to the specific cell type. Currently TensorSignatures doesn’t model a preferred activity of a particular signature in a given tissue type and including such preference may help better ascertain the sets of signatures active in a particular genome. As mutational signature analysis is an essential element of the cancer genome analysis toolkit, TensorSignatures may help make the growing catalogues of mutational signatures more insightful by highlighting mutagenic mechanisms, or hypotheses thereof, to be investigated in greater depth.

## Methods

### Count tensor

#### Transcription

To assign single base substitutions to template and coding strand, we partitioned the genome by transcription directionality (trx(+)/trx(-)) using gencode V19 definitions. Nucleic acids can only be synthesized in 5’→3’ direction implying that template and coding strand of trx(-) genes are 5’→з’ and 3’ →5’ oriented, and vice versa for trx(+) genes. Since mutations are called on the + strand of the reference genome, and representing single base substitutions in a pyrimidine base context, we can unambiguously determine whether the pyrimidine of the mutated Watson-Crick base pair was on the coding or template strand. For example, a G>A substitution in a trx(-) gene corresponds to a coding strand C>T mutation, because the transcription directionality dictates that the mutated G sits on the template strand. Splitting all SNVs in this fashion requires us to introduce an additional dimension of size three (coding, template and unknown strand) to the count matrix (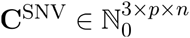 where *p*=96 and n is the number of samples).

#### Replication

To assign single base substitutions to leading and lagging strand, we leveraged Repli-seq data from the ENCODE consortium^45,46^, which map the sequences of nascent DNA replication strands throughout the whole genome during each cell cycle phase. Repli-seq profiles relate genomic coordinates to replication timing (early and late), where local maxima (peaks) and minima (valleys) correspond to replication initiation and termination zones. Regions between those peaks and valleys are characterized by steep slopes, whose sign (rep(+) or rep(-)) indicates whether the leading strand is replicated into the right (right-replicating) or left direction (left-replicating) when the DNA is viewed in standard orientation, respectively.

To partition the genome into non-overlapping right and left replicating regions, we computed the mean of slopes from Repli-seq profiles of five cell lines (GM12818, K564, Hela, Huvec and Hepgг) using finite differences. We marked regions with a plus (+) if the slope was positive (and therefore right-replicating) and with minus (-) if the slope was negative (and henceforth left-replicating). To confidently assign these states, we required that the absolute value of the mean of slopes was at least larger than two times its standard deviations, otherwise we assigned the unknown (*) state to the respective region. Using this convention, a C>A variant in a rep(+) region corresponds to a template C for leading strand DNA synthesis (and a template G for lagging strand). Subsequent assignment of single base substitutions to leading and lagging strand is analogous to the procedure we used for transcription strand assignment, and adds another dimension of size of three to the count tensor 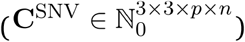.

#### Nucleosomal states

To assign single base substitutions to minor grooves facing away from and towards histones, and linker regions between nucleosomes, we used nucleosome dyad (midpoint) positions of human lymphoblastoid cell lines mapped in MNase cut efficiency experiments^23^. Although nucleosomal DNA binding is mediated by non-sequence specific minor groove-histone interactions, histone bound DNA features 5 bp AT-rich (minor in) followed by 5 bp GC-rich (minor out) DNA, as this composition bends the molecule favorably, while its characteristic structure may lead to differential susceptibility for mutational processes. Therefore, we partitioned nucleosomal DNA by first adding 7 bp to both sides of a dyad, and assigning the following 10 alternating 5 bp DNA stretches to minor out and minor in DNA, followed by a linker region with a maximum of 58 bp. Subsequent assignment of SNVs to these states adds another dimension of size four to the count tensor 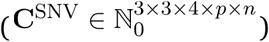.

#### Epigenetic states

To assign single base substitutions to different epigenetic environments, we used functional annotations from the 15-state ChromHMM model provided by the Roadmap epigenomics consortium^27^, which integrates multiple chromatin datasets such as ChlP-seq data of various histone modifications. To find state annotations that are robust across all cancer tissues, we defined an epigenetic consensus state by combining state annotations from 127 different Roadmap cell lines. Here, we required that at least 70 *%* of the cell lines agreed in the Chrom-HMM state to accept the state for a given genomic region. Partitioning SNVs by Chrom-HMM states adds another dimension of size 16 to the count tensor 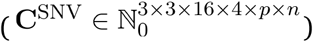.

#### Clustered mutations

To identify clustered single base substitutions, we used inter mutation distances (*Y_k_* in bp) between consecutive mutations on a chromosome as observations for a two state (*X_k_* = {clustered, unclustered}) hidden markov model with initial/transition distribution

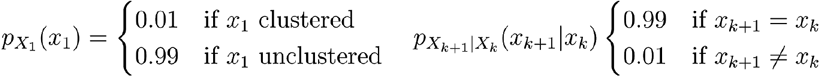

and observation distribution

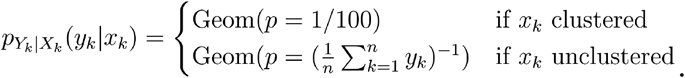

We then computed the maximum a posteriori (MAP) state using the Viterbi algorithm to assign to each mutation the state clustered or unclustered, respectively.

### Signature Tensor

In mutational signature analysis, NMF is used to decompose a catalogue of cancer genomes **C** to a set of mutational signatures **S** and their constituent activities or exposures E. This operation can be compactly expressed as

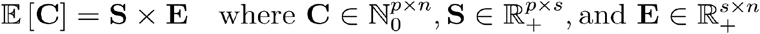

where *p* is the number of mutation types (usually *P* = 96), *n* the number of cancer genomes and *s* the number of mutational signatures.

Similarly, TensorSignatures identifies a low dimensional representation of a mutation count tensor, but decomposes it to mutational spectra for coding and template strand, leading and lagging strand, and signature specific multiplicative factors quantifying the propensities of mutational processes within specific genomic contexts. To enable strand specific extraction of mutational spectra requires to increase the dimensionality of the *p × s* sized signature matrix. To understand this, consider that two *p × s* matrices are at least needed to represent spectra for coding (C) and template (T) strand, suggesting a three dimensional (2 × *p × s*) signature representation. Our model, however, also considers replication, which adds another dimension of size two for leading (L) and lagging (G) strand, and thus we represent mutational spectra in the four dimensional core signature tensor **T**_0_ ∈ ℝ^2×2×*p×s*^

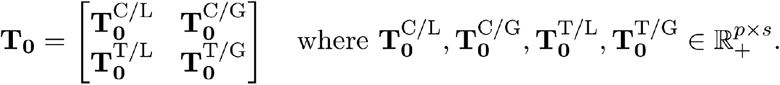

The mutation spectra 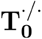 are normalised to 1 for each signature *s*, i.e., 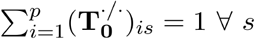. However, the mutation count tensor also contains mutations from genomic regions for which strand assignment was not applicable. To still use these data for the factorization, we map such counts to a linear combinations of **T**_0_’s sub matrices. This is enabled by *stacking* strand specific *p × s* matrices of the core signature tensor. For example, coding strand mutations for which replicational strand assignment was not applicable, are mapped to a linear combination of both coding strand specific sub matrices 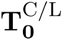 and 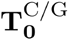. Stacking sub matrices of **T**_0_ results in 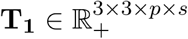

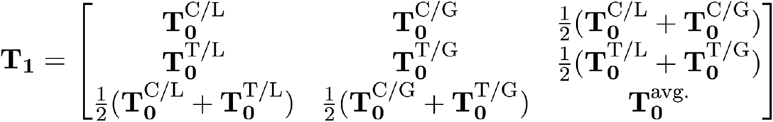

where 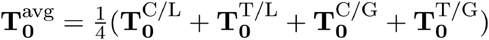.

#### Tensor factors

We use the term *tensor factor* for variables of the model that are factored into the signature tensor to quantify different genomic properties of a mutational signature. The key idea is to express a mutational process in terms of a product of strand specific spectra and a set of scalars, which modulate the magnitude of spectra dependent on the genomic state combination presented in the count tensor. However, to understand how tensor factors enter the factorization, it is necessary to introduce the concept of broadcasting, which is the process of making tensors with different shapes compatible for arithmetic operations.

It is important to realize that it is possible to increase the number of dimensions of a tensor by prepending their shapes with ones. For example, a three dimensional tensor **X** of shape 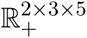 has 2 rows, 3 columns and a depth of 5. However, we could reshape **X** to 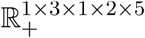, or 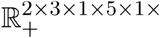, which would eventually change the order of values in the array, but not its content. These extra (empty) dimensions of **X** are called singletons or degenerates, and are required to make entities of different dimensionality compatible for arithmetic operations via *broadcasting.* To understand this, consider the following example

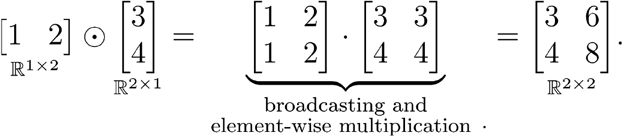

The ⊙ operator first copies the elements along their singleton axes such that the shape of both resulting arrays match, and then performs element-wise multiplication as indicated by the · symbol. This concept is similar to the tensor product ⊗ for vectors, but also applies to higher dimensional arrays, although this requires to define the shapes of all tensors carefully. For example if **F** ∈ ℝ^2×2^ and **H** ∈ ℝ^1×1×3^ then **F** ⊙ **H** is an invalid operation, however, if **G** ∈ ℝ^2×2×1^, then (**G ⊙ H**) ∈ ℝ^2×2×3^ ļ_s_ valid Also, note that such operations are not necessarily commutative.

#### Transcriptional and replicational strand biases

To quantify spectral asymmetries in context of transcription and replication, we introduce two vectors 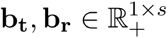 stack and reshape them such that the resulting bias tensor 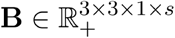

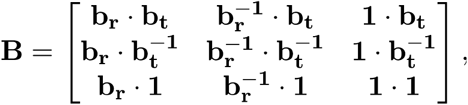

matches the shape of **T**_1_. Also, note that signs of b_t_ and b_r_ are chosen such that positive values correspond to a bias towards coding and leading strand, while negative values indicate shifts towards template and lagging strand

#### Signature activities in transcribed/untranscribed and early/late replicating regions

To assess the activity of mutational processes in transcribed versus untranscribed, and early versus late replicating regions, we introduce two additional scalars per signature represented in two vectors **a_t_** 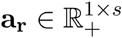. Both vectors are stacked and reshaped to match the shape of **T**_1_,

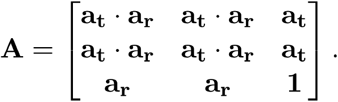

#### Mutational composition

To quantify the percentage of SNVs and other mutation types requires another 1 × *s* sized vector m, satisfying the constraint 0 ≤ **m**_*i*_ ≤ 1 for *i* – 1, …, *s*. In order to include **m** in the tensor factorization we reshape the vector to 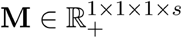, while (1 – **m**) is multiplied with the secondary signature matrix **S**.

We define the strand-specific signature tensor as

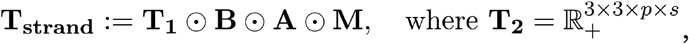

which therefore subsumes all parameters to describe a mutational process with regard to transcription and replication, and quantifies to what extent the signature is composed of SNVs. To understand this, consider the entry of the count tensor representative for coding strand mutations, e.g. 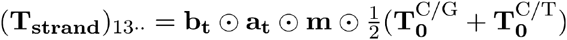; which explicitly states how the low dimensional tensor factors for transcription are broadcasted into the signature tensor.

#### Signature activities for nucleosomal, epigenetic and clustering states

The strand-specific signature tensor T_strand_ can be considered as the basic building block of the signature tensor, as we instantiate “copies” of T_strand_ by broadcasting scalar variables for each genomic state and signature along their respective dimensions. To understand this, recall that we, for example, split SNVs in *t* = 3 nucleosome states (minor in, minor out and linker regions). However, since SNVs may also fall into regions with no nucleosomal occupancy, we distributed mutations across *t* + 1 = 4 states in the corresponding dimension of the mutation count tensor. To fit parameters assessing the activity of each signature along these states, we initialize a matrix *k* ∈ ℝ^(*t*+1),*x*^, which can be considered as a composite of a 1 × *s* constant vector (**k**_1*i*_ = 1 for *i* = 1,…, *s*) and a *t ×s* matrix of state variables, allowing the model to adjust these parameters with respect to the first row, which corresponds to the non-nucleosomal mutations (baseline). To include these parameters in the factorization we first introduce a singleton dimensions in the strand specific signature tensor such that 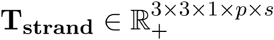, and reshape **k** to match the dimensionality of **T**_strand_,

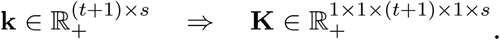

Both tensors have now the right shape such that element wise multiplication with broadcasting is valid

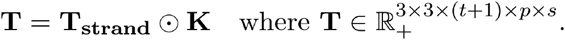

We proceed similarly for all remaining genomic properties such as activities along epigenetic domains, and clustering propensities. Generally, to assess *l* genomic properties, we first introduce *l* singleton dimensions to the strand-specific signature tensor **T**_strand_, instantiate *l* matrices 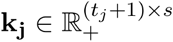 for *j* = 1, …, *l* each with *t_j_* states, reshape them appropriately to tensor factors **K**_j_, and broadcast them into the strand specific signature tensor **T**_2_. Here, we introduced new dimensions for epigenetic domains (epi), nucleosomal location (nuc) and clustering propensities (clu), and thus we reshaped the strand specific signature tensor to 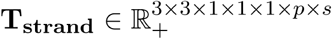, instantiated 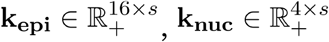 and 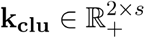 and computed

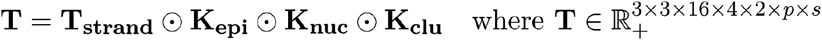

to obtain the final signature tensor **T**.

#### Model assumptions

The model assumes that the expected values of **C**^SNV^ and **C**^other^ are determined by the inner product of the signature tensor **T** (using the convention that × is taken over the last dimension of the array on its left – denoting each different signature – and the first dimension of the array on its right) and the exposure matrix **E** and similarly for the non-SNV signature matrix S and the same exposure matrix **E**

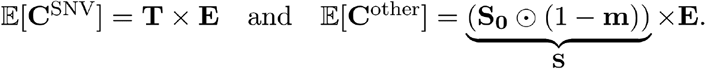

To prevent over segmentation and ensure a robust fit of signatures, we assume that the data follows a negative binomial distribution with mean **T × E** and **S × E**, and dispersion *τ*

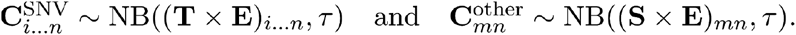

We use the Tensorflow framework to find the maximum likelihood estimates (MLE) 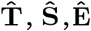 for **T, S** and **E** respectively using the parametrization defined in the previous section. We initialize the parameters of the model with values drawn from a truncated normal distribution and compute 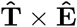 and 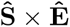 which are fed into the negative binomial likelihood function

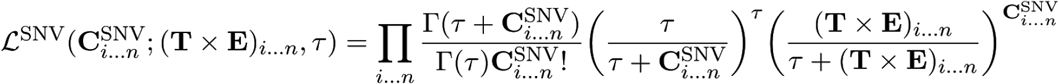

and

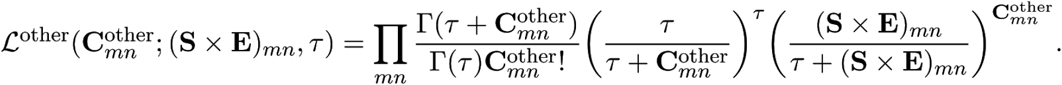

The total log likelihood log 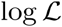 is then given by the sum of individual log likelihoods

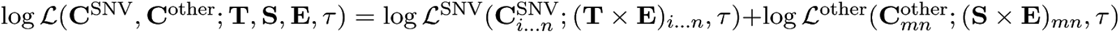

and thus the optimization problem boils down to maximize the total log likelihood (or equivalently to minimize the negative total log likelihood)

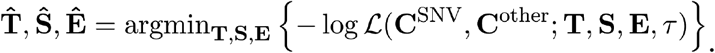

Moreover, inferring 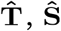, and 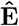 enables us to calculate log likelihood of the MLE

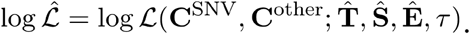

To calculate the value of each parameter in the model, we minimize the negative total log likelihood using an ADAM Grad optimizer with an exponentially decreasing learning rate of 0.1 and approximately 50,000 epochs.

### Model selection

To select the appropriate number of signatures for a model with dispersion rand dataset, we compute for each rank *s* the Bayesian Information Criterion (BIC)

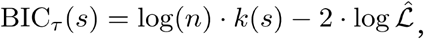

where *n* is the number of observations (total number of counts in **C**^SNV^ and **C**^other^), *k*(*s*) represents number of parameters in the model (which depends on the rank *s*), and log 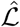 is the log-likelihood of the MLE. The BIC tries to find a trade-off between the log-likelihood and the number of parameters in the model; chosen is the rank which minimizes the BIC.

### Bootstrap Confidence Intervals

To compute bootstrap confidence intervals (CIs) for inferred parameters, we randomly select ¾ of the samples in the dataset, initialize the model with the MLE for 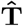 and 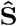 while randomly perturbing the 10% of their estimates, and subsequently refit 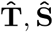 and 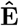 to the subset of samples. Initializing the parameters with the MLE results from computational constraints, as this step needs to be repeated for 300 – 500 times to obtain a representative distributions of the parameter space. Next, we match refitted signatures to the MLE reference by computing pairwise cosine distances, and accept bootstrap samples if the total variation distance between the bootstrap candidate and the reference is smaller than 0.2. Finally, we compute 5% and 95% percentiles on accepted bootstrap samples to indicate the CIs of our inference.

### XPC genomes

Somatic single nucleotide variants were called from .bam files were called as described in^47^. Subsequently these were aggregated into a mutation count tensor as described above.

### Comparing TensorSignatures to conventional NMF

#### Extracting signature properties using conventional NMF

We tried to quantify the genomic properties of mutational signatures using a less principled approach by simulating mutational signatures plus their genomic properties, sample exposures, and resulting mutation counts. To recover mutational signatures and corresponding sample exposures, we factorized (summed) mutation counts of simulated data using 96-trinucleotide channels only. To determine the strand biases and signatures activities across genomic states, we fixed the spectra of previously identified signatures, and refitted their exposures to the count matrix containing the mutations of a specific state only (eg. template strand mutations, TssA). To obtain a scalar parameter descriptive signature properties, we regressed state specific exposures to their respective baseline exposures (eg. exposures of template strand mutations against exposures of unassigned mutations) and compared obtained regression coefficients with the equivalent parameter of the tensor factorization and ground truth. To assess the error, we computed the vector 2-norm and cosine similarity for strand biases, genomic activities and exposures, and signature spectra respectively. We performed this experiment for datasets with sizes (100,1000,10000) and different numbers of mutations per sample (100,1000,10000). Note, in this approach it is not possible to recover signature activities in untranscribed/transcribed and early/late replicating regions (indicated as “Amplitudes” in the following plot).

To assess TensorSignatures’ ability of assigning mutations to their appropriate source signature, we designed a simulation experiment in which we used two very similar signatures (TS05 and TS06) to simulate a mutation count tensor. We then applied conventional NMF on the marginalized (summed) count tensor and determined the maximum a posteriori (MAP) signature for each trinucleotide context in each simulated sample as described in (Morganella et al. 2016).

#### TensorSignatures in comparison to previously applied methods

Absolute errors (vector 2-norm) of post-hoc assigned mutations increase as the number of mutations per sample get larger, while the predictions of the equivalent tensor factorisation become more accurate. This is to be expected as the post-hoc signature posterior probability is only conditioned on the mutation type and the sample exposure. Furthermore, shown results are likely to underestimate errors as our simulations/inferences were performed using only two signatures, and thus correct signature assignment is likely to happen by chance **(Fig. S2c).**

#### Stability of solutions

Another challenge in mutational signature analysis is the problem of unambiguously associating other variant types to their respective mutational processes. Common practice is to perform independent NMFs on each variant type, and to subsequently match subtype specific signatures to their SNV correlate by assessing exposures. In contrast, TensorSignatures decomposes SNV and other mutation type counts simultaneously, thus circumventing the problem of post-hoc associating different mutation types, and delivering a more robust signature inference by pooling evidence from the entire mutational imprint.

To illustrate this, we ran independent NMFs on SNV and other mutation count matrices of the PCAWG dataset. To match resulting mutational spectra, we computed the correlation coefficients of their exposures and paired highest correlating signatures. We repeated these steps 50 times to obtain a set of 50 initializations of paired mutational signatures (SNV + other mutation types), and compared the stability of these solutions with TensorSignature decompositions by computing the silhouette scores across several ranks **(Fig. S2d).**

Our results indicate a higher stability of TensorSignatures solutions across all tested ranks implying that the tensor framework more consistently reproduced SNV and their accompanying other mutation type spectra.

## Acknowledgements

We thank Oriol Pich, Santiago Gonzalez and Nuria Lopez for help in providing the genome coordinates for nucleosomes positions and variant calls for XPC genomes. Also, we thank Nadezda Volkova and Jose Guillherme de Almeida for commenting on our manuscript. This publication and the underlying study have been made possible partly on the basis of the data that Hartwig Medical Foundation and the Center of Personalised Cancer Treatment (CPCT) have made available to the study.

## Author contributions

H.V. conducted all bioinformatic analyses and produced the figures. A.v.H. and E.C. curated HMF data and provided computing resources for HMF data analysis by H.V.. M.G. conceived and supervised the analysis and developed code for categorising mutations. H.V. and M.G. wrote the manuscript with input from A.v.H. and E.C.

## Data and software availability

The analysis presented here is based on data generated by the PCAWG consortium, available at http://dcc.icgc.org/pcawg and the Hartwig Medical Foundation, available at https://www.hartwigmedicalfoundation.nl/applying-for-data/. The analysis involving XPC genomes is based on data from https://www.ncbi.nlm.nih.gov/projects/gap/cgi-bin/study.cgi?study_id=phs00080.v1.p1. Annotations used to classify variants by transcription and replication directionality stem from https://www.gencodegenes.org/human/release_19.html. and https://www.encodeproject.org/search/?type=Experiment&assav_title=Repli-seq_respectively. ChromHMM and nucleosome annotations were downloaded from https://egg2.wustl.edu/roadmap/web_portal/chr_state_learning.html and http://eqtl.uchicago.edu/nucleosomes/mnase_seq.html. All analyses were performed using Python 3.6 or R 3.4.

TensorSignature source code is available at http://github.com/gerstung-lab/tensorsignatures and as a pypi package “tensorsignatures”. This repository contains code for data preprocessing, genomic annotation and signature discovery and fitting. TensorSignatures can also be run as TensorSignaturesOnline, a web application accessible under http://193.62.55.163/home. that enables users to analyse their VCF data by attributing variants to a set of predefined Tensorsignatures. For the purpose of review please use these login data: Username: *review,* Password: *test123*.

## Supplementary Figures

**Supplementary Figure 1: Simulation experiments, a,** Accuracy of signature inference with respect to the number of samples (*n*) and the number of mutations per sample (*m*) in the simulated dataset. Signature recognition is defined as 1 minus cosine distance of the inferred and true signature, **b,** Accuracy of exposure inference with respect to the number of samples (*n*) and the number of mutations per sample (*m*) in the simulated dataset, **c,** Accuracy of inferred transcriptional and replicational activities (*a*_0_) and strand biases (*b*_0_), and SNV composition (*m*_1_) with respect to the number of samples (*n*), and the number of mutations per sample (m) in the simulated dataset, **d,** Accuracy of inferred epigenetic (*k*_0_) and nucleosomal activities (*k*_1_, and clustering propensites (*k*_2_ with respect to the number of samples (*m*) and the number of mutations per sample (*m*) in the simulated dataset, **e,** Accuracy of signature recognition at different ranks with respect to sample size (*n*) and number of mutations (*m*). **f,** Model selection via BIC (true rank 10).

**Supplementary Figure 2:** Upper panel: Schematic depiction of a simulation approach which recovers signature activities similar to TensorSignatures by regressing their prevalence in different genomic dimensions post-hoc. Lower two panels depict relative and absolute errors respectively.

**Supplementary Figure 3:** Upper panel: Schematic depiction of the simulation approach which uses a MAP estimation to assign mutations to their respective mutational signature as described by Morganella et al. Lower panel: Per sample absolute errors of the tensor factorisation and the aforementioned strategy.

**Supplementary Figure 4:** Left panel: Matching SNV and other mutation type spectra (from independent NMFs) by correlating their exposures. Right panel: Signature stability across several ranks using TensorSignatures and the signature matching approach.

**Supplementary Figure** 5: Model selection in the PCAWG dataset (chosen number of signatures 20 with a size *τ* of 50).

**Supplementary Figure 6: a,** Model selection in the HMF dataset (chosen number of signatures 27 with a size *τ* of 30) b, Squared errors of tensor factors from the PCAWG discovery and HMF validation analysis, **c,** C>T mutation type probabilities of TS22 for coding and template strand DNA, and the MNV spectrum of TS23.

**Supplementary Figure** 7: **a,** Correlation of TS05 and TS06; exposures in Skin-Melanoma samples, **b,** Heptanucleotide context normalized C>T mutation counts in active and quiescent genomic regions, **c,** Pooled C>T variants from cSCC XPC^-/-^ and cSCC XPC^wt^ genomes from active and quiescent regions respectively. Transcriptional strand bias and C[C>T]/T[C>T] spectral shift in GG-NER deficient XPC^wt^ cSCC genomes.

**Supplementary Figure 8: a,** Correlation of predicted TS07 and TS08 mutation counts in Liver-HCC samples, **b,** T>C mutation counts from active genomic regions in samples with high TS08 activity (other than Liver-HCC).

**Supplementary Figure 9:** Pancancer-wide pooled C>G and C>T clustered variants proximal and distal to SVs.

**Supplementary Figure 10:** Correlation of TS13 and TS14 exposures in lymphoid cancers (Lymph-BNHL/CLL/NOS).

## Notes

https://github.com/gerstung-lab/tensorsignatures

### Competing Interest Statement

The authors have declared no competing interest.

### Summary of Updates

This version of the manuscript has been revised to include a validation analysis performed on a second independent whole genome cancer cohort of the Hartwig Medical Foundation. Figure 3, 4, 5 and 6 revised; Figure 7 added; Author list updated; Supplemental files updated.

## References

1. Nik-Zainal, S., Alexandrov, L. B., Wedge, D. C., Van Loo, P., Greenman, C. D., Raine, K., Jones, D., Hinton, J., Marshall, J., Stebbings, L. A., Menzies, A., Martin, S., Leung, K., Chen, L., Leroy, C., Ramakrishna, M., Rance, R., Lau, K. W., Mudie, L. J., Varela, I., McBride, D. J., Bignell, G. R., Cooke, S. L., Shlien, A., Gamble, J., Whitmore, I., Maddison, M., Tarpey, P. S., Davies, H. R., Papaemmanuil, E., Stephens, P. J., McLaren, S., Butler, A. P., Teague, J. W., Jönsson, G., Garber, J. E., Silver, D., Miron, P., Fatima, A., Boyault, S., Langerød, A., Tutt, A., Martens, J. W. M., Aparicio, S. A. J. R., Borg, A., Salomon, A. V., Thomas, G., Børresen-Dale, A.-L., Richardson, A. L., Neuberger, M. S., Futreal, P. A., Campbell, P. J., Stratton, M. R. & the Breast Cancer Working Group of the International Cancer Genome Consortium. Mutational Processes Molding the Genomes of 21 Breast Cancers. Cell 149, 979–993 (2012).

2. Alexandrov, L. B., Nik-Zainal, S., Wedge, D. C., Campbell, P. J. & Stratton, M. R. Deciphering signatures of mutational processes operative in human cancer. Cell Rep. 3, 246–259 (2013).

3. Alexandrov, L. B., Nik-Zainal, S., Wedge, D. C., Aparicio, S. A. J. R., Behjati, S., Biankin, A. V., Bignell, G. R., Bolli, N., Borg, A., Børresen-Dale, A.-L., Boyault, S., Burkhardt, B., Butler, A. P., Caldas, C., Davies, H. R., Desmedt, C., Eils, R., Eyfjörd, J. E., Foekens, J. A., Greaves, M., Hosoda, F., Hutter, B., Ilicic, T., Imbeaud, S., Imielinski, M., Jäger, N., Jones, D. T. W., Jones, D., Knappskog, S., Kool, M., Lakhani, S. R., López-Otín, C., Martin, S., Munshi, N. C., Nakamura, H., Northcott, P. A., Pajic, M., Papaemmanuil, E., Paradiso, A., Pearson, J. V., Puente, X. S., Raine, K., Ramakrishna, M., Richardson, A. L., Richter, J., Rosenstiel, P., Schlesner, M., Schumacher, T. N., Span, P. N., Teague, J. W., Totoki, Y., Tutt, A. N. J., Valdés-Mas, R., van Buuren, M. M., van‘t Veer, L., Vincent-Salomon, A., Waddell, N., Yates, L. R., Australian Pancreatic Cancer Genome Initiative, ICGC Breast Cancer Consortium, ICGC MMML-Seq Consortium, ICGC PedBrain, Zucman-Rossi, J., Futreal, P. A., McDermott, U., Lichter, P., Meyerson, M., Grimmond, S. M., Siebert, R., Campo, E., Shibata, T., Pfister, S. M., Campbell, P. J. & Stratton, M. R. Signatures of mutational processes in human cancer. Nature 500, 415–421 (2013).

4. Fischer, A., Illingworth, C. J. R., Campbell, P. J. & Mustonen, V. EMu: probabilistic inference of mutational processes and their localization in the cancer genome. Genome Biol. 14, R39 (2013).

5. Macintyre, G., Goranova, T. E., De Silva, D., Ennis, D., Piskorz, A. M., Eldridge, M., Sie, D., Lewsley, L.-A., Hanif, A., Wilson, C., Dowson, S., Glasspool, R. M., Lockley, M., Brockbank, E., Montes, A., Walther, A., Sundar, S., Edmondson, R., Hall, G. D., Clamp, A., Gourley, C., Hall, M., Fotopoulou, C., Gabra, H., Paul, J., Supernat, A., Millan, D., Hoyle, A., Bryson, G., Nourse, C., Mincarelli, L., Sanchez, L. N., Ylstra, B., Jimenez-Linan, M., Moore, L., Hofmann, O., Markowetz, F., McNeish, I. A. & Brenton, J. D. Copy number signatures and mutational processes in ovarian carcinoma. Nat. Genet. 50, 1262–1270 (2018).

6. Funnell, T., Zhang, A. W., Grewal, D., McKinney, S., Bashashati, A., Wang, Y. K. & Shah, S. P. Integrated structural variation and point mutation signatures in cancer genomes using correlated topic models. PLoS Comput. Biol. 15, e1006799 (2019).

7. Kasar, S., Kim, J., Improgo, R., Tiao, G., Polak, P., Haradhvala, N., Lawrence, M. S., Kiezun, A., Fernandes, S. M., Bahl, S., Sougnez, C., Gabriel, S., Lander, E. S., Kim, H. T., Getz, G. & Brown, J. R. Whole-genome sequencing reveals activation-induced cytidine deaminase signatures during indolent chronic lymphocytic leukaemia evolution. Nat. Commun. 6, 8866 (2015).

8. Kim, J., Mouw, K. W., Polak, P., Braunstein, L. Z., Kamburov, A., Kwiatkowski, D. J., Rosenberg, J. E., Van Allen, E. M., D‘Andrea, A. & Getz, G. Somatic ERCC2 mutations are associated with a distinct genomic signature in urothelial tumors. Nat. Genet. 48, 600–606 (2016).

9. Alexandrov, L., Kim, J., Haradhvala, N. J., Huang, M. N., Ng, A. W. T., Boot, A., Covington, K. R., Gordenin, D. A., Bergstrom, E., Lopez-Bigas, N., Klimczak, L. J., McPherson, J. R., Morganella, S., Sabarinathan, R., Wheeler, D. A., Mustonen, V., Getz, G., Rozen, S. G. & Stratton, M. R. The Repertoire of Mutational Signatures in Human Cancer. bioRxiv (2018). doi:10.1101/322859

10. Pleasance, E. D., Cheetham, R. K., Stephens, P. J., McBride, D. J., Humphray, S. J., Greenman, C. D., Varela, I., Lin, M.-L., Ordóñez, G. R., Bignell, G. R., Ye, K., Alipaz, J., Bauer, M. J., Beare, D., Butler, A., Carter, R. J., Chen, L., Cox, A. J., Edkins, S., Kokko-Gonzales, P. I., Gormley, N. A., Grocock, R. J., Haudenschild, C. D., Hims, M. M., James, T., Jia, M., Kingsbury, Z., Leroy, C., Marshall, J., Menzies, A., Mudie, L. J., Ning, Z., Royce, T., Schulz-Trieglaff, O. B., Spiridou, A., Stebbings, L. A., Szajkowski, L., Teague, J., Williamson, D., Chin, L., Ross, M. T., Campbell, P. J., Bentley, D. R., Futreal, P. A. & Stratton, M. R. A comprehensive catalogue of somatic mutations from a human cancer genome. Nature 463, 191–196 (2010).

11. Alexandrov, L. B., Ju, Y. S., Haase, K., Van Loo, P., Martincorena, I., Nik-Zainal, S., Totoki, Y., Fujimoto, A., Nakagawa, H., Shibata, T., Campbell, P. J., Vineis, P., Phillips, D. H. & Stratton, M. R. Mutational signatures associated with tobacco smoking in human cancer. Science 354, 618–622 (2016).

12. Imielinski, M., Guo, G. & Meyerson, M. Insertions and Deletions Target Lineage-Defining Genes in Human Cancers. Cell 168, 460–472.e14 (2017).

13. Meier, B., Volkova, N., Hong, Y., Schofield, P., Campbell, P. J., Gerstung, M. & Gartner, A. Mutational signatures of DNA mismatch repair deficiency in C. elegans and human cancers. bioRxiv (2018). doi:10.1101/149153

14. Cortes-Ciriano, I., Lee, S., Park, W.-Y., Kim, T.-M. & Park, P. J. A molecular portrait of microsatellite instability across multiple cancers. Nat. Commun. 8, 15180 (2017).

15. Kucab, J. E., Zou, X., Morganella, S., Joel, M., Nanda, A. S., Nagy, E., Gomez, C., Degasperi, A., Harris, R., Jackson, S. P., Arlt, V. M., Phillips, D. H. & Nik-Zainal, S. A Compendium of Mutational Signatures of Environmental Agents. Cell 177, 821–836.e16 (2019).

16. Nik-Zainal, S., Davies, H., Staaf, J., Ramakrishna, M., Glodzik, D., Zou, X., Martincorena, I., Alexandrov, L. B., Martin, S., Wedge, D. C., Van Loo, P., Ju, Y. S., Smid, M., Brinkman, A. B., Morganella, S., Aure, M. R., Lingjærde, O. C., Langerød, A., Ringnér, M., Ahn, S.-M., Boyault, S., Brock, J. E., Broeks, A., Butler, A., Desmedt, C., Dirix, L., Dronov, S., Fatima, A., Foekens, J. A., Gerstung, M., Hooijer, G. K. J., Jang, S. J., Jones, D. R., Kim, H.-Y., King, T. A., Krishnamurthy, S., Lee, H. J., Lee, J.-Y., Li, Y., McLaren, S., Menzies, A., Mustonen, V., O‘Meara, S., Pauporté, I., Pivot, X., Purdie, C. A., Raine, K., Ramakrishnan, K., Rodríguez-González, F. G., Romieu, G., Sieuwerts, A. M., Simpson, P. T., Shepherd, R., Stebbings, L., Stefansson, O. A., Teague, J., Tommasi, S., Treilleux, I., Van den Eynden, G. G., Vermeulen, P., Vincent-Salomon, A., Yates, L., Caldas, C., van‘t Veer, L., Tutt, A., Knappskog, S., Tan, B. K. T., Jonkers, J., Borg, Å., Ueno, N. T., Sotiriou, C., Viari, A., Futreal, P. A., Campbell, P. J., Span, P. N., Van Laere, S., Lakhani, S. R., Eyfjord, J. E., Thompson, A. M., Birney, E., Stunnenberg, H. G., van de Vijver, M. J., Martens, J. W. M., Børresen-Dale, A.-L., Richardson, A. L., Kong, G., Thomas, G. & Stratton, M. R. Landscape of somatic mutations in 560 breast cancer whole-genome sequences. Nature 534, 47–54 (2016).

17. Li, Y., Roberts, N., Weischenfeldt, J., Wala, J. A., Shapira, O., Schumacher, S., Khurana, E., Korbel, J. O., Imielinski, M., Beroukhim, R. & Campbell, P. Patterns of structural variation in human cancer. bioRxiv (2017). doi:10.1101/181339

18. Haradhvala, N. J., Polak, P., Stojanov, P., Covington, K. R., Shinbrot, E., Hess, J. M., Rheinbay, E., Kim, J., Maruvka, Y. E., Braunstein, L. Z., Kamburov, A., Hanawalt, P. C., Wheeler, D. A., Koren, A., Lawrence, M. S. & Getz, G. Mutational Strand Asymmetries in Cancer Genomes Reveal Mechanisms of DNA Damage and Repair. Cell 164, 538–549 (2016).

19. Morganella, S., Alexandrov, L. B., Glodzik, D., Zou, X., Davies, H., Staaf, J., Sieuwerts, A. M., Brinkman, A. B., Martin, S., Ramakrishna, M., Butler, A., Kim, H.-Y., Borg, Å., Sotiriou, C., Futreal, P. A., Campbell, P. J., Span, P. N., Van Laere, S., Lakhani, S. R., Eyfjord, J. E., Thompson, A. M., Stunnenberg, H. G., van de Vijver, M. J., Martens, J. W. M., Børresen-Dale, A.-L., Richardson, A. L., Kong, G., Thomas, G., Sale, J., Rada, C., Stratton, M. R., Birney, E. & Nik-Zainal, S. The topography of mutational processes in breast cancer genomes. Nat. Commun. 7, 11383 (2016).

20. Tomkova, M., Tomek, J., Kriaucionis, S. & Schuster-Böckler, B. Mutational signature distribution varies with DNA replication timing and strand asymmetry. Genome Biol. 19, 129 (2018).

21. Schuster-Böckler, B. & Lehner, B. Chromatin organization is a major influence on regional mutation rates in human cancer cells. Nature 488, 504–507 (2012).

22. Polak, P., Lawrence, M. S., Haugen, E., Stoletzki, N., Stojanov, P., Thurman, R. E., Garraway, L. A., Mirkin, S., Getz, G., Stamatoyannopoulos, J. A. & Sunyaev, S. R. Reduced local mutation density in regulatory DNA of cancer genomes is linked to DNA repair. Nat. Biotechnol. 32, 71–75 (2014).

23. Pich, O., Muiños, F., Sabarinathan, R., Reyes-Salazar, I., Gonzalez-Perez, A. & Lopez-Bigas, N. Somatic and Germline Mutation Periodicity Follow the Orientation of the DNA Minor Groove around Nucleosomes. Cell 175, 1074–1087.e18 (2018).

24. Campbell, P. J., Getz, G., Stuart, J. M., Korbel, J. O., Stein, L. D. & Pcawg. Pan-cancer analysis of whole genomes. bioRxiv (2017). doi:10.1101/162784

25. Abadi, M., Barham, P., Chen, J., Chen, Z., Davis, A., Dean, J., Devin, M., Ghemawat, S., Irving, G., Isard, M. & Others. Tensorflow: A system for large-scale machine learning. in 12th ${USENIX} Symposium on Operating Systems Design and Implementation ({OSDI}$ 16) 265–283 (usenix.org, 2016).

26. Letouzé, E., Shinde, J., Renault, V., Couchy, G., Blanc, J.-F., Tubacher, E., Bayard, Q., Bacq, D., Meyer, V., Semhoun, J., Bioulac-Sage, P., Prévôt, S., Azoulay, D., Paradis, V., Imbeaud, S., Deleuze, J.-F. & Zucman-Rossi, J. Mutational signatures reveal the dynamic interplay of risk factors and cellular processes during liver tumorigenesis. Nat. Commun. 8, 1315 (2017).

27. Roadmap Epigenomics Consortium, Kundaje, A., Meuleman, W., Ernst, J., Bilenky, M., Yen, A., Heravi-Moussavi, A., Kheradpour, P., Zhang, Z., Wang, J., Ziller, M. J., Amin, V., Whitaker, J. W., Schultz, M. D., Ward, L. D., Sarkar, A., Quon, G., Sandstrom, R. S., Eaton, M. L., Wu, Y.-C., Pfenning, A. R., Wang, X., Claussnitzer, M., Liu, Y., Coarfa, C., Harris, R. A., Shoresh, N., Epstein, C. B., Gjoneska, E., Leung, D., Xie, W., Hawkins, R. D., Lister, R., Hong, C., Gascard, P., Mungall, A. J., Moore, R., Chuah, E., Tam, A., Canfield, T. K., Hansen, R. S., Kaul, R., Sabo, P. J., Bansal, M. S., Carles, A., Dixon, J. R., Farh, K.-H., Feizi, S., Karlic, R., Kim, A.-R., Kulkarni, A., Li, D., Lowdon, R., Elliott, G., Mercer, T. R., Neph, S. J., Onuchic, V., Polak, P., Rajagopal, N., Ray, P., Sallari, R. C., Siebenthall, K. T., Sinnott-Armstrong, N. A., Stevens, M., Thurman, R. E., Wu, J., Zhang, B., Zhou, X., Beaudet, A. E., Boyer, L. A., De Jager, P. L., Farnham, P. J., Fisher, S. J., Haussler, D., Jones, S. J. M., Li, W., Marra, M. A., McManus, M. T., Sunyaev, S., Thomson, J. A., Tlsty, T. D., Tsai, L.-H., Wang, W., Waterland, R. A., Zhang, M. Q., Chadwick, L. H., Bernstein, B. E., Costello, J. F., Ecker, J. R., Hirst, M., Meissner, A., Milosavljevic, A., Ren, B., Stamatoyannopoulos, J. A., Wang, T. & Kellis, M. Integrative analysis of 111 reference human epigenomes. Nature 518, 317–330 (2015).

28. Supek, F. & Lehner, B. Clustered Mutation Signatures Reveal that Error-Prone DNA Repair Targets Mutations to Active Genes. Cell 170, 534–547.e23 (2017).

29. Tate, J. G., Bamford, S., Jubb, H. C., Sondka, Z., Beare, D. M., Bindal, N., Boutselakis, H., Cole, C. G., Creatore, C., Dawson, E., Fish, P., Harsha, B., Hathaway, C., Jupe, S. C., Kok, C. Y., Noble, K., Ponting, L., Ramshaw, C. C., Rye, C. E., Speedy, H. E., Stefancsik, R., Thompson, S. L., Wang, S., Ward, S., Campbell, P. J. & Forbes, S. A. COSMIC: the Catalogue Of Somatic Mutations In Cancer. Nucleic Acids Res. 47, D941–D947 (2019).

30. Shinbrot, E., Henninger, E. E., Weinhold, N., Covington, K. R., Göksenin, A. Y., Schultz, N., Chao, H., Doddapaneni, H., Muzny, D. M., Gibbs, R. A., Sander, C., Pursell, Z. F. & Wheeler, D. A. Exonuclease mutations in DNA polymerase epsilon reveal replication strand specific mutation patterns and human origins of replication. Genome Res. 24, 1740–1750 (2014).

31. Priestley, P., Baber, J., Lolkema, M. P., Steeghs, N., de Bruijn, E., Shale, C., Duyvesteyn, K., Haidari, S., van Hoeck, A., Onstenk, W., Roepman, P., Voda, M., Bloemendal, H. J., Tjan-Heijnen, V. C. G., van Herpen, C. M. L., Labots, M., Witteveen, P. O., Smit, E. F., Sleijfer, S., Voest, E. E. & Cuppen, E. Pan-cancer whole-genome analyses of metastatic solid tumours. Nature 575, 210–216 (2019).

32. Pich, O., Muiños, F., Lolkema, M. P., Steeghs, N., Gonzalez-Perez, A. & Lopez-Bigas, N. The mutational footprints of cancer therapies. Nat. Genet. 51, 1732–1740 (2019).

33. Christensen, S., vd Roest, B., Besselink, N. & Janssen, R. 5-Fluorouracil treatment induces characteristic T> G mutations in human cancer. bioRxiv (2019). at <https://www.biorxiv.org/content/10.1101/681262v1.abstract>

34. Pleguezuelos-Manzano, C., Puschhof, J., Rosendahl Huber, A., van Hoeck, A., Wood, H. M., Nomburg, J., Gurjao, C., Manders, F., Dalmasso, G., Stege, P. B., Paganelli, F. L., Geurts, M. H., Beumer, J., Mizutani, T., Miao, Y., van der Linden, R., van der Elst, S., Genomics England Research Consortium, Garcia, K. C., Top, J., Willems, R. J. L., Giannakis, M., Bonnet, R., Quirke, P., Meyerson, M., Cuppen, E., van Boxtel, R. & Clevers, H. Mutational signature in colorectal cancer caused by genotoxic pks+ E. coli. Nature 580, 269–273 (2020).

35. Hayward, N. K., Wilmott, J. S., Waddell, N., Johansson, P. A., Field, M. A., Nones, K., Patch, A.-M., Kakavand, H., Alexandrov, L. B., Burke, H., Jakrot, V., Kazakoff, S., Holmes, O., Leonard, C., Sabarinathan, R., Mularoni, L., Wood, S., Xu, Q., Waddell, N., Tembe, V., Pupo, G. M., De Paoli-Iseppi, R., Vilain, R. E., Shang, P., Lau, L. M. S., Dagg, R. A., Schramm, S.-J., Pritchard, A., Dutton-Regester, K., Newell, F., Fitzgerald, A., Shang, C. A., Grimmond, S. M., Pickett, H. A., Yang, J. Y., Stretch, J. R., Behren, A., Kefford, R. F., Hersey, P., Long, G. V., Cebon, J., Shackleton, M., Spillane, A. J., Saw, R. P. M., López-Bigas, N., Pearson, J. V., Thompson, J. F., Scolyer, R. A. & Mann, G. J. Whole-genome landscapes of major melanoma subtypes. Nature 545, 175–180 (2017).

36. Zheng, C. L., Wang, N. J., Chung, J., Moslehi, H., Sanborn, J. Z., Hur, J. S., Collisson, E. A., Vemula, S. S., Naujokas, A., Chiotti, K. E., Cheng, J. B., Fassihi, H., Blumberg, A. J., Bailey, C. V., Fudem, G. M., Mihm, F. G., Cunningham, B. B., Neuhaus, I. M., Liao, W., Oh, D. H., Cleaver, J. E., LeBoit, P. E., Costello, J. F., Lehmann, A. R., Gray, J. W., Spellman, P. T., Arron, S. T., Huh, N., Purdom, E. & Cho, R. J. Transcription restores DNA repair to heterochromatin, determining regional mutation rates in cancer genomes. Cell Rep. 9, 1228–1234 (2014).

37. Pleasance, E. D., Stephens, P. J., O‘Meara, S., McBride, D. J., Meynert, A., Jones, D., Lin, M.-L., Beare, D., Lau, K. W., Greenman, C., Varela, I., Nik-Zainal, S., Davies, H. R., Ordoñez, G. R., Mudie, L. J., Latimer, C., Edkins, S., Stebbings, L., Chen, L., Jia, M., Leroy, C., Marshall, J., Menzies, A., Butler, A., Teague, J. W., Mangion, J., Sun, Y. A., McLaughlin, S. F., Peckham, H. E., Tsung, E. F., Costa, G. L., Lee, C. C., Minna, J. D., Gazdar, A., Birney, E., Rhodes, M. D., McKernan, K. J., Stratton, M. R., Futreal, P. A. & Campbell, P. J. A small-cell lung cancer genome with complex signatures of tobacco exposure. Nature 463, 184–190 (2010).

38. Sankar, T. S., Wastuwidyaningtyas, B. D., Dong, Y., Lewis, S. A. & Wang, J. D. The nature of mutations induced by replication-transcription collisions. Nature 535, 178–181 (2016).

39. Chan, K., Roberts, S. A., Klimczak, L. J., Sterling, J. F., Saini, N., Malc, E. P., Kim, J., Kwiatkowski, D. J., Fargo, D. C., Mieczkowski, P. A., Getz, G. & Gordenin, D. A. An APOBEC3A hypermutation signature is distinguishable from the signature of background mutagenesis by APOBEC3B in human cancers. Nat. Genet. 47, 1067–1072 (2015).

40. Nik-Zainal, S., Wedge, D. C., Alexandrov, L. B., Petljak, M., Butler, A. P., Bolli, N., Davies, H. R., Knappskog, S., Martin, S., Papaemmanuil, E., Ramakrishna, M., Shlien, A., Simonic, I., Xue, Y., Tyler-Smith, C., Campbell, P. J. & Stratton, M. R. Association of a germline copy number polymorphism of APOBEC3A and APOBEC3B with burden of putative APOBEC-dependent mutations in breast cancer. Nat. Genet. 46, 487–491 (2014).

41. Petljak, M., Alexandrov, L. B., Brammeld, J. S., Price, S., Wedge, D. C., Grossmann, S., Dawson, K. J., Ju, Y. S., Iorio, F., Tubio, J. M. C., Koh, C. C., Georgakopoulos-Soares, I., Rodríguez-Martín, B., Otlu, B., O‘Meara, S., Butler, A. P., Menzies, A., Bhosle, S. G., Raine, K., Jones, D. R., Teague, J. W., Beal, K., Latimer, C., O‘Neill, L., Zamora, J., Anderson, E., Patel, N., Maddison, M., Ng, B. L., Graham, J., Garnett, M. J., McDermott, U., Nik-Zainal, S., Campbell, P. J. & Stratton, M. R. Characterizing Mutational Signatures in Human Cancer Cell Lines Reveals Episodic APOBEC Mutagenesis. Cell 176, 1282–1294.e20 (2019).

42. Muramatsu, M., Kinoshita, K., Fagarasan, S., Yamada, S., Shinkai, Y. & Honjo, T. Class switch recombination and hypermutation require activation-induced cytidine deaminase (AID), a potential RNA editing enzyme. Cell 102, 553–563 (2000).

43. Pham, P., Bransteitter, R., Petruska, J. & Goodman, M. F. Processive AID-catalysed cytosine deamination on single-stranded DNA simulates somatic hypermutation. Nature 424, 103–107 (2003).

44. Kreisel, K., Engqvist, M. K. M., Kalm, J., Thompson, L. J., Boström, M., Navarrete, C., McDonald, J. P., Larsson, E., Woodgate, R. & Clausen, A. R. DNA polymerase η contributes to genome-wide lagging strand synthesis. Nucleic Acids Res. 47, 2425–2435 (2019).

45. Hansen, R. S., Thomas, S., Sandstrom, R., Canfield, T. K., Thurman, R. E., Weaver, M., Dorschner, M. O., Gartler, S. M. & Stamatoyannopoulos, J. A. Sequencing newly replicated DNA reveals widespread plasticity in human replication timing. Proc. Natl. Acad. Sci. U. S. A. 107, 139–144 (2010).

46. Thurman, R. E., Day, N., Noble, W. S. & Stamatoyannopoulos, J. A. Identification of higher-order functional domains in the human ENCODE regions. Genome Res. 17, 917–927 (2007).

47. Sabarinathan, R., Mularoni, L., Deu-Pons, J., Gonzalez-Perez, A. & López-Bigas, N. Nucleotide excision repair is impaired by binding of transcription factors to DNA. Nature 532, 264–267 (2016).

