## Supplementary Figures for "Learning mutational signatures and their multidimensional genomic properties with TensorSignatures"

### Supplementary Figure 1

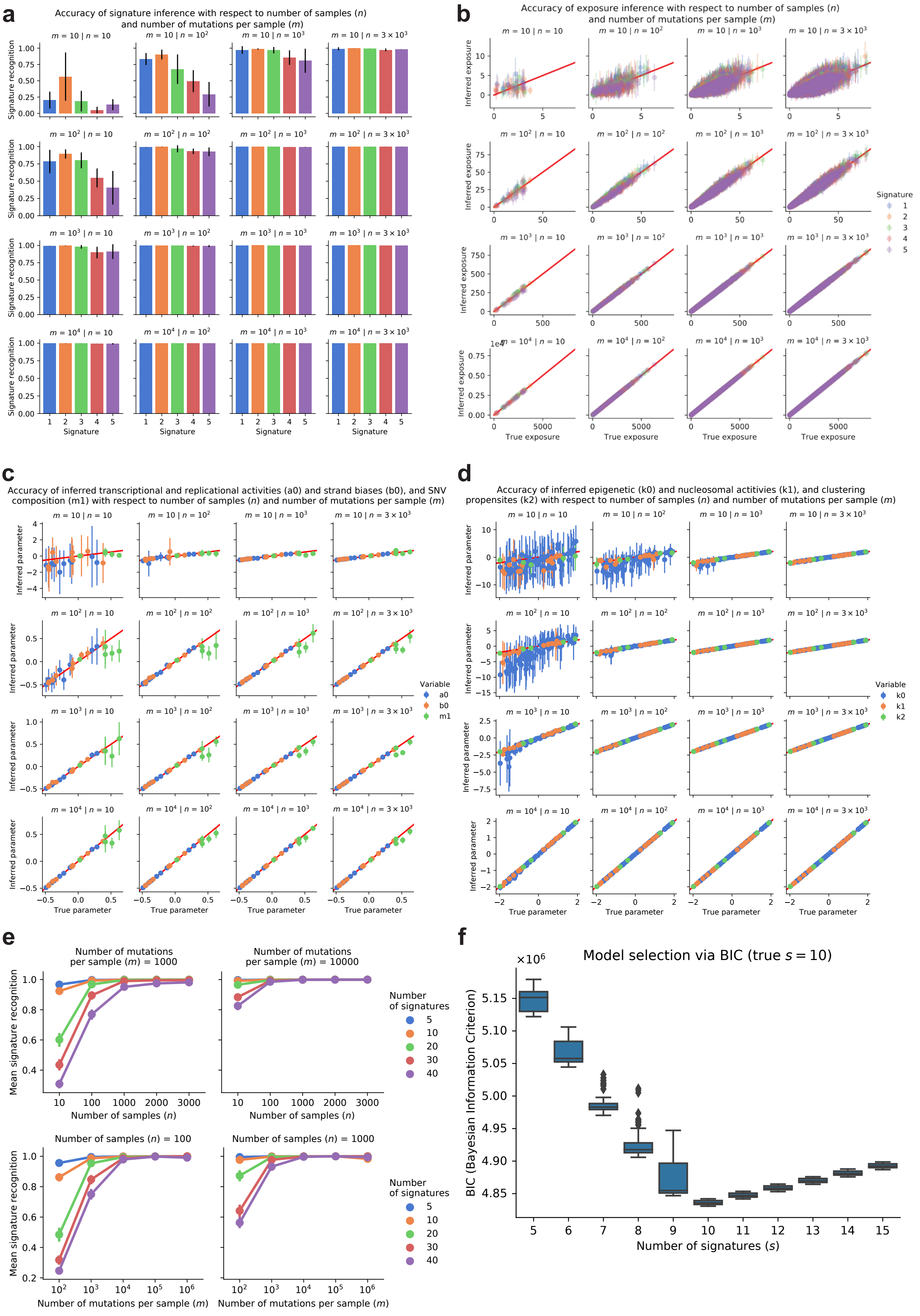

### Supplementary Figure 2

#### TensorSignatures vs. Post-hoc regression approach

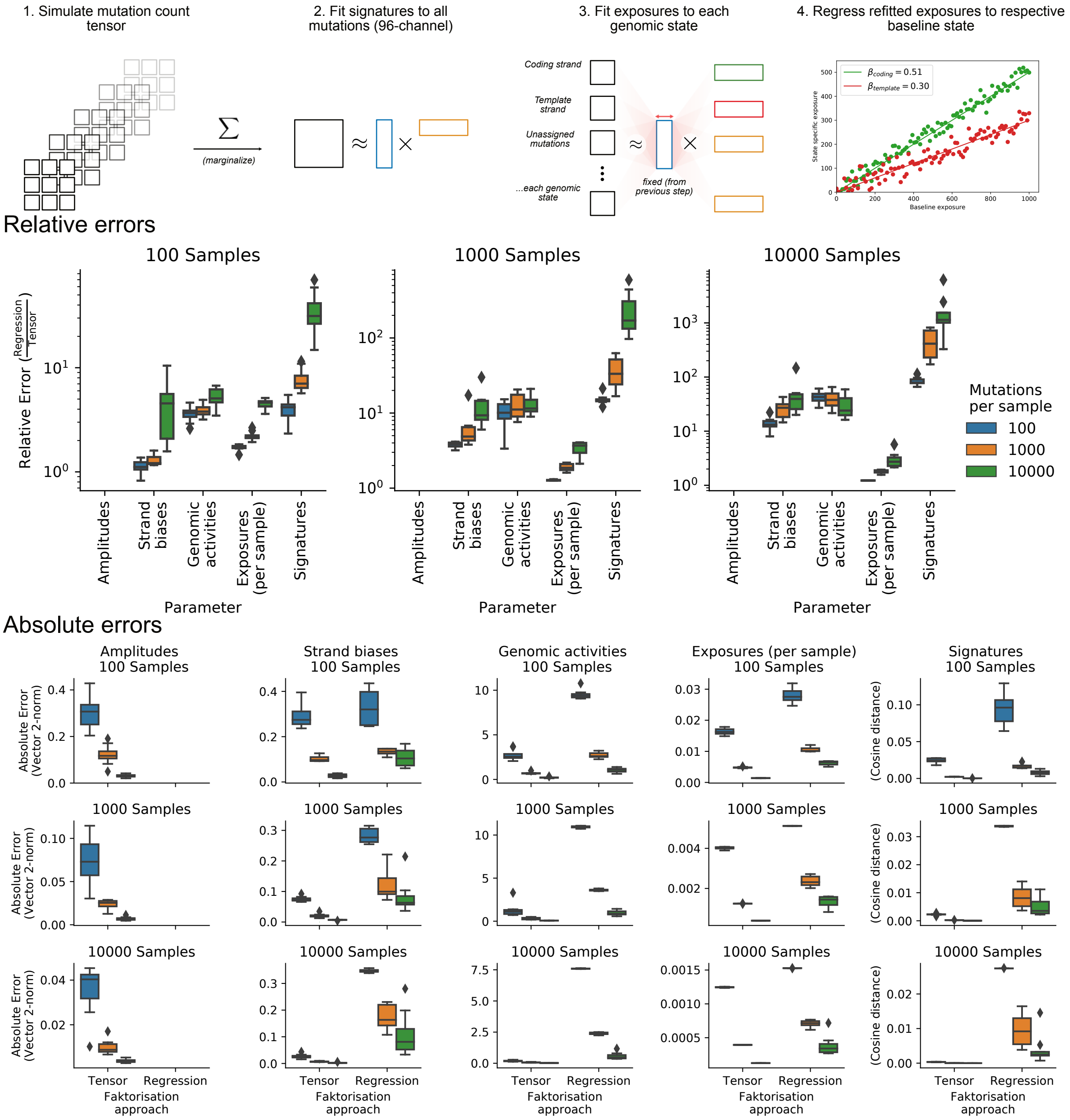

### Supplementary Figure 3

#### TensorSignatures vs. Post-hoc posterior calculations

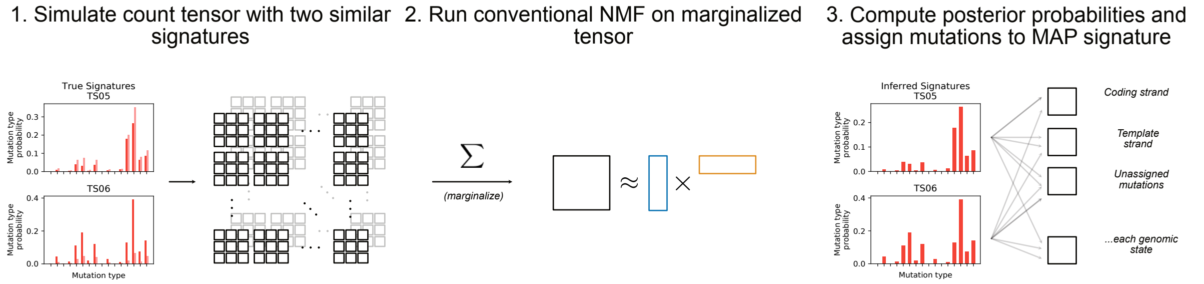


#### Per sample error

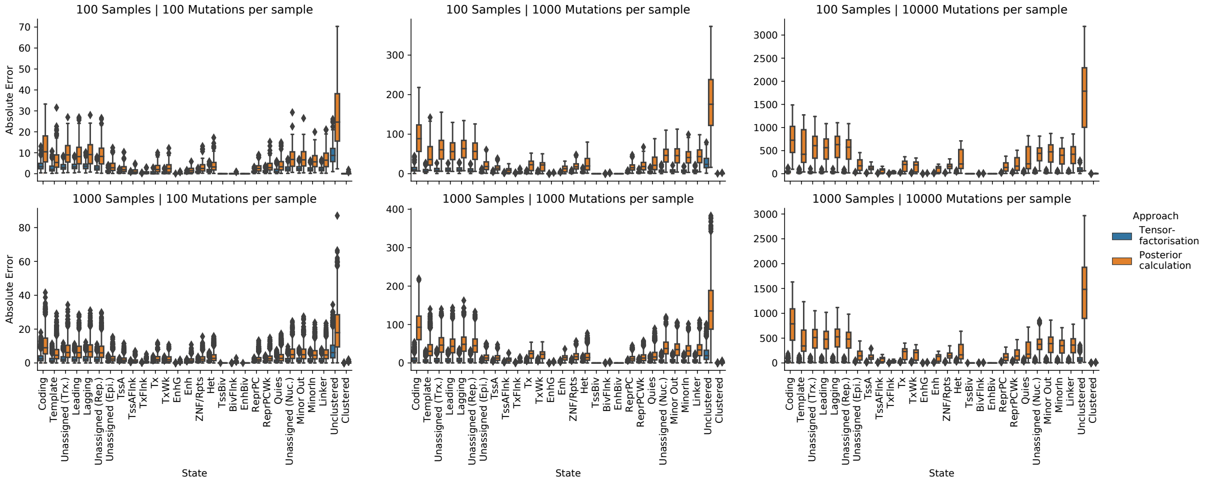

### Supplementary Figure 4

#### Stability of TensorSignature solution vs. concatenated independent NMFs

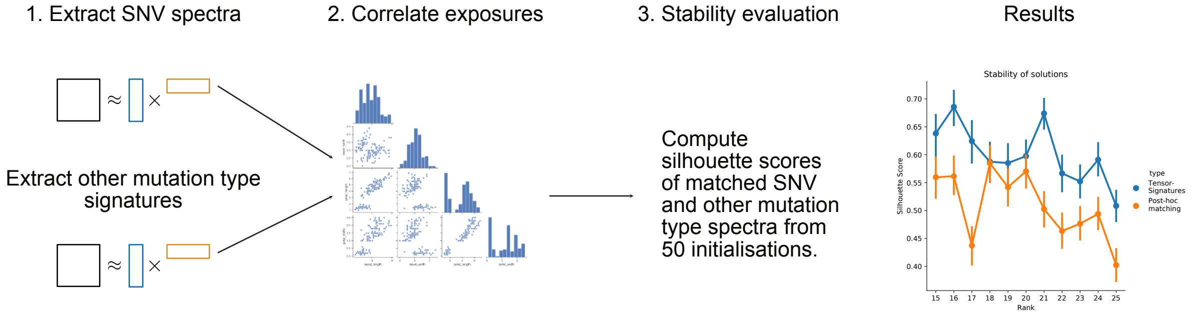

### Supplementary Figure 5

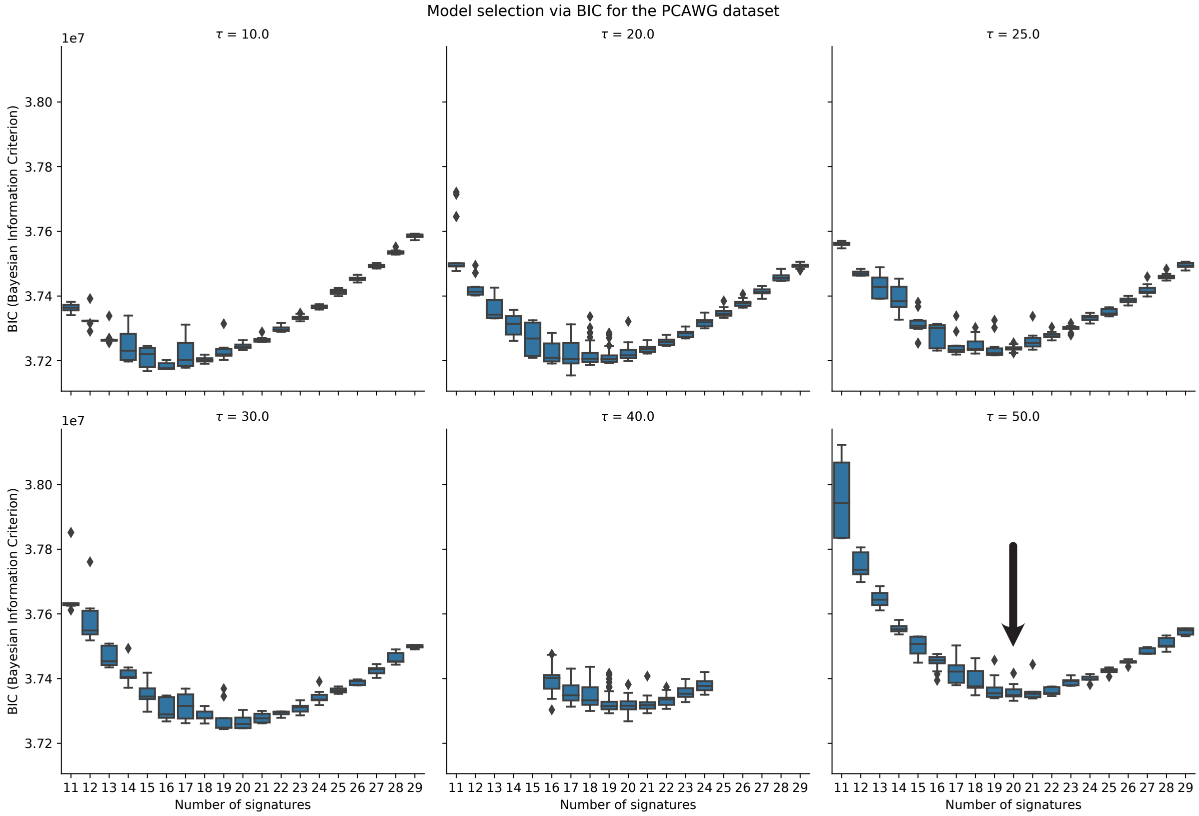

### Supplementary Figure 6

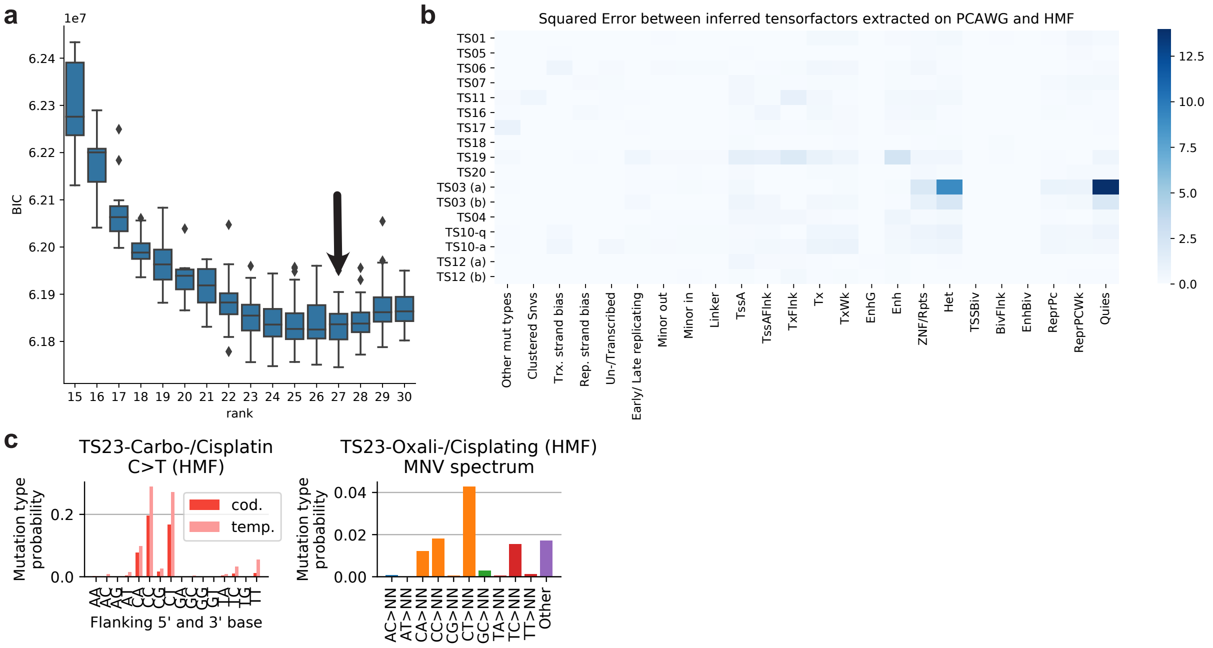

### Supplementary Figure 7

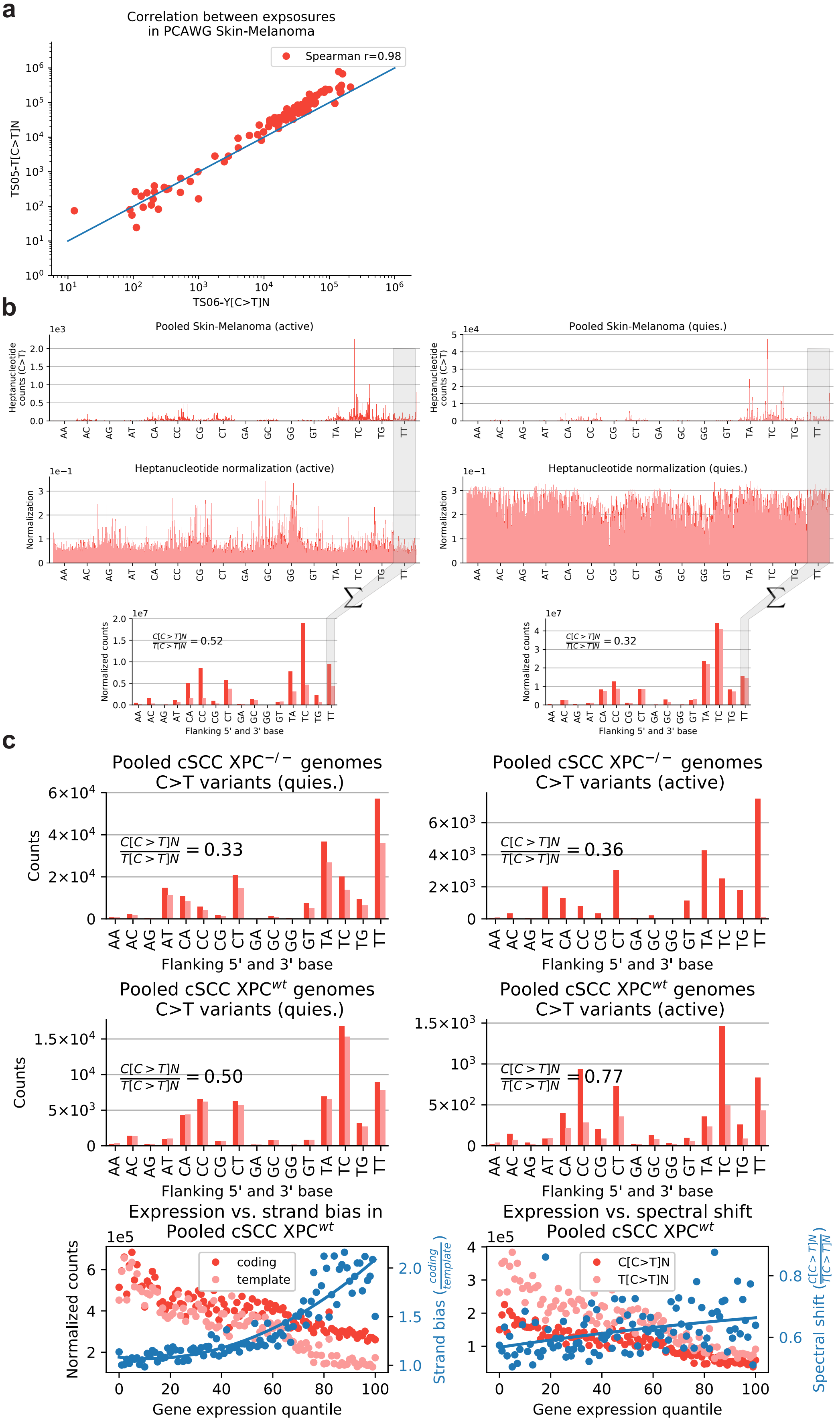

### Supplementary Figure 8

**a**

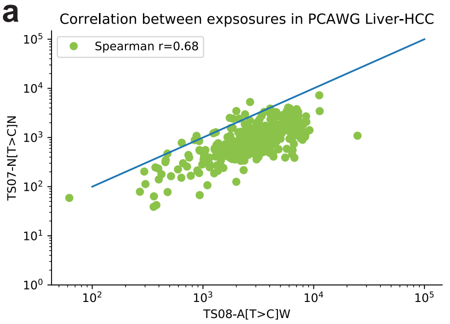


**b**

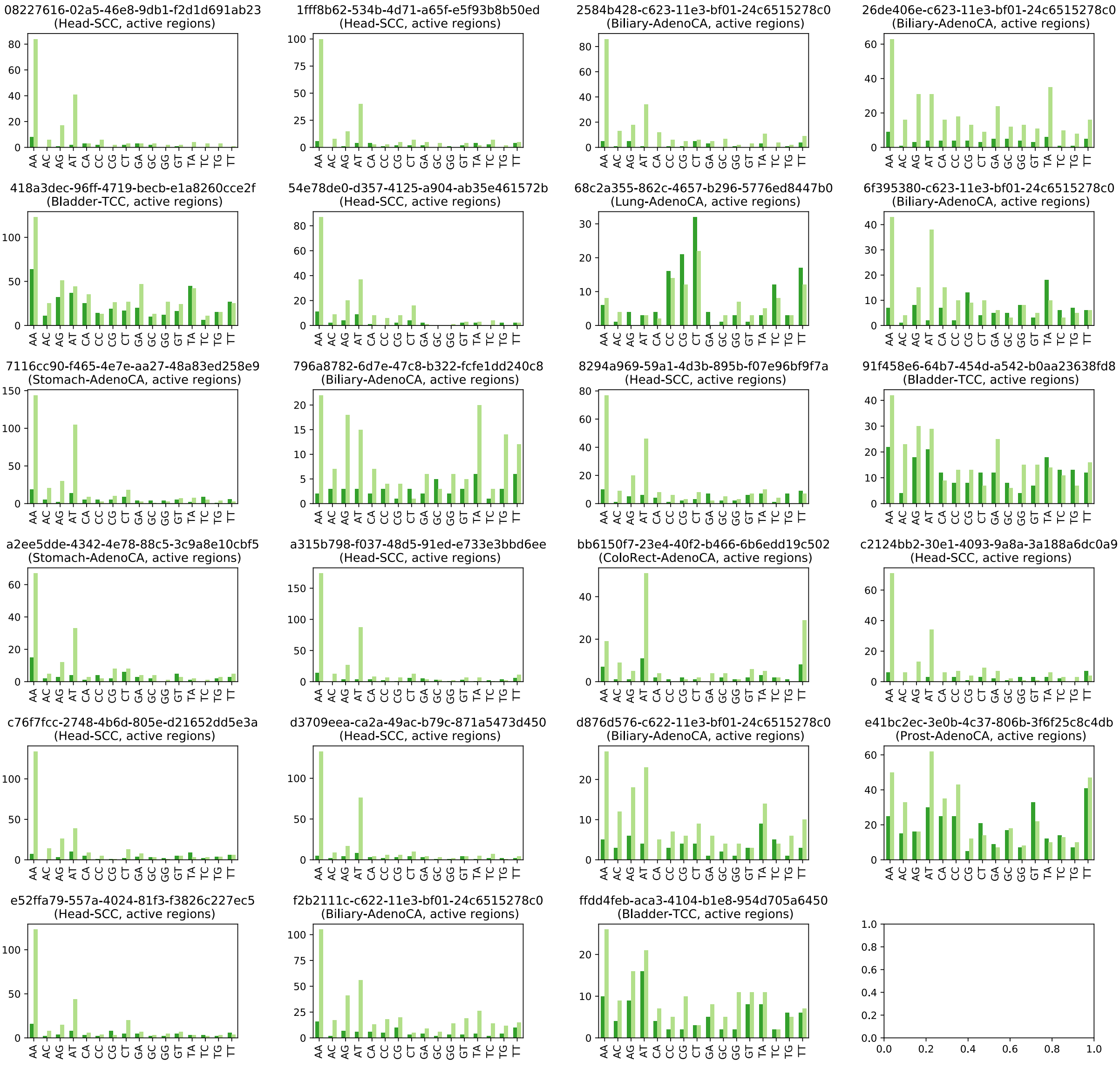

### Supplementary Figure 9

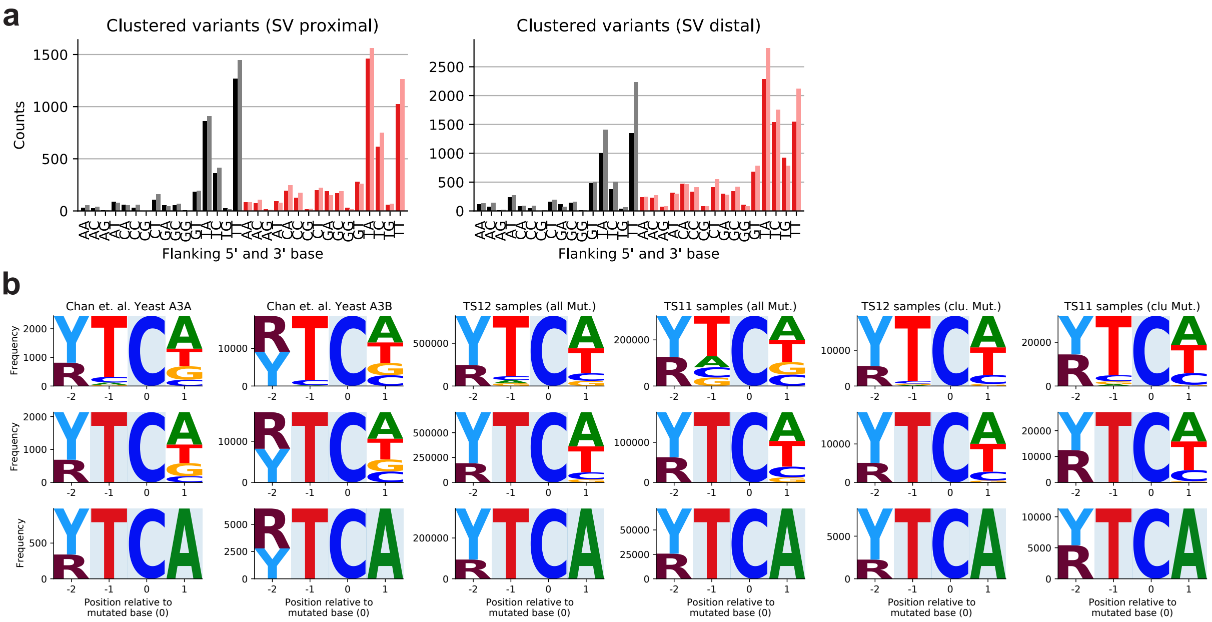

### Supplementary Figure 10

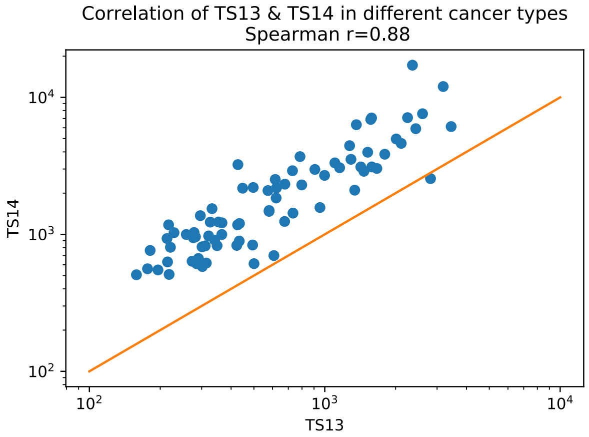
