## Supplementary Note for "Learning mutational signatures and their multidimensional genomic properties with TensorSignatures"

Harald Vöhringer and Moritz Gerstung
EMBL-EBI, University of Cambridge
Cambridgeshire, CB10 1SD

November 20, 2019

**Abstract**

Mutational signature analysis is an essential part of the cancer genome analysis toolkit. Conventionally, mutational signature analysis extracts patterns of different mutation types across many cancer genomes. Here we present TensorSignatures, an algorithm to learn mutational signatures jointly across all variant categories and their genomic context. The analysis of 2,778 cancer genomes of the PCAWG consortium shows that practically all signatures operate dynamically in response to various genomic and epigenomic states. The analysis pins differential spectra of UV mutagenesis found in active and inactive chromatin to global genome nucleotide excision repair. TensorSignatures accurately characterises transcription-associated mutagenesis, which is detected in 7 different cancer types. The analysis also unmasks replication and double strand break repair driven APOBEC mutagenesis, which manifests with differential numbers and length of mutation clusters indicating a differential processivity of the two triggers. As a fourth example, TensorSignatures detects a signature of somatic hypermutation generating highly clustered variants around the transcription start sites of active genes in lymphoid leukaemia, distinct from a more general and less clustered signature of Pol$\eta$-driven TLS found in a broad range of cancer types.

### 1 TS01-N[C>T]G (5meC>T)

#### 1.1 Overall SNV spectrum

Figure 1: Single base substitution spectrum.

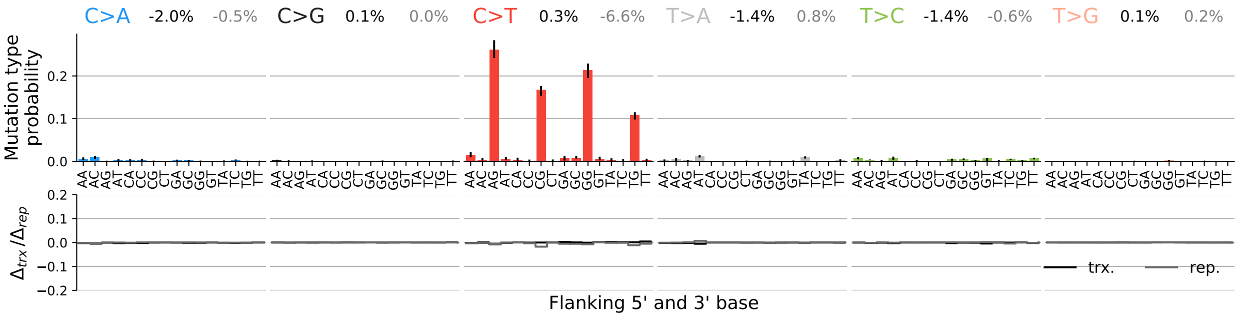

#### 1.2 SNV spectra for transcription and replication

Figure 2: Single base substitution spectra for template/coding and leading/lagging strand DNA.

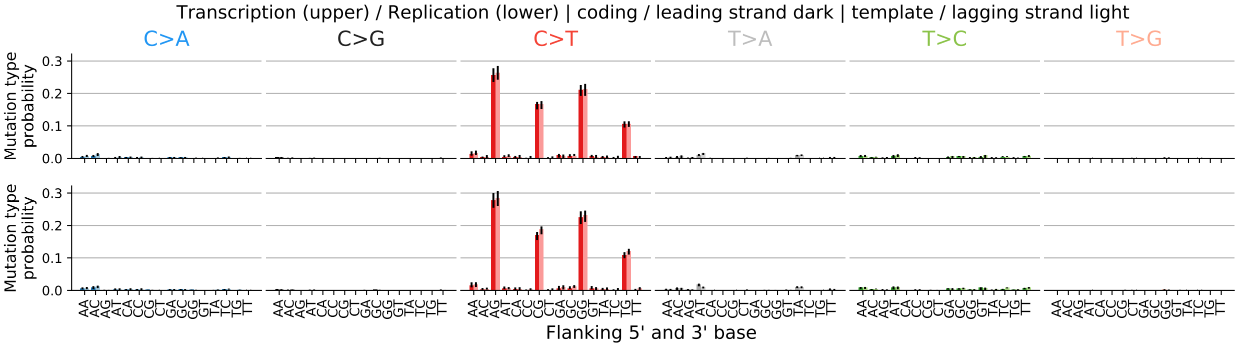

#### 1.3 Spectrum for other mutation types

Figure 3: Spectrum other mutation types.

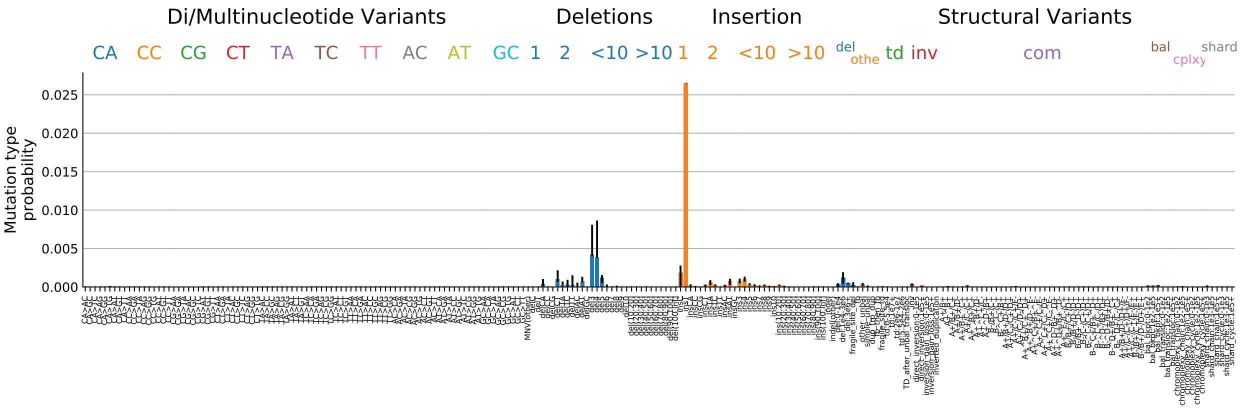

#### 1.4 Signature activity in different cancer types

Figure 4: Signature exposure in different cancer types.

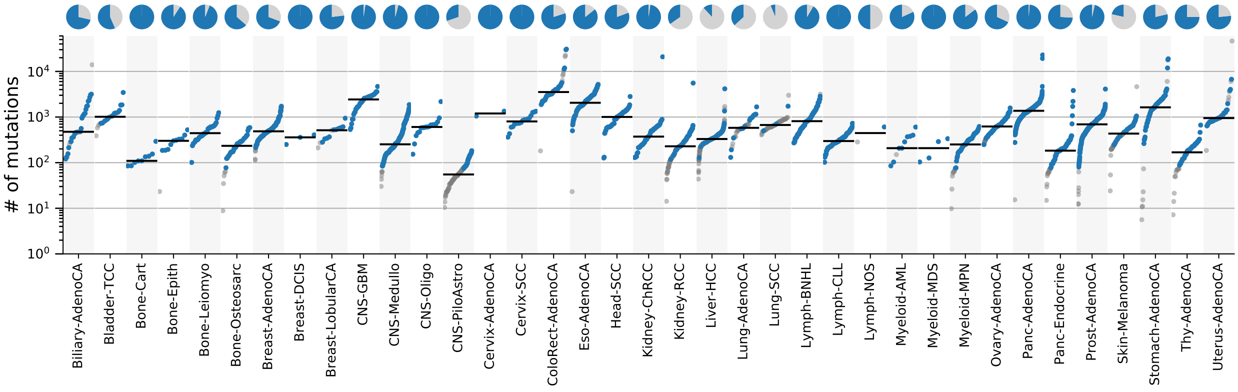

#### 1.5 Signature characterization

Figure 5: Signature specific tensor coefficients.

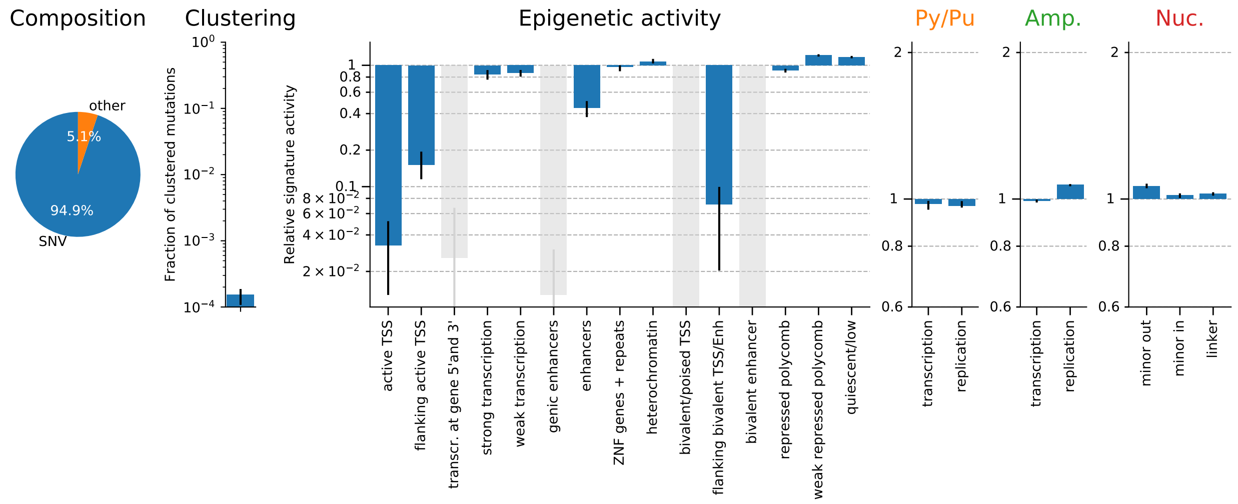

### 2 TS02-N[C>T]N (unknown)

#### 2.1 Overall SNV spectrum

Figure 6: Single base substitution spectrum.

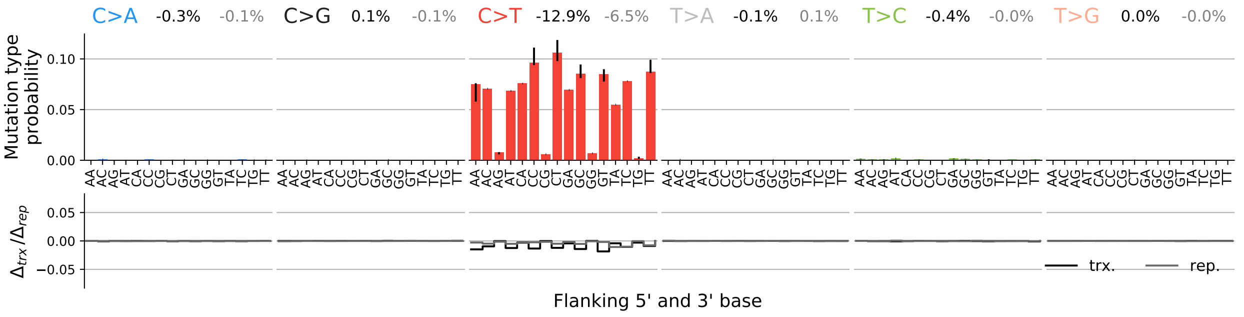

#### 2.2 SNV spectra for transcription and replication

Figure 7: Single base substitution spectra for template/coding and leading/lagging strand DNA.

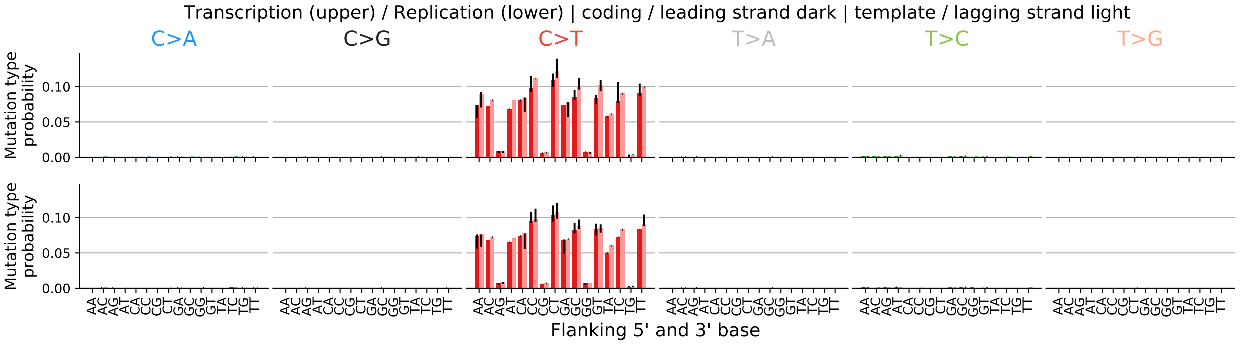

#### 2.3 Spectrum for other mutation types

Figure 8: Spectrum other mutation types.

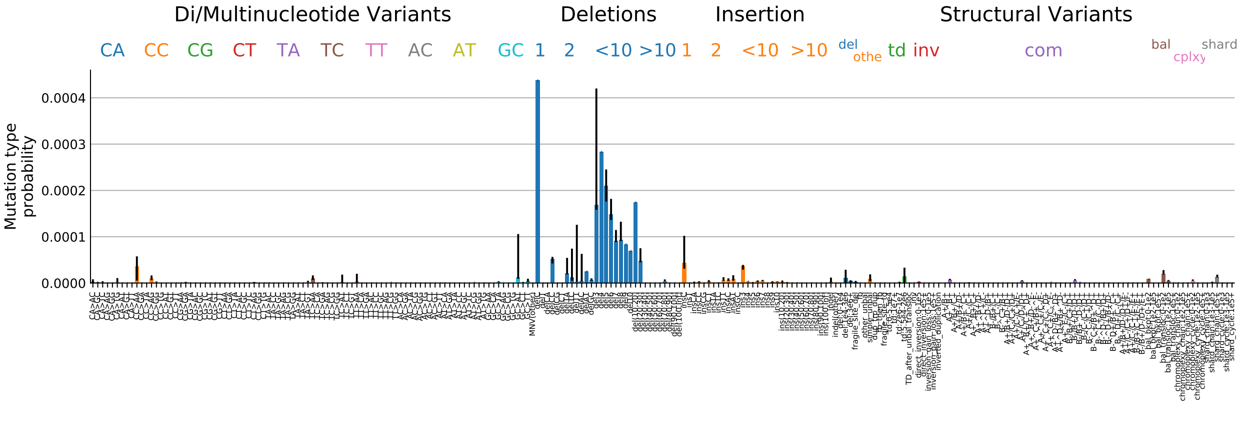

#### 2.4 Signature activity in different cancer types

Figure 9: Signature exposure in different cancer types.

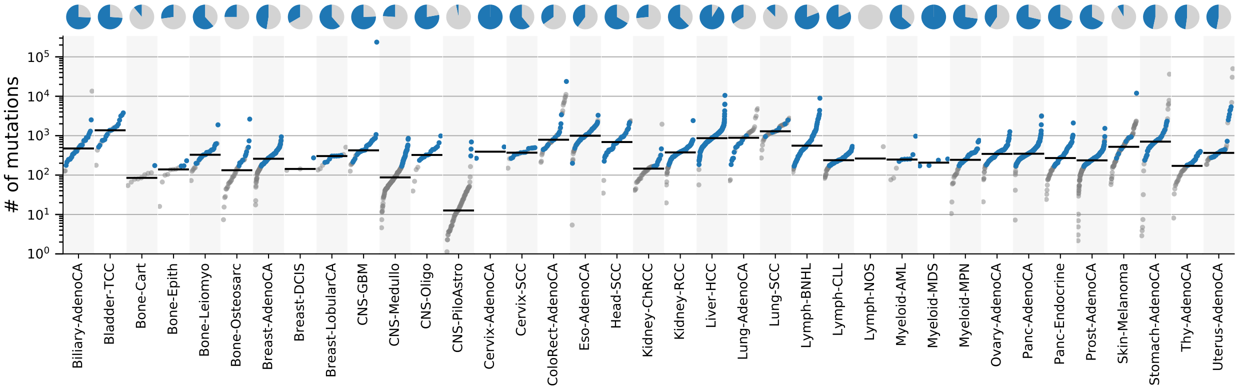

#### 2.5 Signature characterization

Figure 10: Signature specific tensor coefficients.

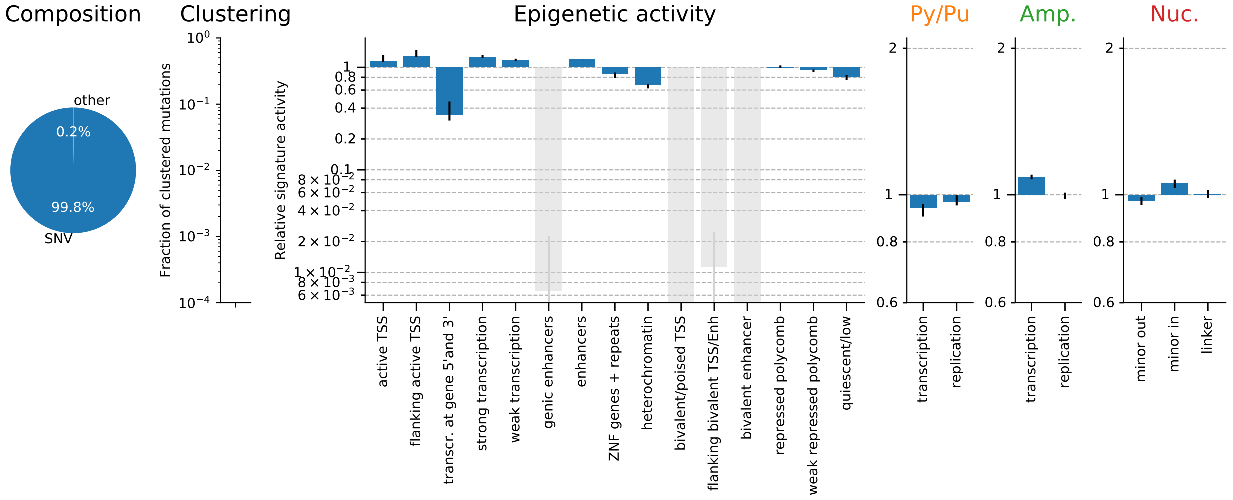

### 3 TS03-N[N>N]N-q (unknown/quiet)

#### 3.1 Overall SNV spectrum

Figure 11: Single base substitution spectrum.

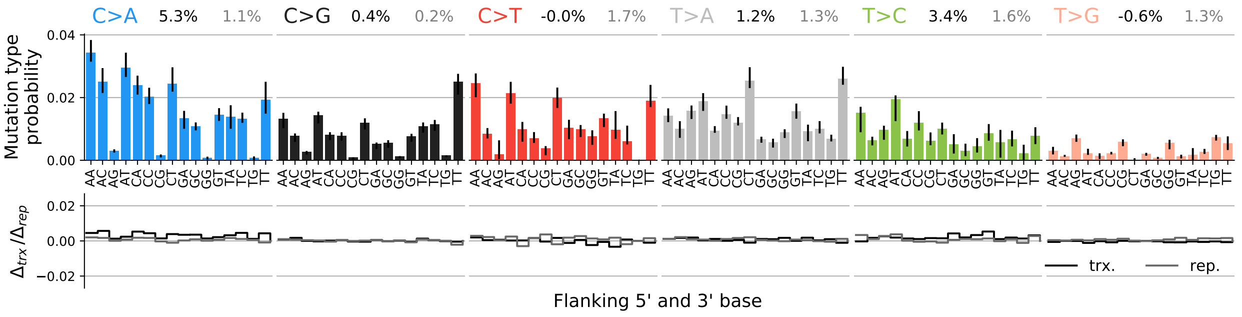

#### 3.2 SNV spectra for transcription and replication

Figure 12: Single base substitution spectra for template/coding and leading/lagging strand DNA.

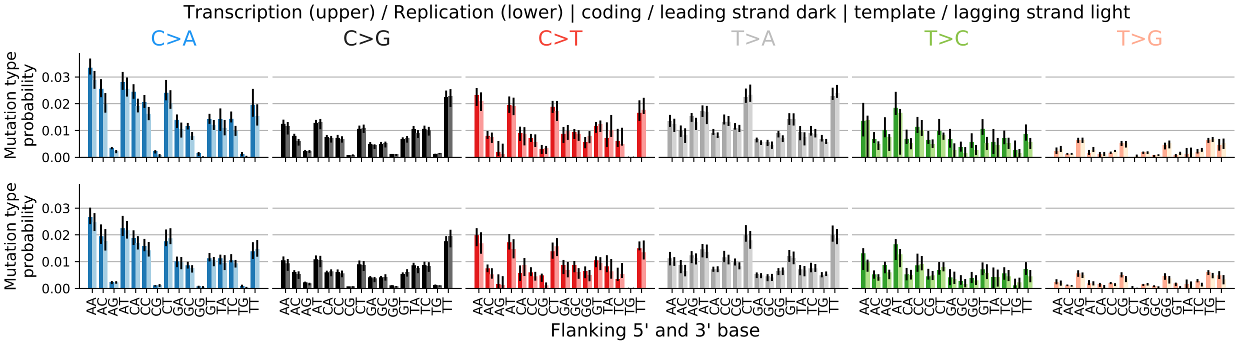

#### 3.3 Spectrum for other mutation types

Figure 13: Spectrum other mutation types.

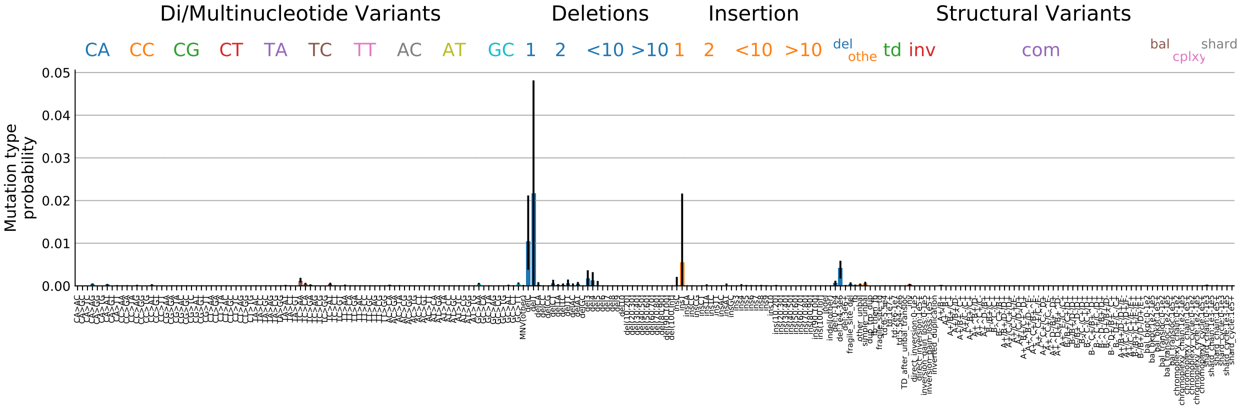

#### 3.4 Signature activity in different cancer types

Figure 14: Signature exposure in different cancer types.

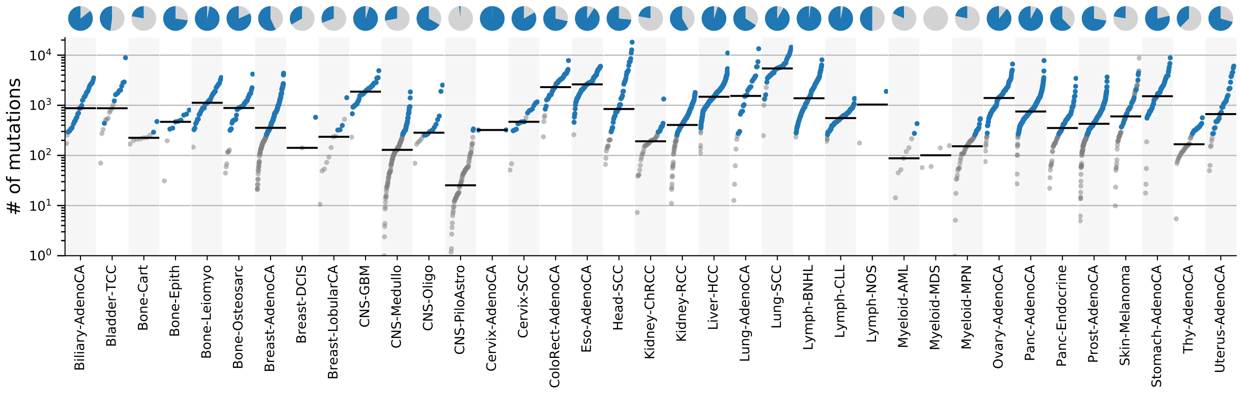

#### 3.5 Signature characterization

Figure 15: Signature specific tensor coefficients.

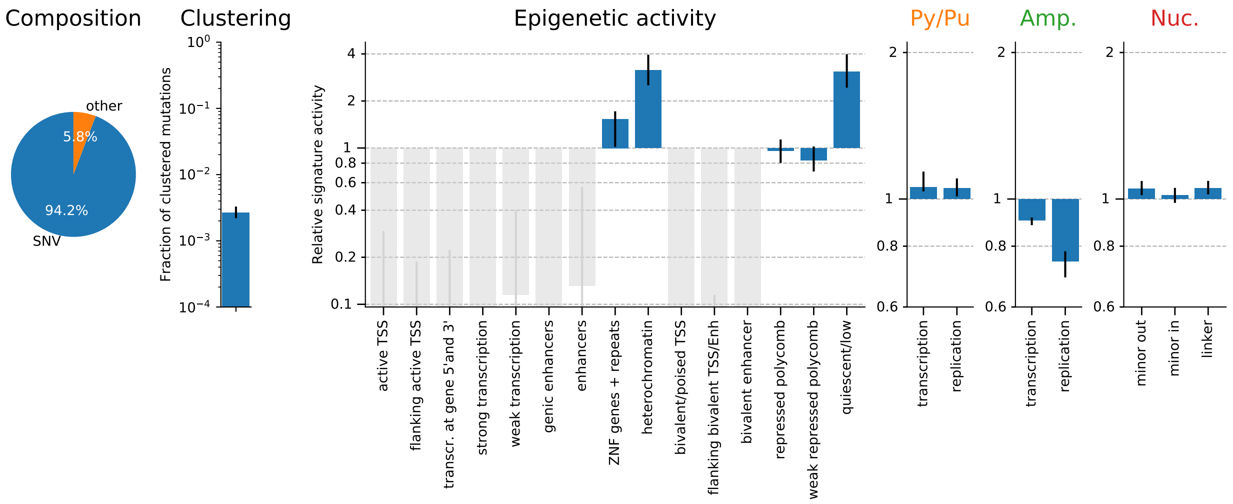

### 4 TS04-N[N>N]N (unknown/active)

#### 4.1 Overall SNV spectrum

Figure 16: Single base substitution spectrum.

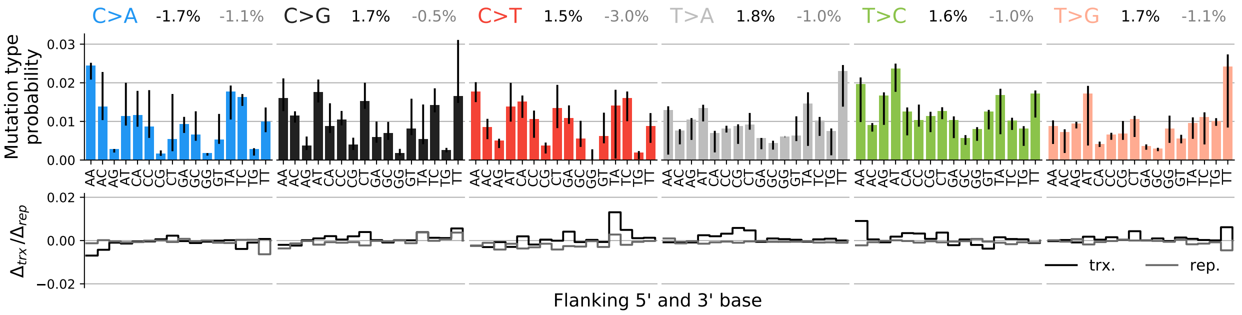

#### 4.2 SNV spectra for transcription and replication

Figure 17: Single base substitution spectra for template/coding and leading/lagging strand DNA.

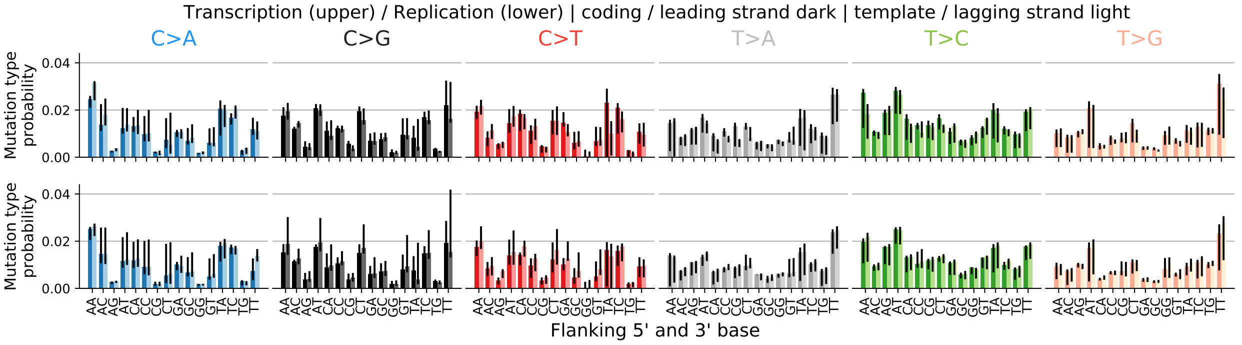

#### 4.3 Spectrum for other mutation types

Figure 18: Spectrum other mutation types.

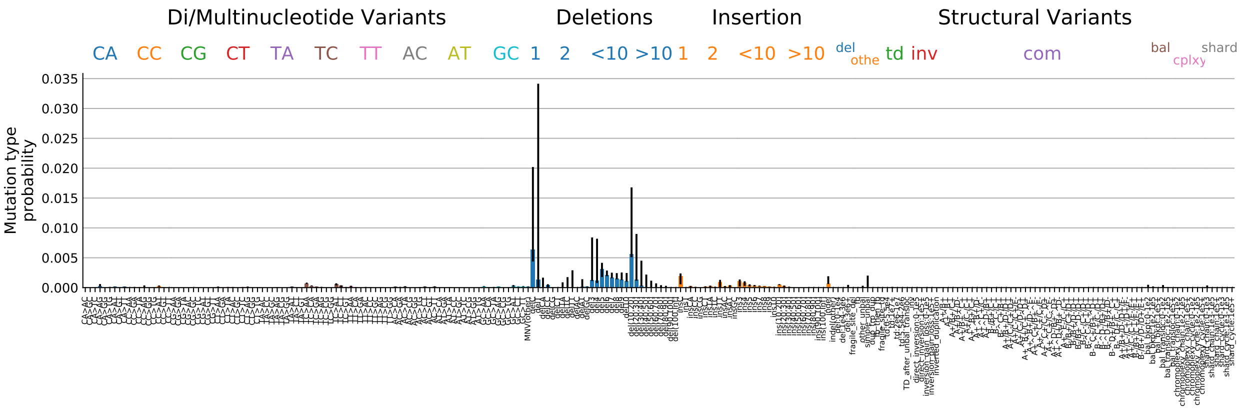

#### 4.4 Signature activity in different cancer types

Figure 19: Signature exposure in different cancer types.

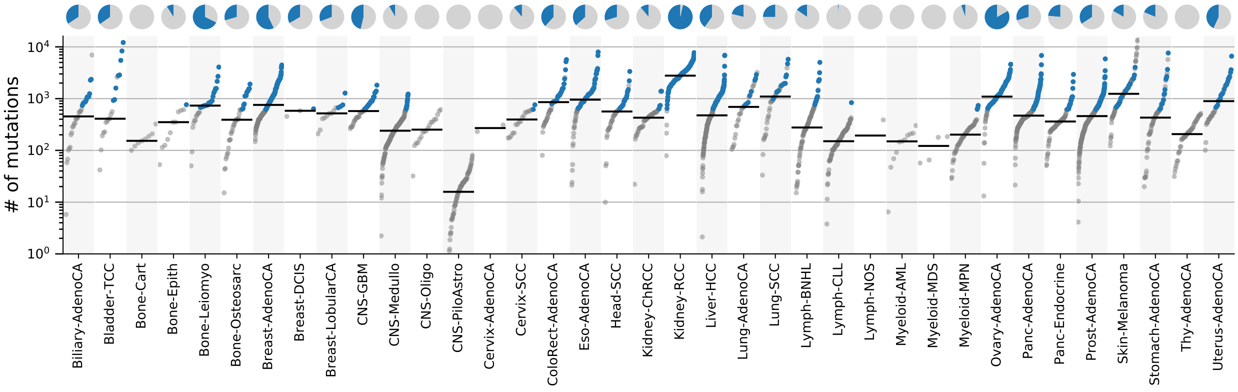

#### 4.5 Signature characterization

Figure 20: Signature specific tensor coefficients.

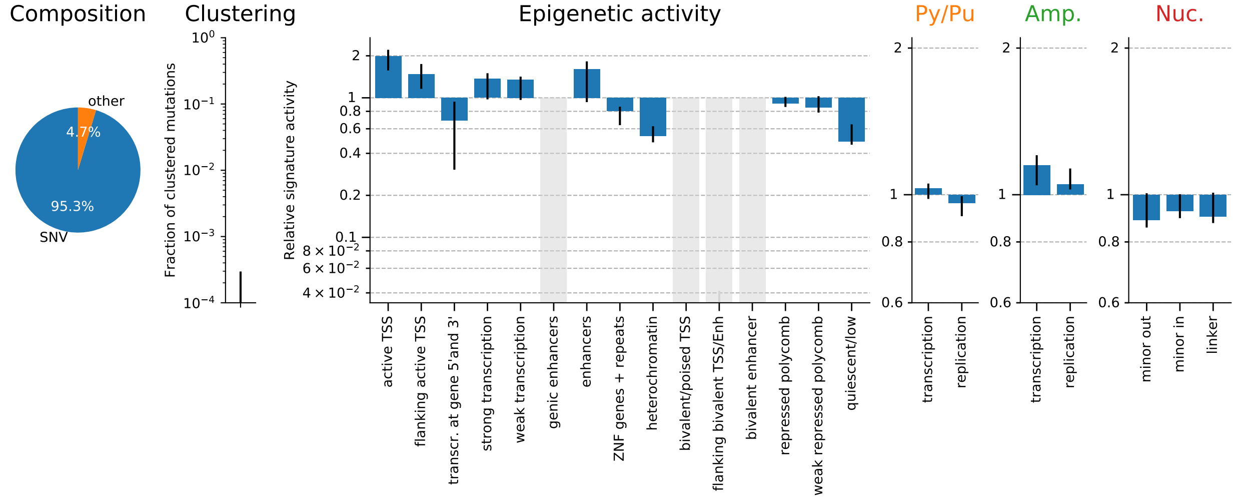

### 5 TS05-T[C>T]N (UV/GG-NER)

#### 5.1 Overall SNV spectrum

Figure 21: Single base substitution spectrum.

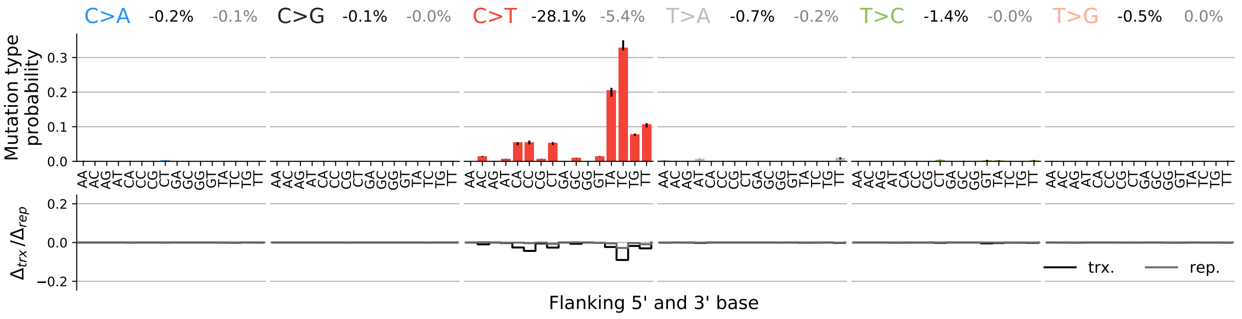

#### 5.2 SNV spectra for transcription and replication

Figure 22: Single base substitution spectra for template/coding and leading/lagging strand DNA.

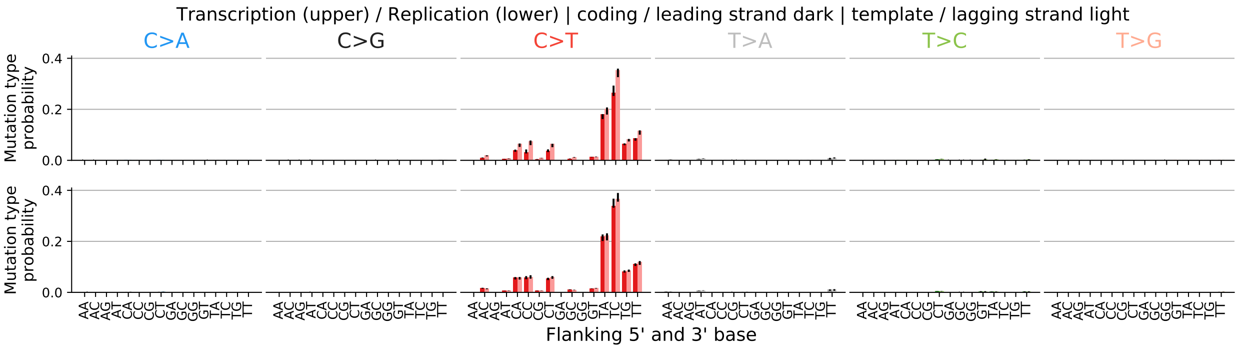

#### 5.3 Spectrum for other mutation types

Figure 23: Spectrum other mutation types.

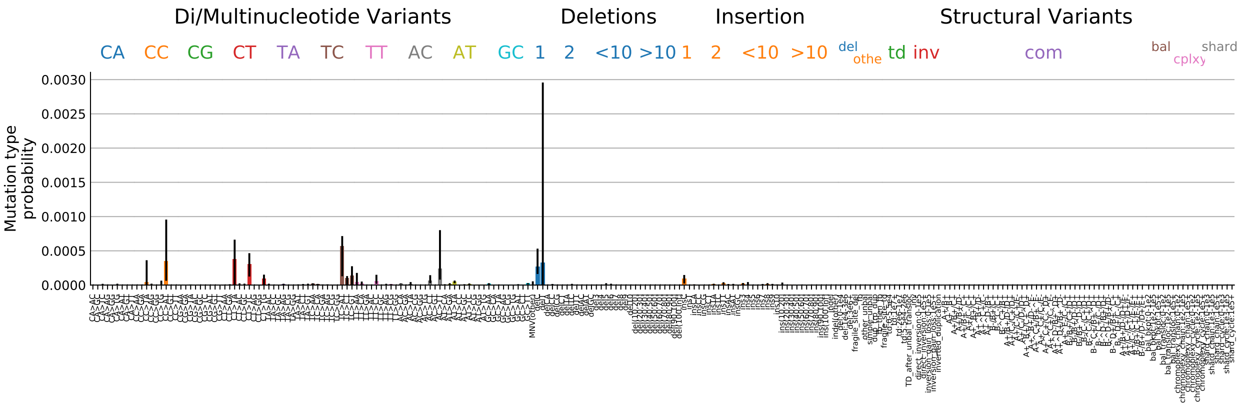

#### 5.4 Signature activity in different cancer types

Figure 24: Signature exposure in different cancer types.

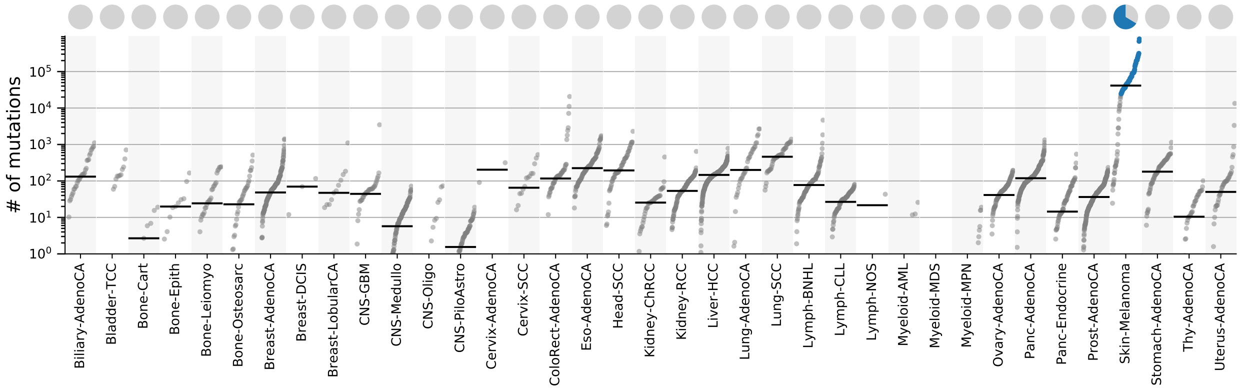

#### 5.5 Signature characterization

Figure 25: Signature specific tensor coefficients.

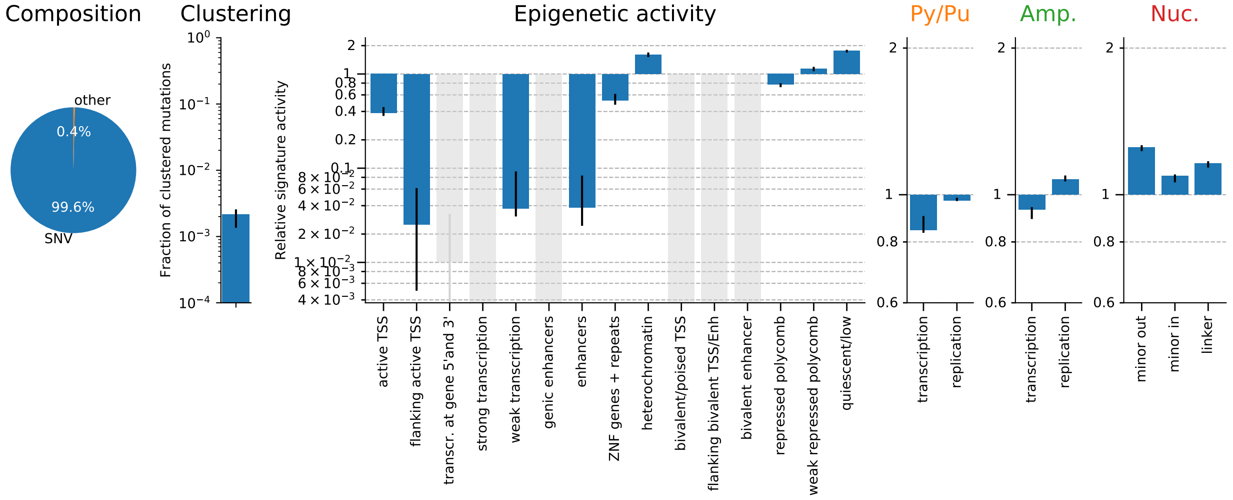

### 6 TS06-Y[C>T]N (UV/GG+TC-NER)

#### 6.1 Overall SNV spectrum

Figure 26: Single base substitution spectrum.

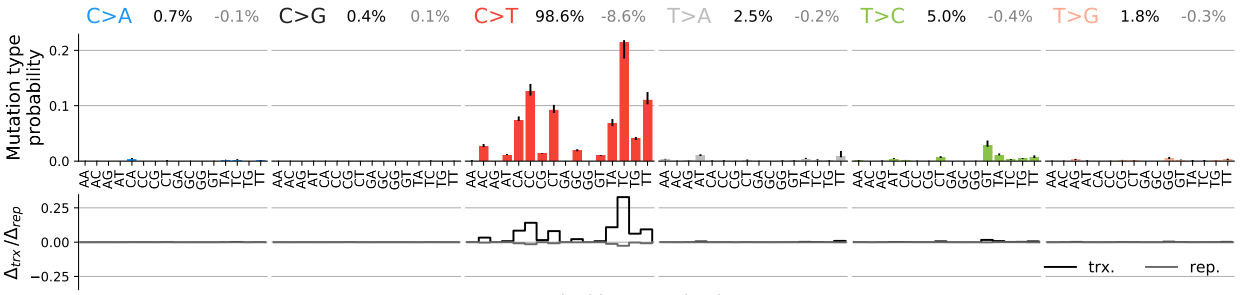

#### 6.2 SNV spectra for transcription and replication

Figure 27: Single base substitution spectra for template/coding and leading/lagging strand DNA.

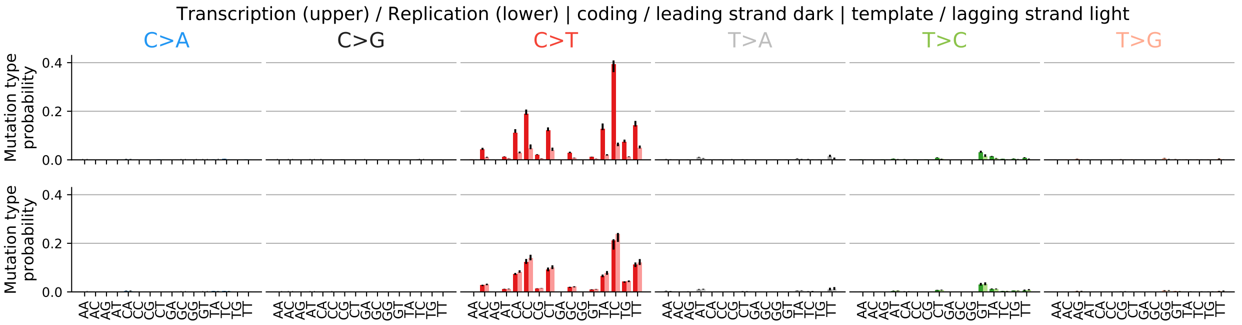

#### 6.3 Spectrum for other mutation types

Figure 28: Spectrum other mutation types.

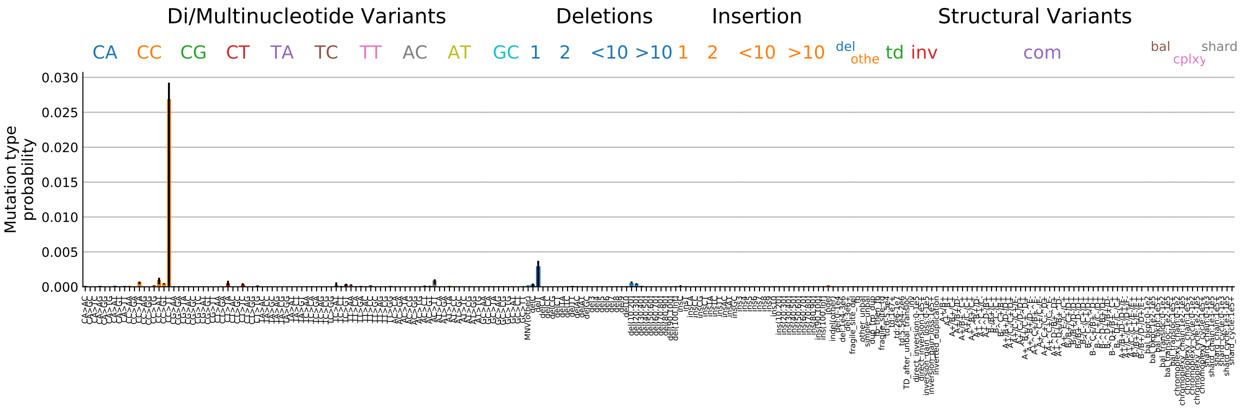

#### 6.4 Signature activity in different cancer types

Figure 29: Signature exposure in different cancer types.

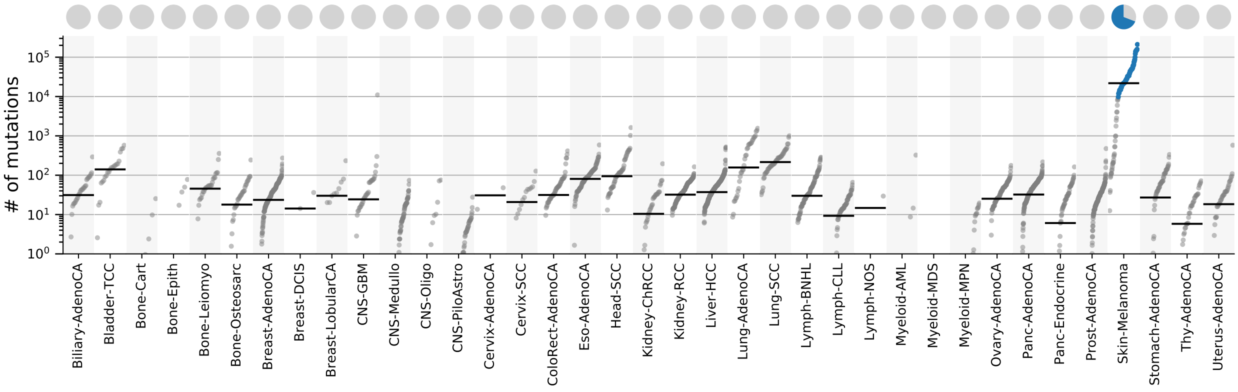

#### 6.5 Signature characterization

Figure 30: Signature specific tensor coefficients.

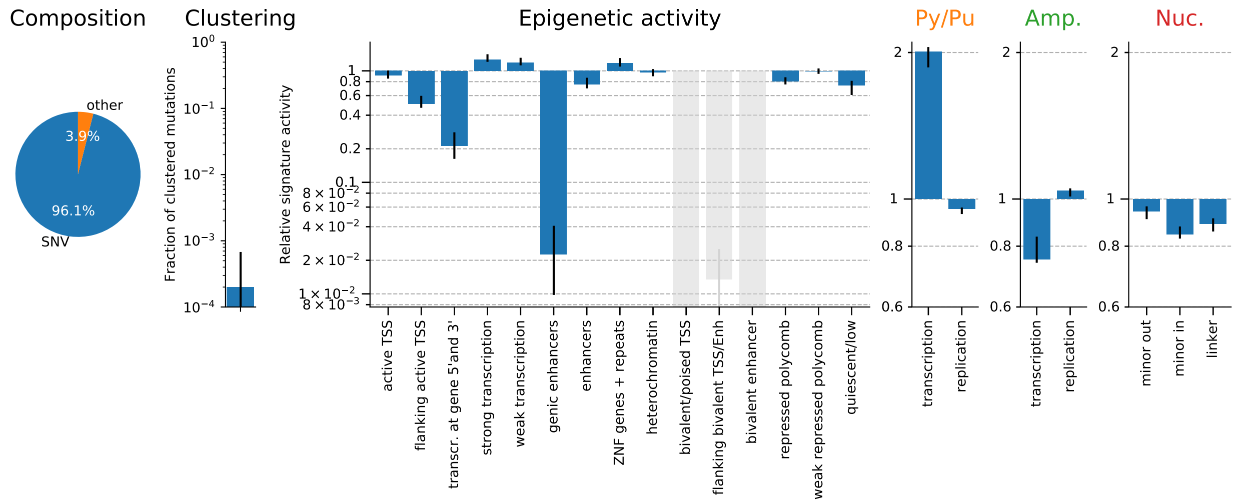

### 7 TS07-N[T>C]N (unknown)

#### 7.1 Overall SNV spectrum

Figure 31: Single base substitution spectrum.

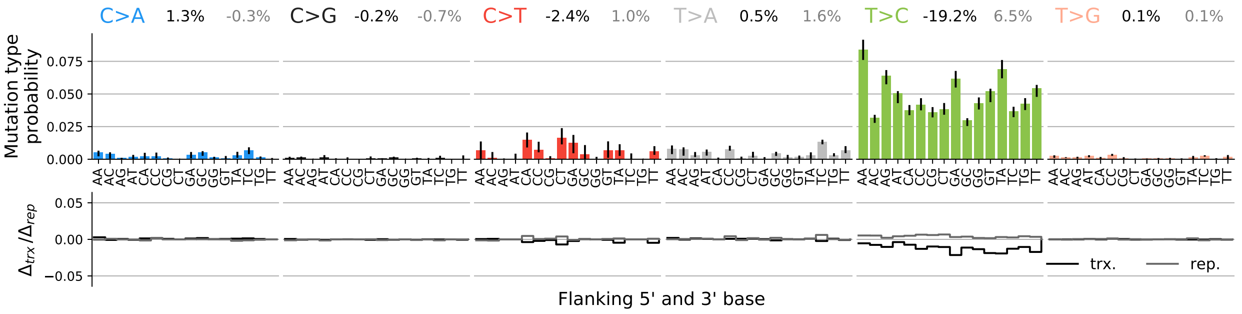

#### 7.2 SNV spectra for transcription and replication

Figure 32: Single base substitution spectra for template/coding and leading/lagging strand DNA.

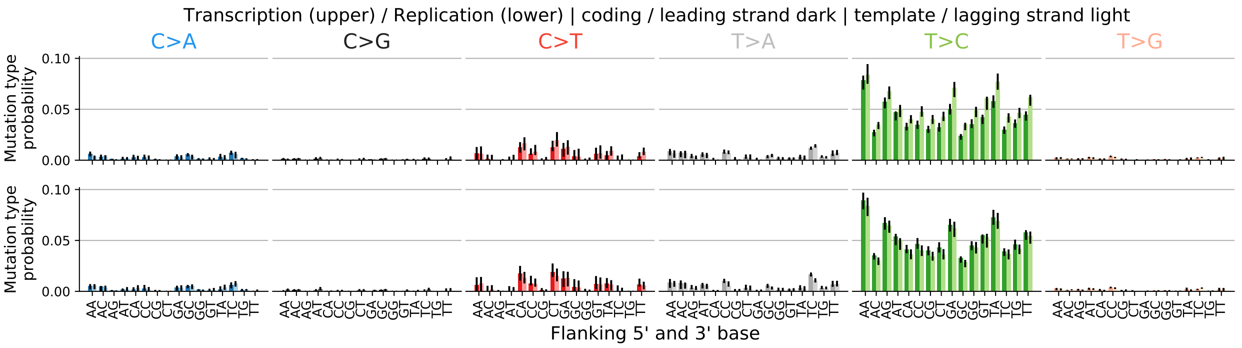

#### 7.3 Spectrum for other mutation types

Figure 33: Spectrum other mutation types.

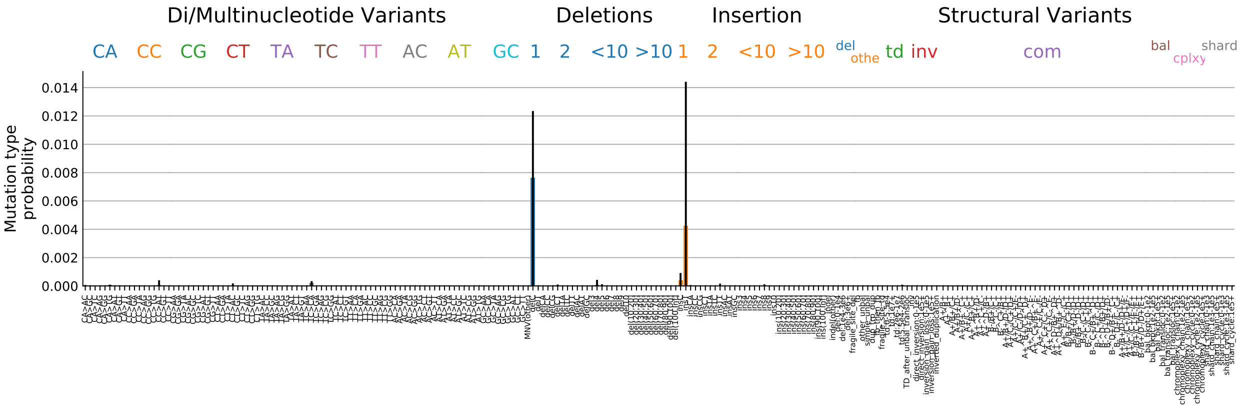

#### 7.4 Signature activity in different cancer types

Figure 34: Signature exposure in different cancer types.

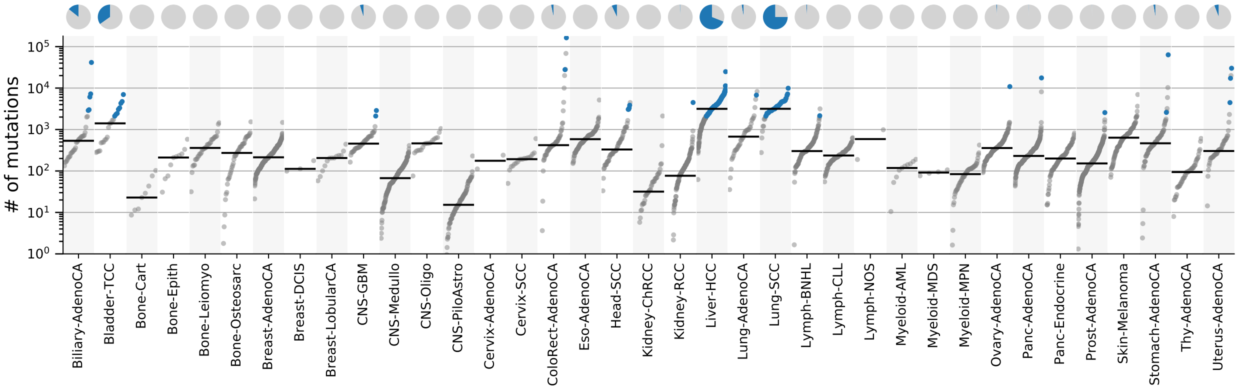

#### 7.5 Signature characterization

Figure 35: Signature specific tensor coefficients.

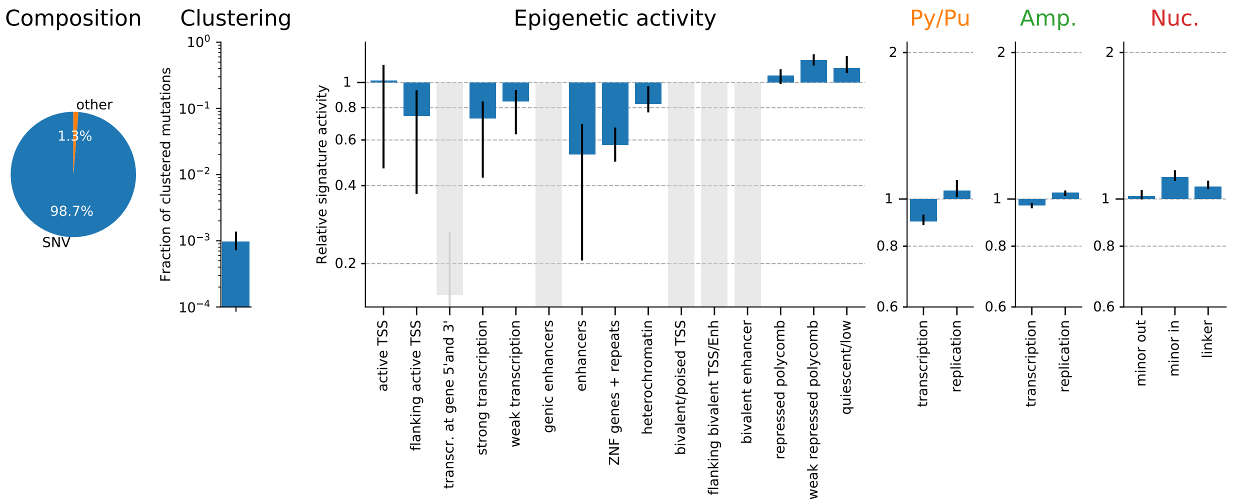

### 8 TS08-A[T>C]W (unknown/TAM)

#### 8.1 Overall SNV spectrum

Figure 36: Single base substitution spectrum.

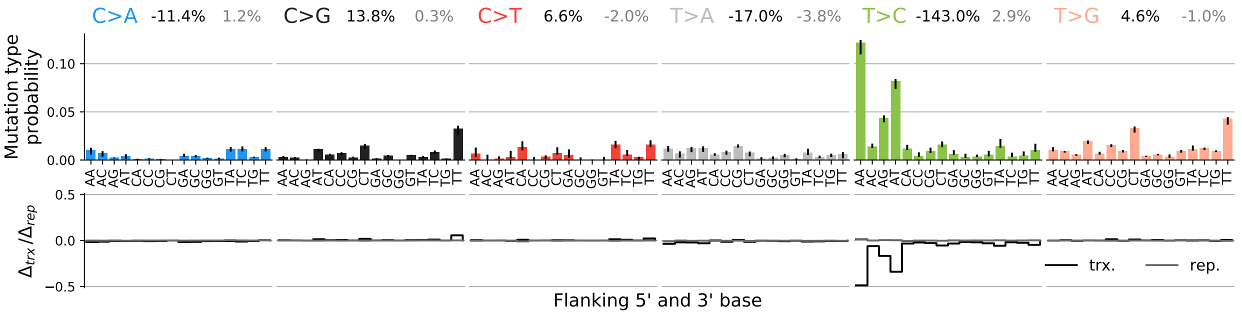

#### 8.2 SNV spectra for transcription and replication

Figure 37: Single base substitution spectra for template/coding and leading/lagging strand DNA.

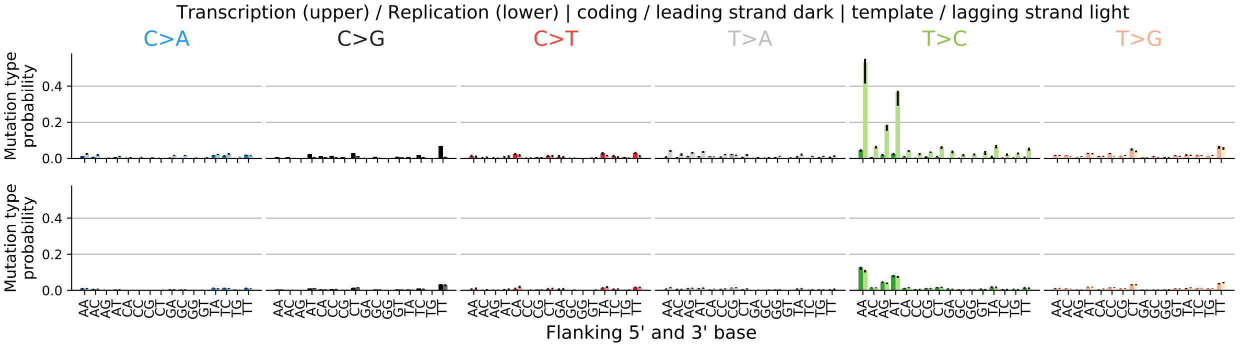

#### 8.3 Spectrum for other mutation types

Figure 38: Spectrum other mutation types.

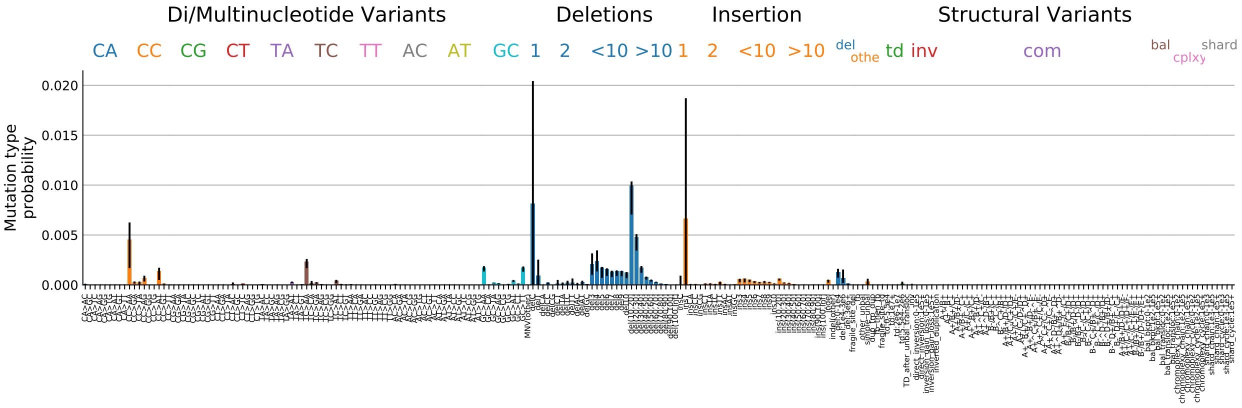

#### 8.4 Signature activity in different cancer types

Figure 39: Signature exposure in different cancer types.

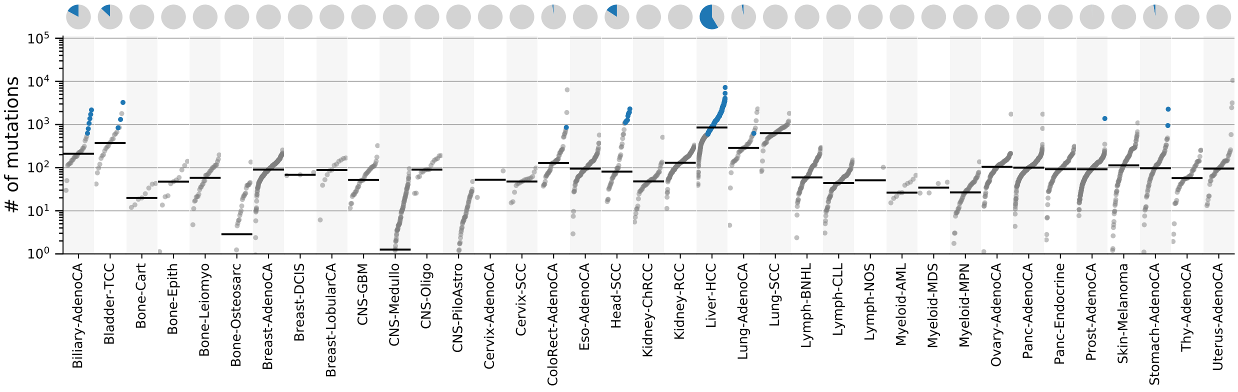

#### 8.5 Signature characterization

Figure 40: Signature specific tensor coefficients.

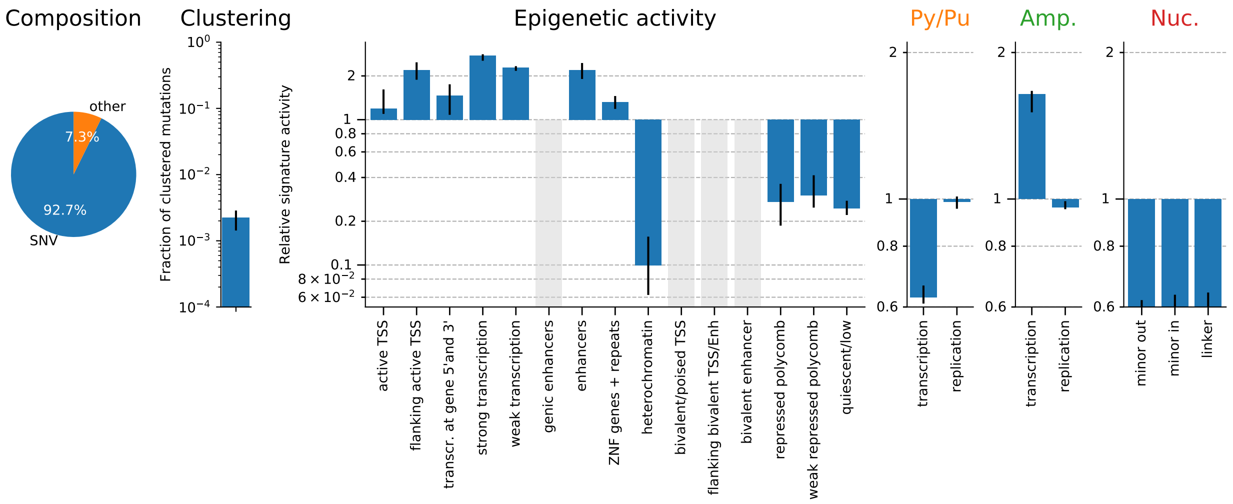

### 9 TS09-N[T>A]N (PAH/AA)

#### 9.1 Overall SNV spectrum

Figure 41: Single base substitution spectrum.

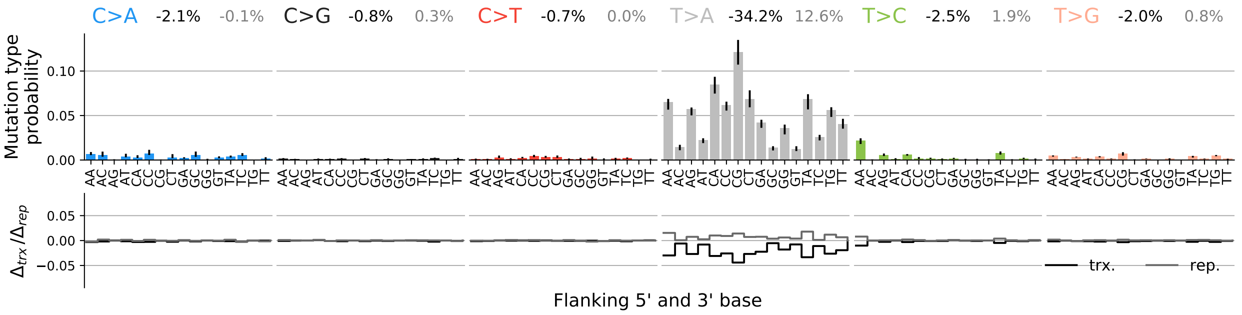

#### 9.2 SNV spectra for transcription and replication

Figure 42: Single base substitution spectra for template/coding and leading/lagging strand DNA.

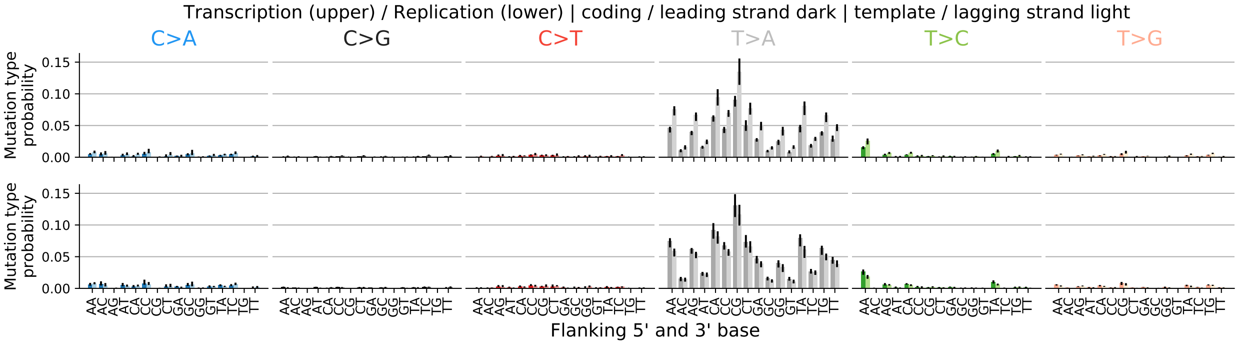

#### 9.3 Spectrum for other mutation types

Figure 43: Spectrum other mutation types.

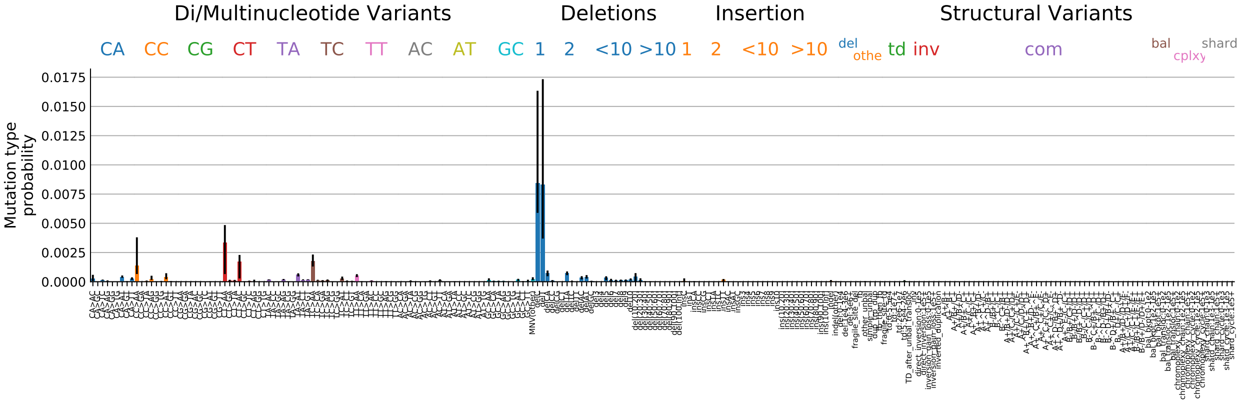

#### 9.4 Signature activity in different cancer types

Figure 44: Signature exposure in different cancer types.

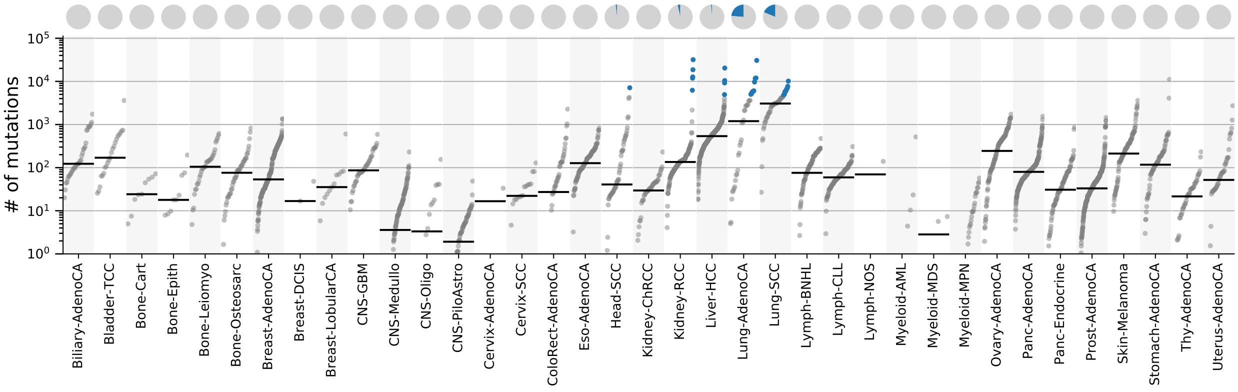

#### 9.5 Signature characterization

Figure 45: Signature specific tensor coefficients.

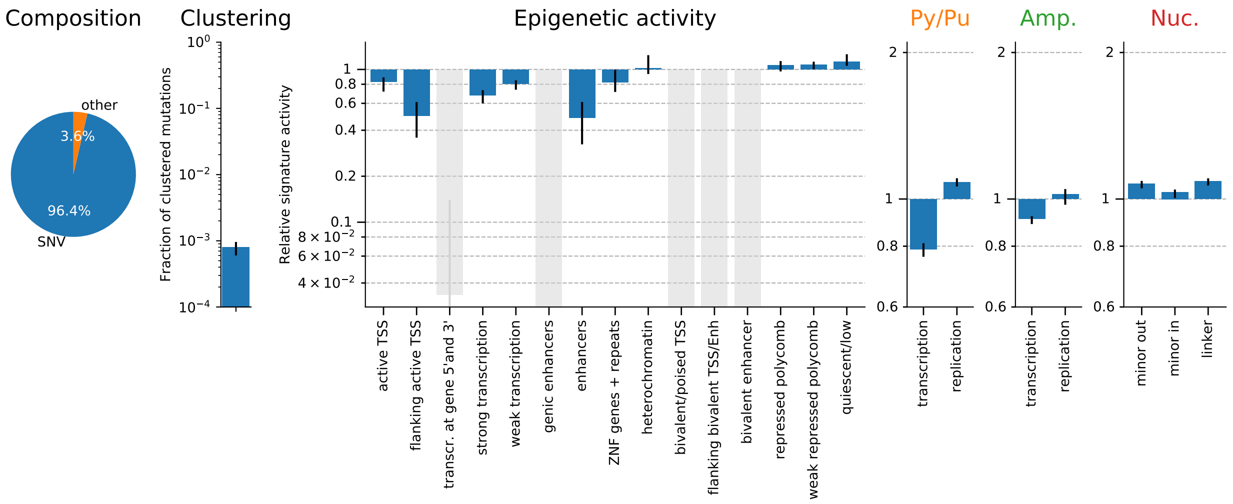

### 10 TS10-N[C>A]N (PAH/B[a]P)

#### 10.1 Overall SNV spectrum

Figure 46: Single base substitution spectrum.

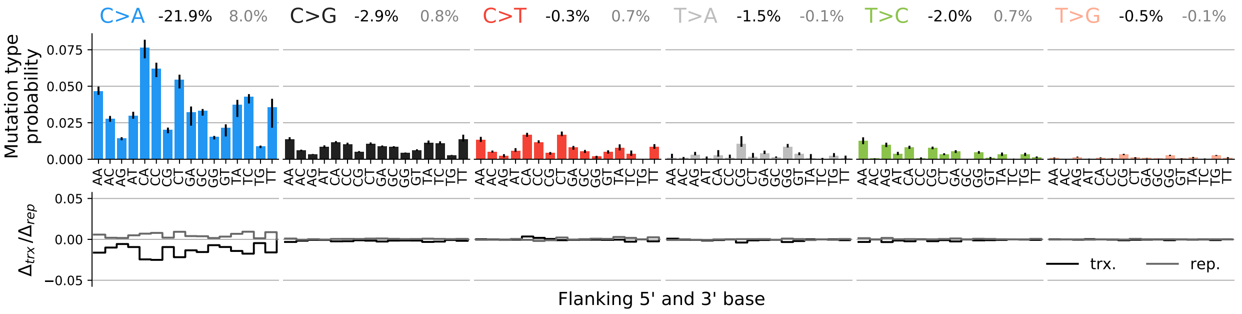

#### 10.2 SNV spectra for transcription and replication

Figure 47: Single base substitution spectra for template/coding and leading/lagging strand DNA.

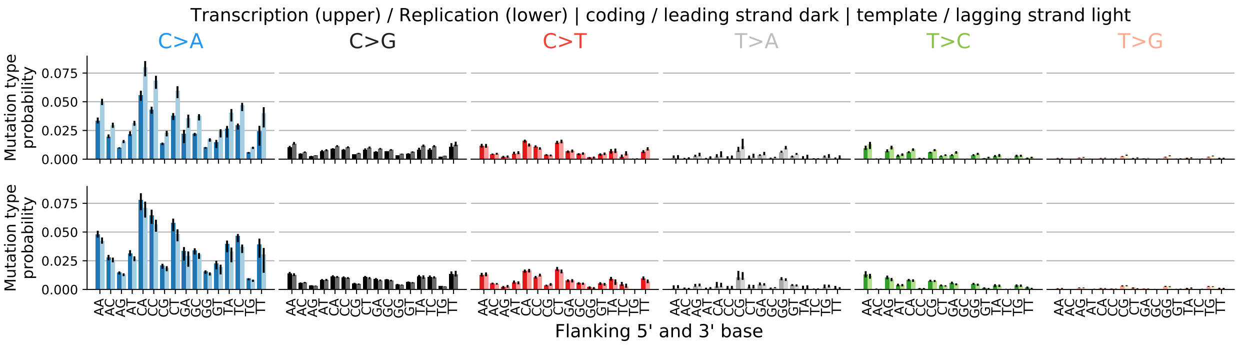

#### 10.3 Spectrum for other mutation types

Figure 48: Spectrum other mutation types.

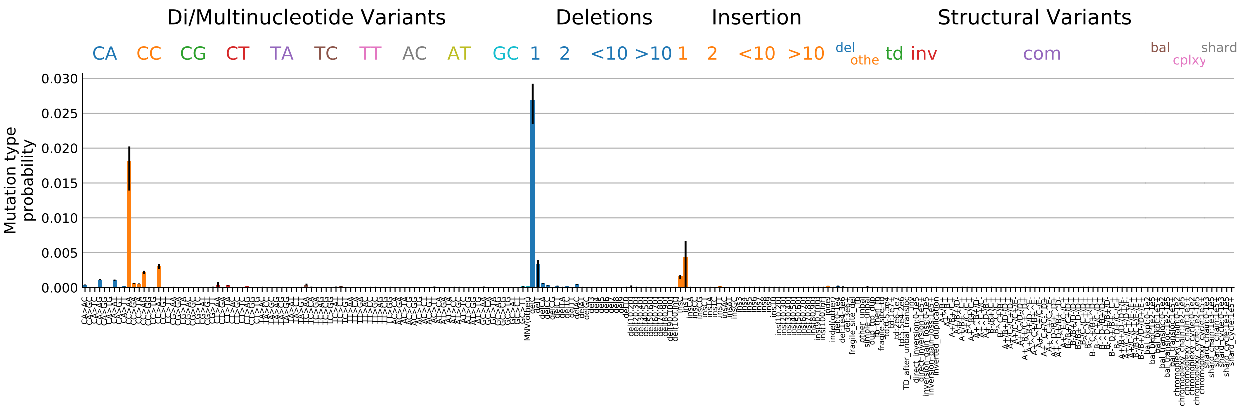

#### 10.4 Signature activity in different cancer types

Figure 49: Signature exposure in different cancer types.

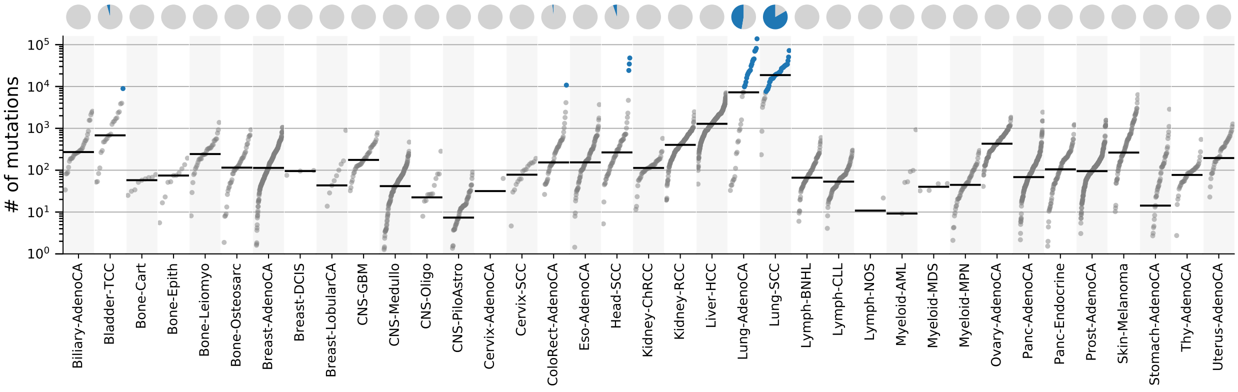

#### 10.5 Signature characterization

Figure 50: Signature specific tensor coefficients.

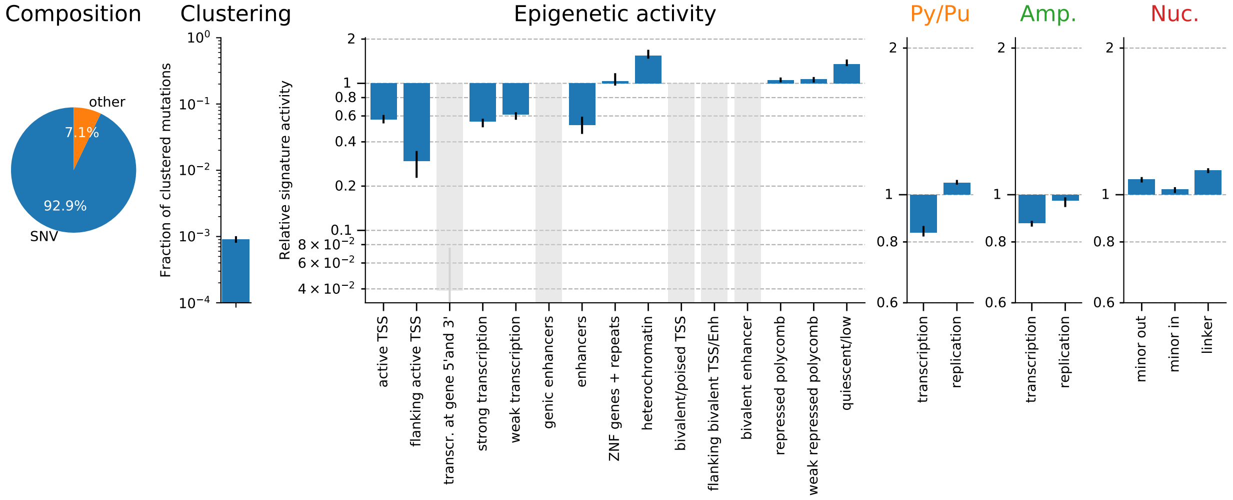

### 11 TS11-T[C>D]W;SV (APOBEC)

#### 11.1 Overall SNV spectrum

Figure 51: Single base substitution spectrum.

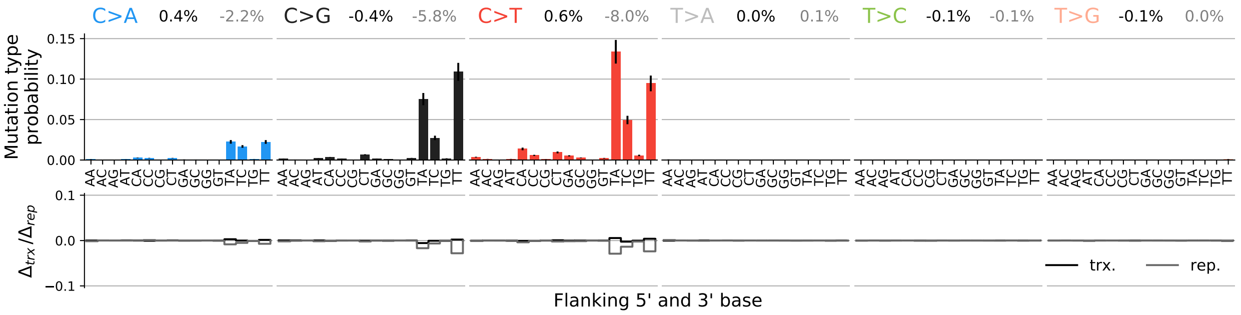

#### 11.2 SNV spectra for transcription and replication

Figure 52: Single base substitution spectra for template/coding and leading/lagging strand DNA.

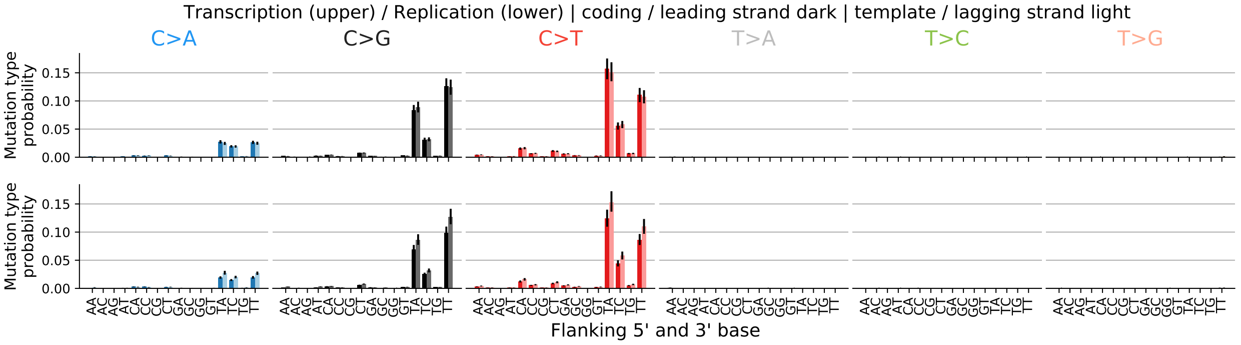

#### 11.3 Spectrum for other mutation types

Figure 53: Spectrum other mutation types.

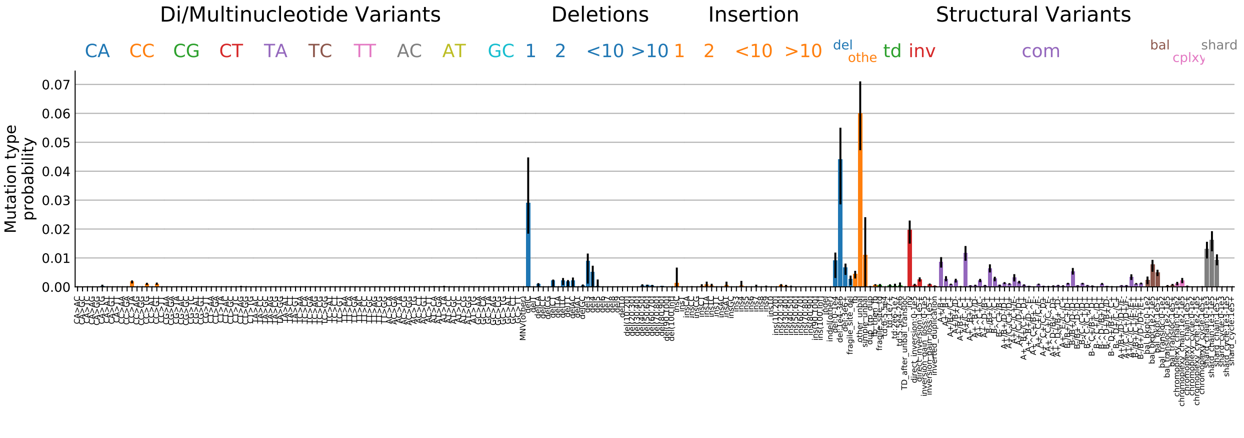

#### 11.4 Signature activity in different cancer types

Figure 54: Signature exposure in different cancer types.

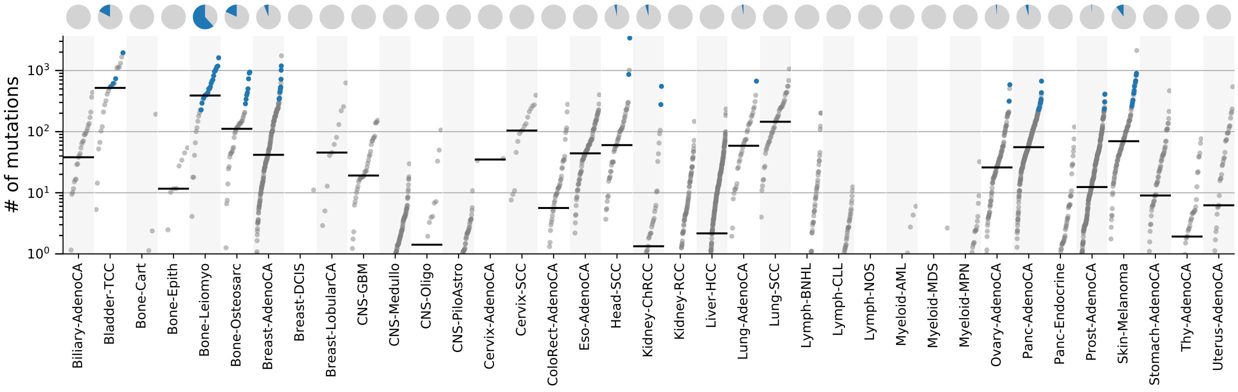

#### 11.5 Signature characterization

Figure 55: Signature specific tensor coefficients.

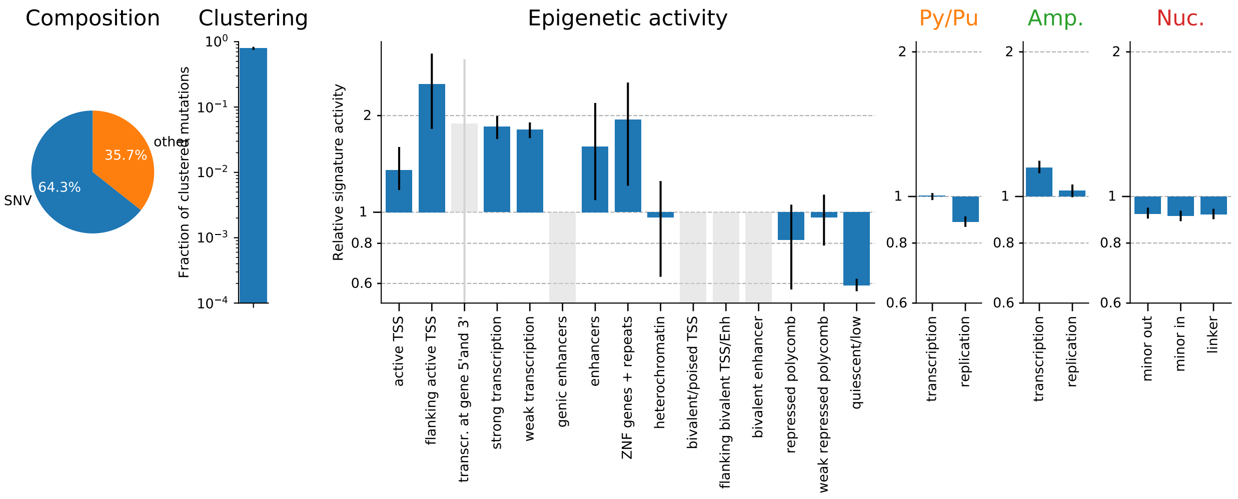

### 12 TS12-T[C>D]W (APOBEC)

#### 12.1 Overall SNV spectrum

Figure 56: Single base substitution spectrum.

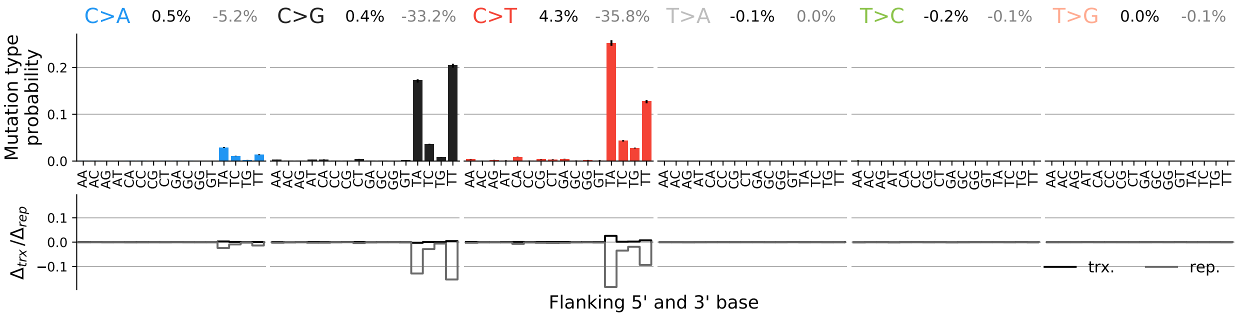

#### 12.2 SNV spectra for transcription and replication

Figure 57: Single base substitution spectra for template/coding and leading/lagging strand DNA.

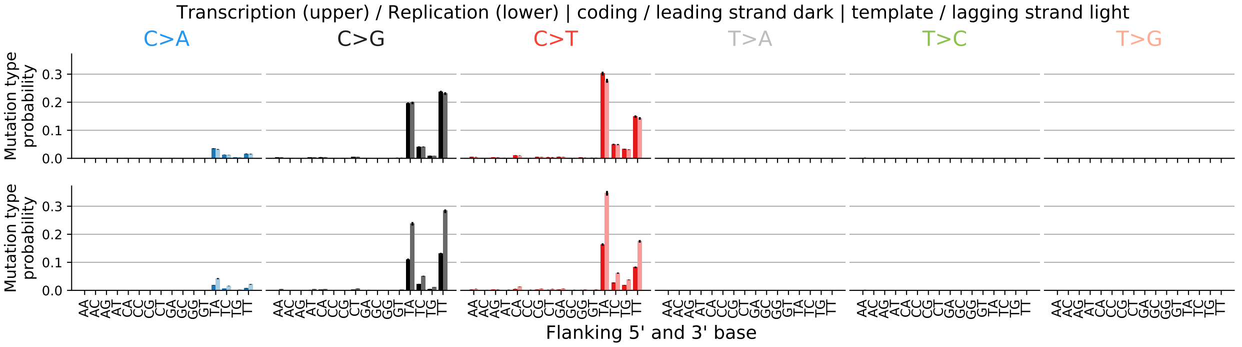

#### 12.3 Spectrum for other mutations type

Figure 58: Spectrum other mutation types.

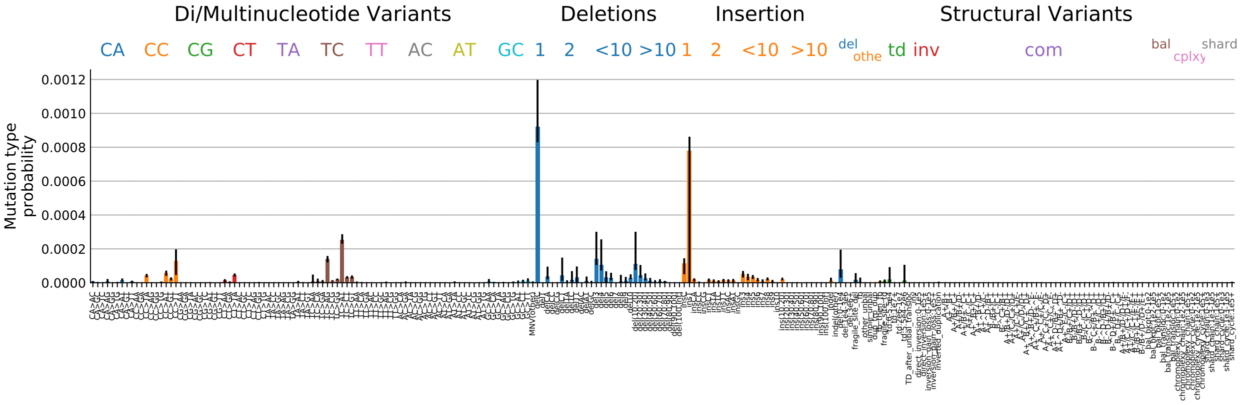

#### 12.4 Signature activity in different cancer types

Figure 59: Signature exposure in different cancer types.

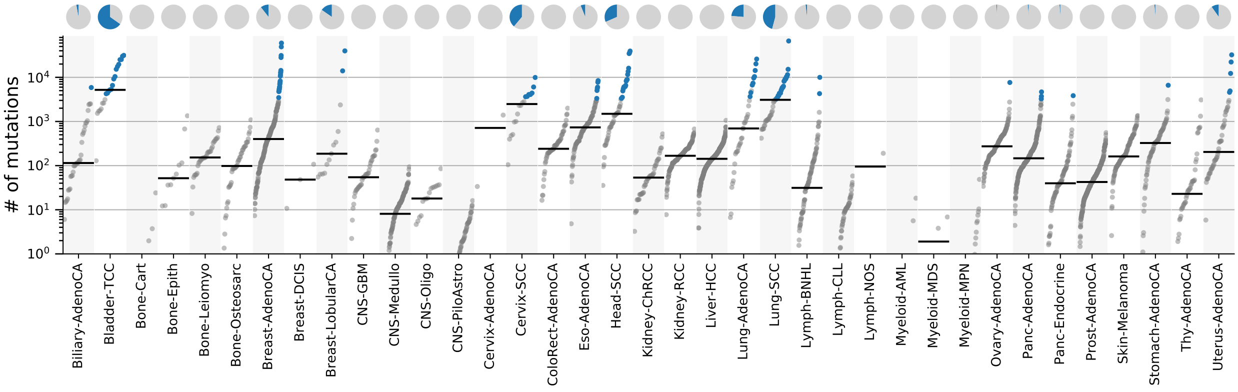

#### 12.5 Signature characterization

Figure 60: Signature specific tensor coefficients.

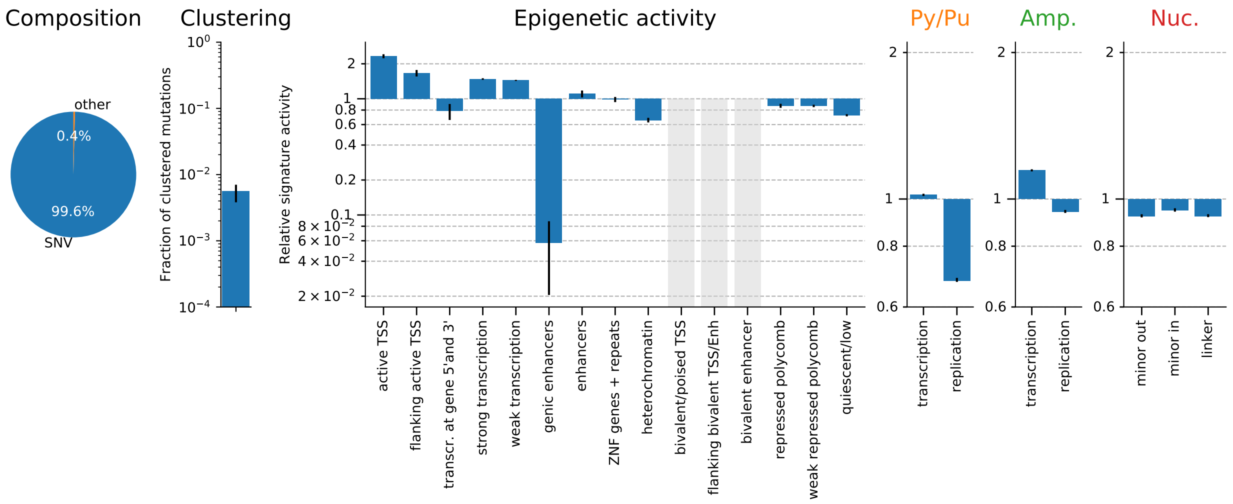

### 13 TS13-N[C>K]H (AID/SHM)

#### 13.1 Overall SNV spectrum

Figure 61: Single base substitution spectrum.

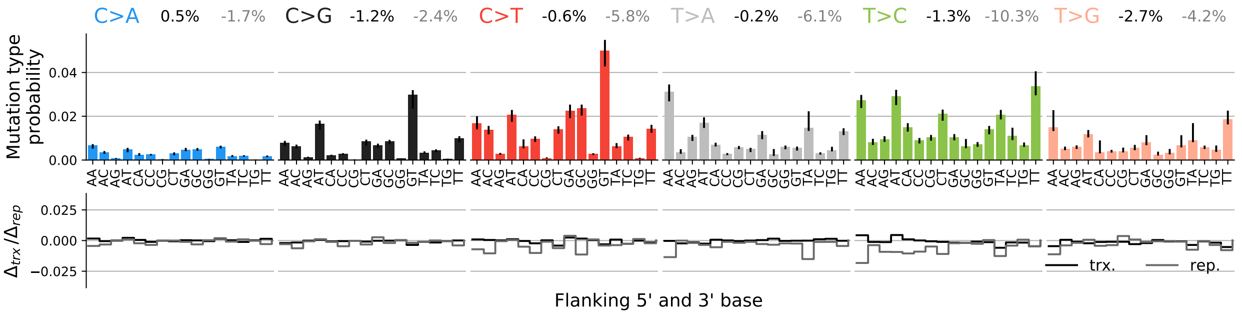

#### 13.2 SNV spectra for transcription and replication

Figure 62: Single base substitution spectra for template/coding and leading/lagging strand DNA.

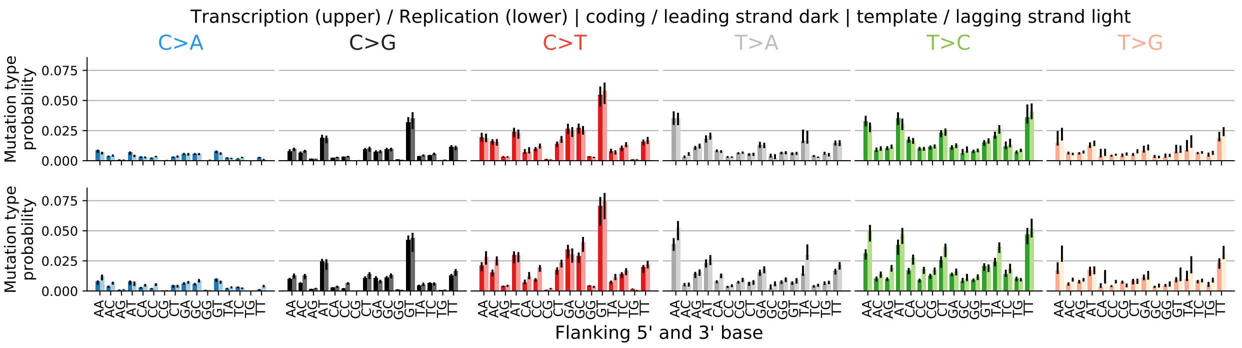

#### 13.3 Spectrum for other mutation types

Figure 63: Spectrum other mutation types.

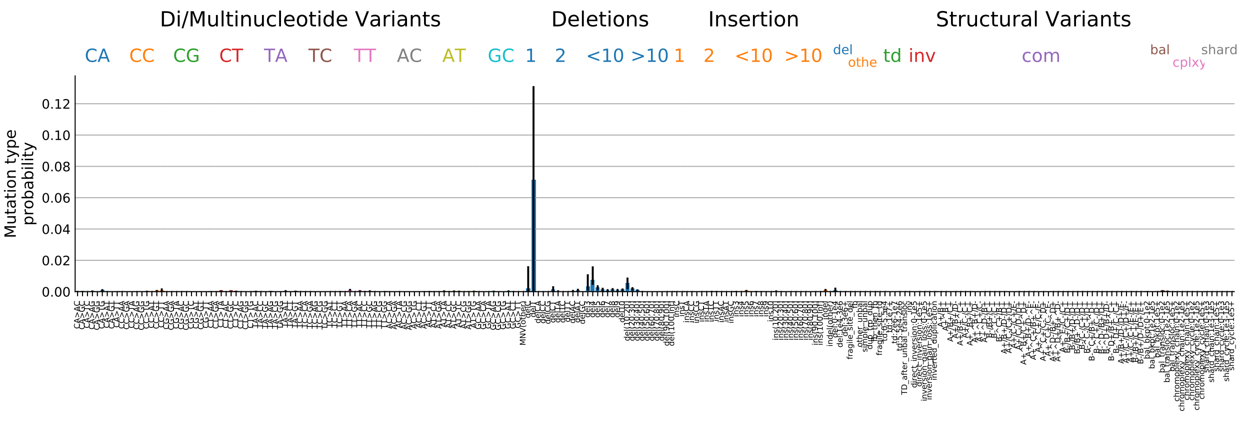

#### 13.4 Signature activity in different cancer types

Figure 64: Signature exposure in different cancer types.

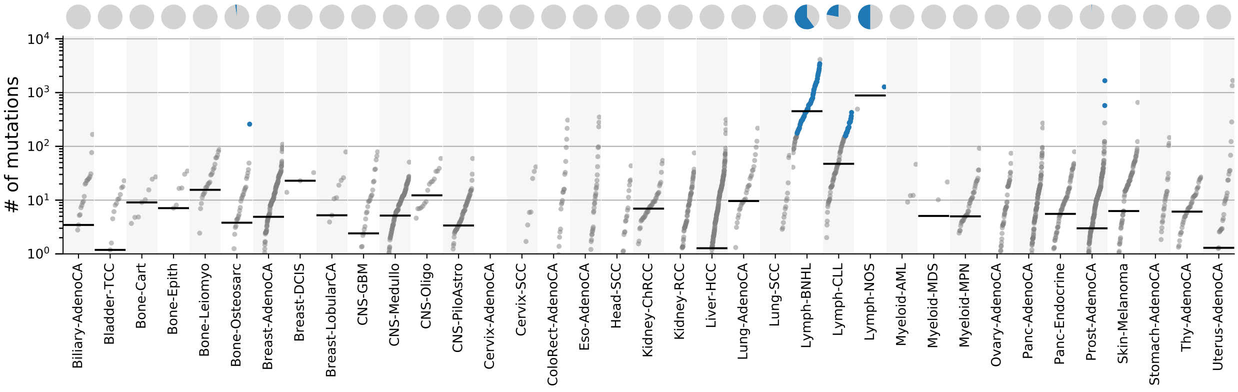

#### 13.5 Signature characterization

Figure 65: Signature specific tensor coefficients.

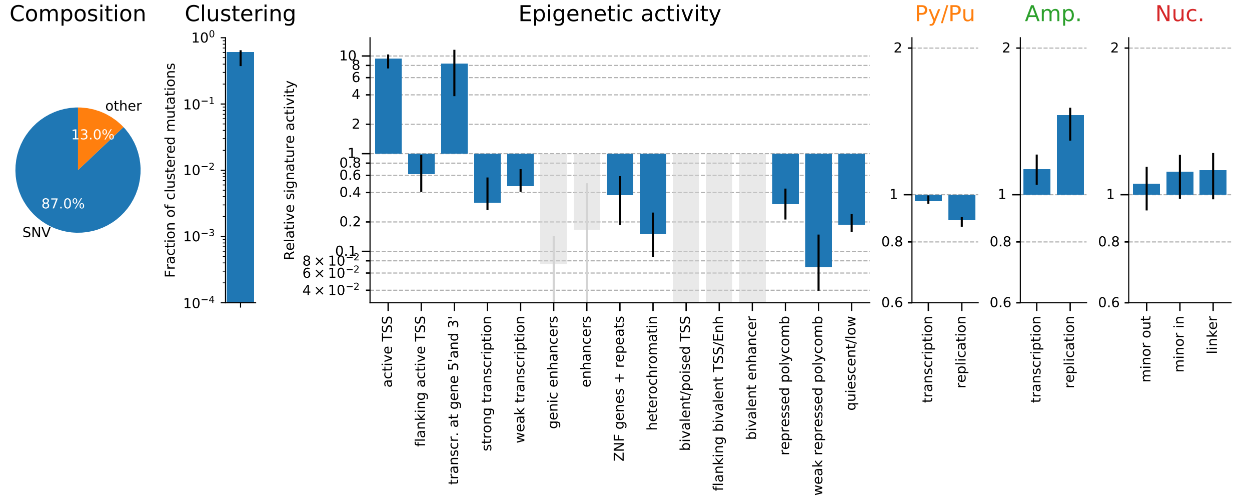

### 14 TS14-W[T>V]W (POLH)

#### 14.1 Overall SNV spectrum

Figure 66: Single base substitution spectrum.

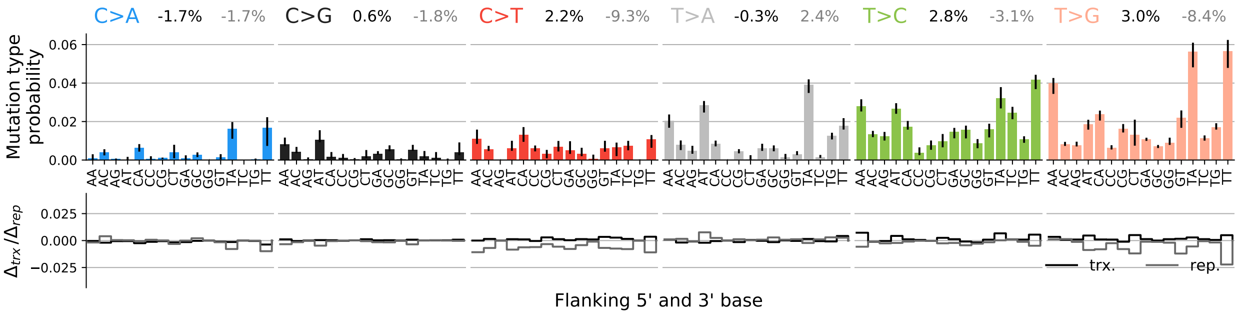

#### 14.2 SNV spectra for transcription and replication

Figure 67: Single base substitution spectra for template/coding and leading/lagging strand DNA.

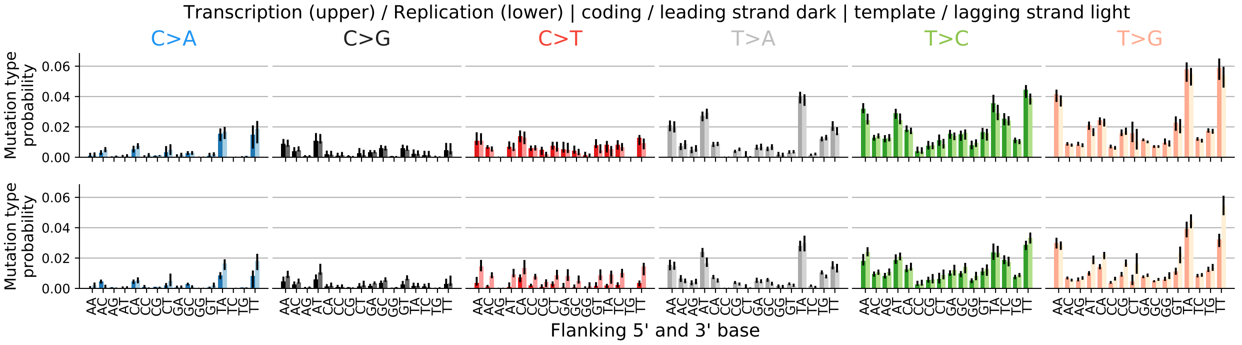

#### 14.3 Spectrum for other mutation types

Figure 68: Spectrum other mutation types.

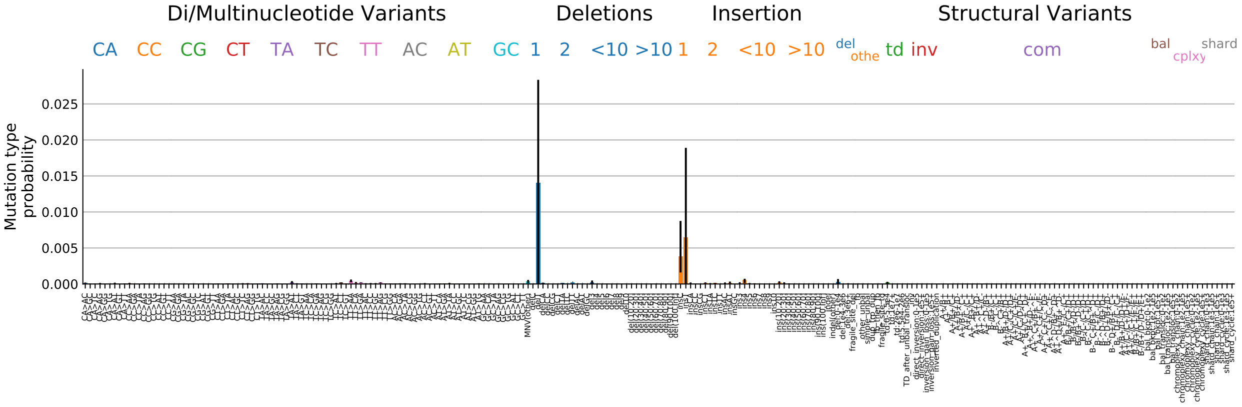

#### 14.4 Signature activity in different cancer types

Figure 69: Signature exposure in different cancer types.

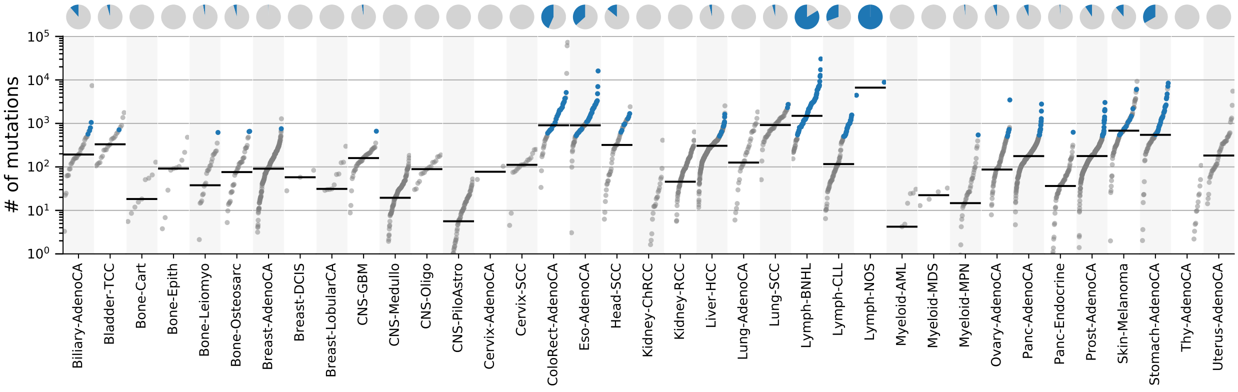

#### 14.5 Signature characterization

Figure 70: Signature specific tensor coefficients.

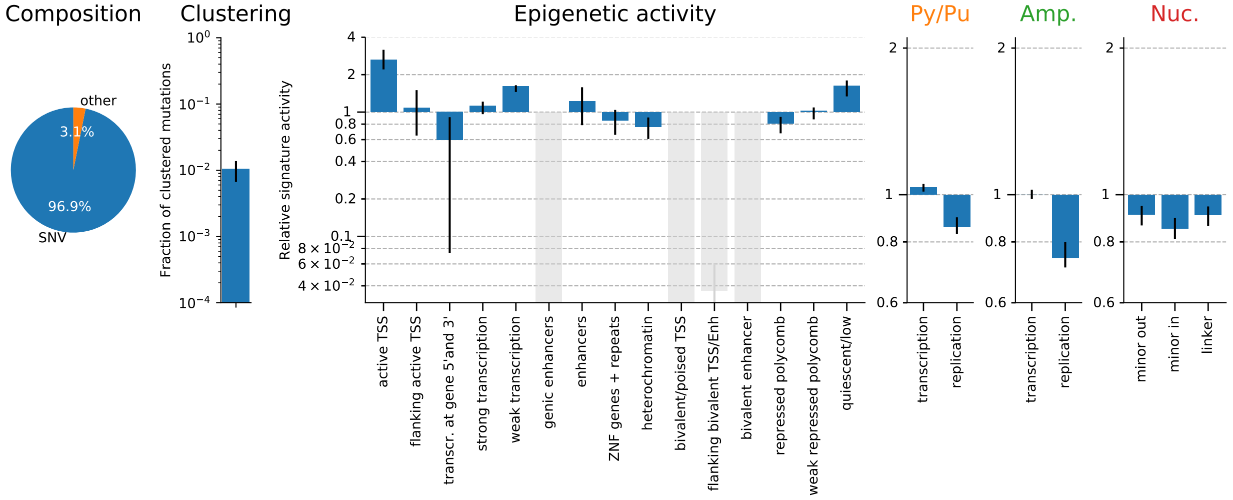

### 15 TS15-G[C>T]N;ID (MMRD)

#### 15.1 Overall SNV spectrum

Figure 71: Single base substitution spectrum.

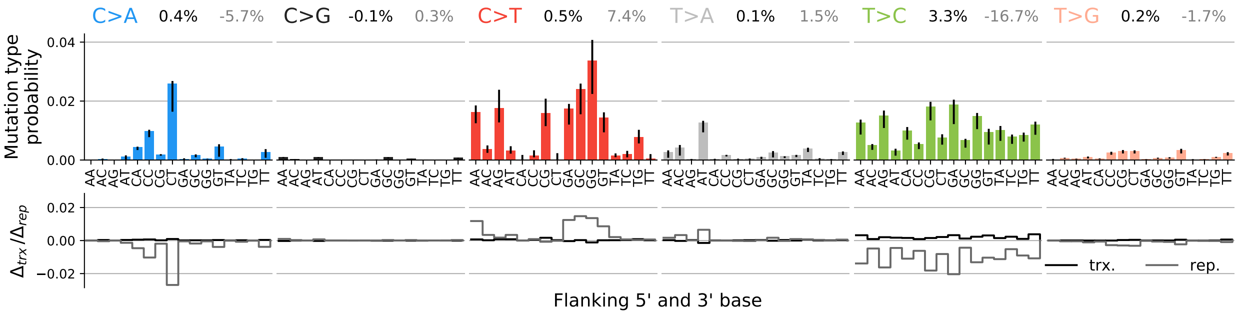

#### 15.2 SNV spectra for transcription and replication

Figure 72: Single base substitution spectra for template/coding and leading/lagging strand DNA.

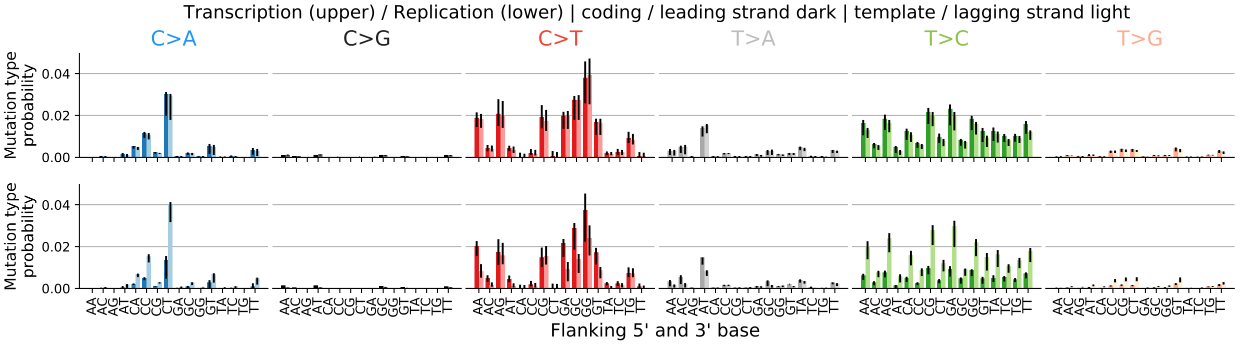

#### 15.3 Spectrum for other mutation types

Figure 73: Spectrum other mutation types.

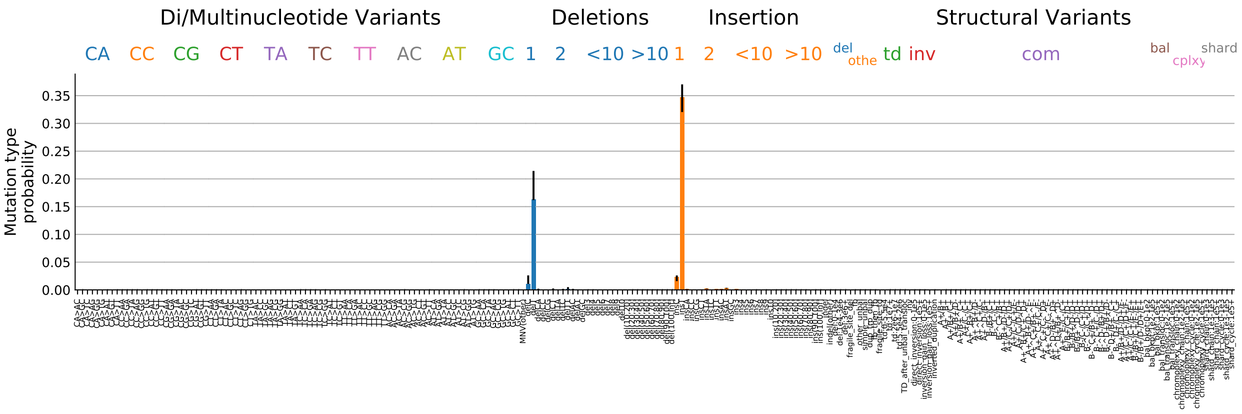

#### 15.4 Signature activity in different cancer types

Figure 74: Signature exposure in different cancer types.

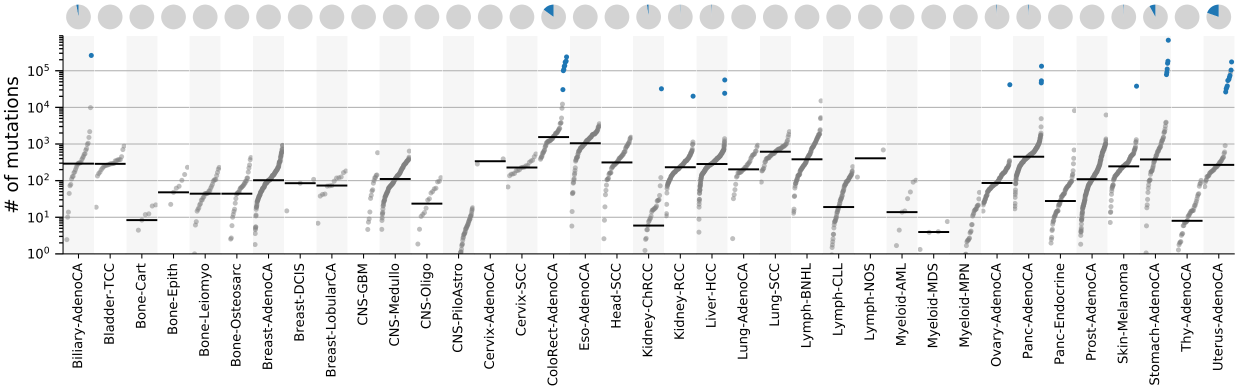

#### 15.5 Signature characterization

Figure 75: Signature specific tensor coefficients.

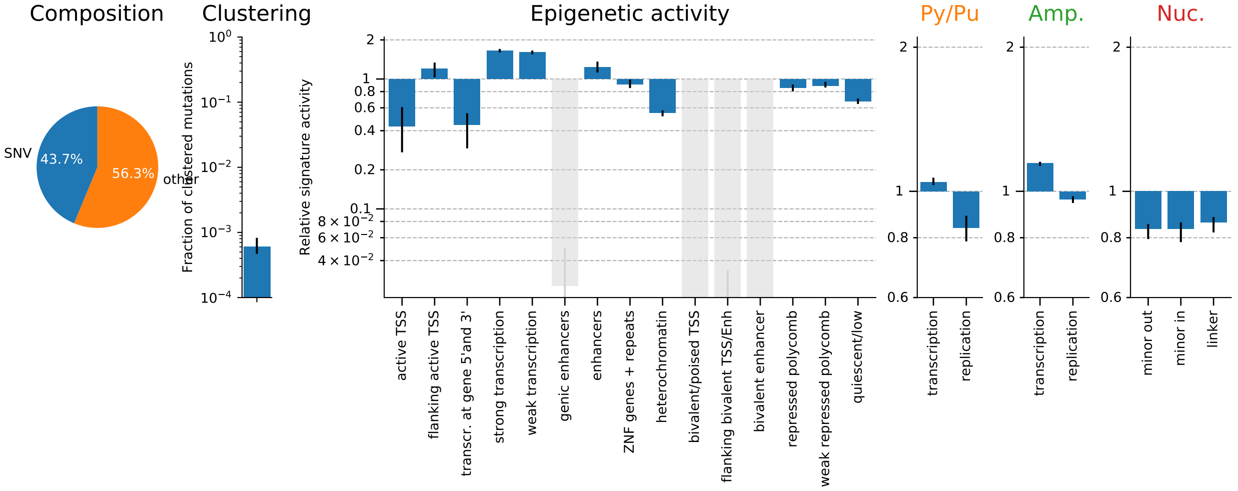

### 16 TS16-N[C>A]T;ID (MMRD:POLE-exo)

#### 16.1 Overall SNV spectrum

Figure 76: Single base substitution spectrum.

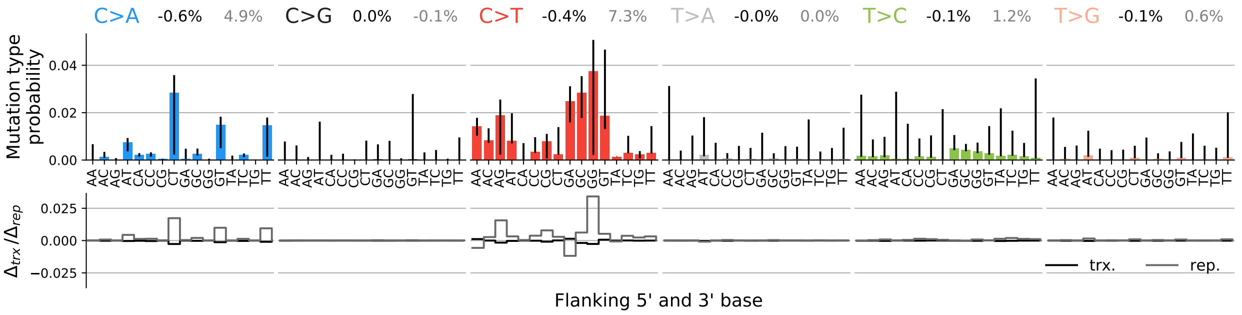

#### 16.2 SNV spectra for transcription and replication

Figure 77: Single base substitution spectra for template/coding and leading/lagging strand DNA.

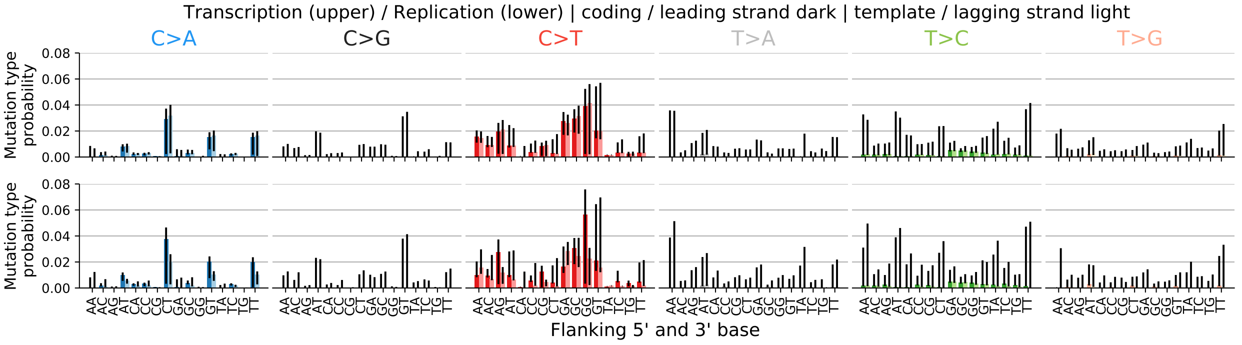

#### 16.3 Spectrum for other mutation types

Figure 78: Spectrum other mutation types.

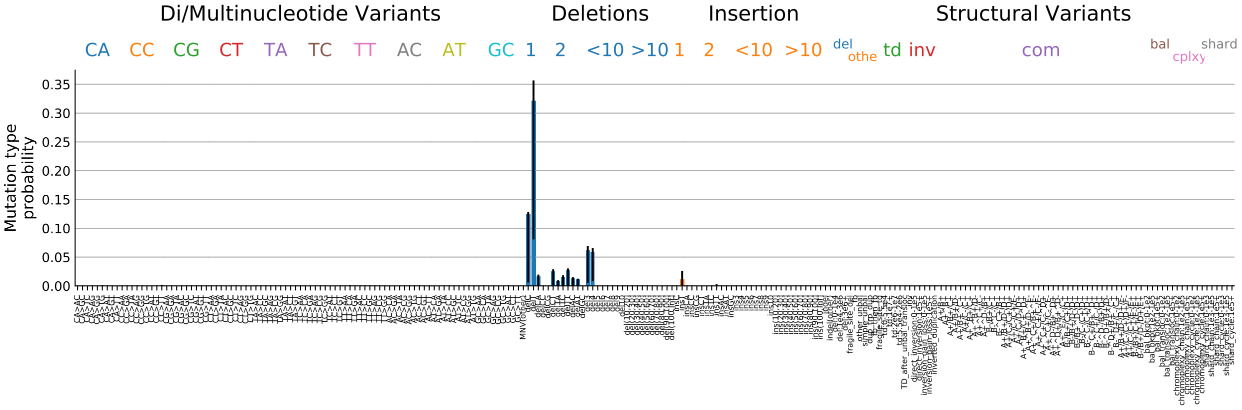

#### 16.4 Signature activity in different cancer types

Figure 79: Signature exposure in different cancer types.

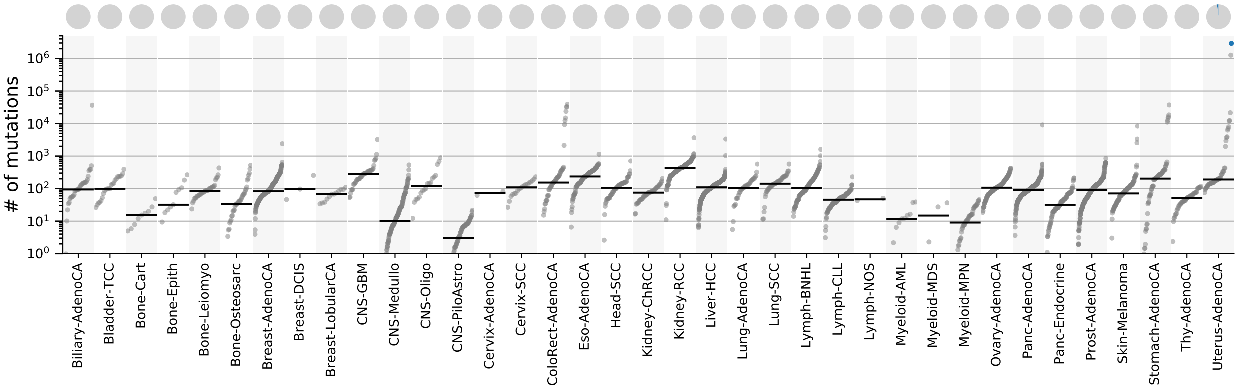

#### 16.5 Signature characterization

Figure 80: Signature specific tensor coefficients.

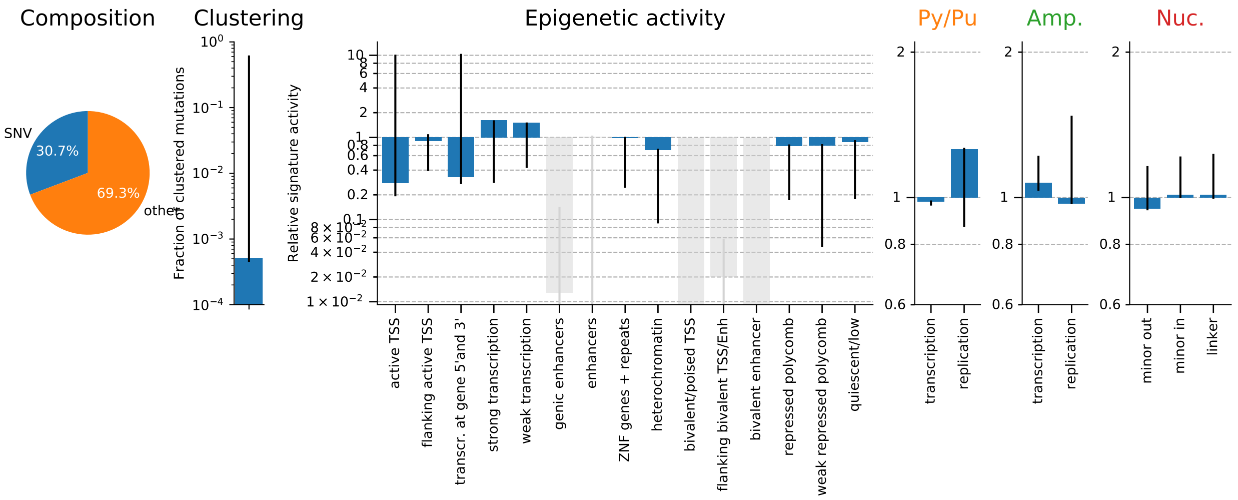

### 17 TS17-T[C>A]T (POLE-exo)

#### 17.1 Overall SNV spectrum

Figure 81: Single base substitution spectrum.

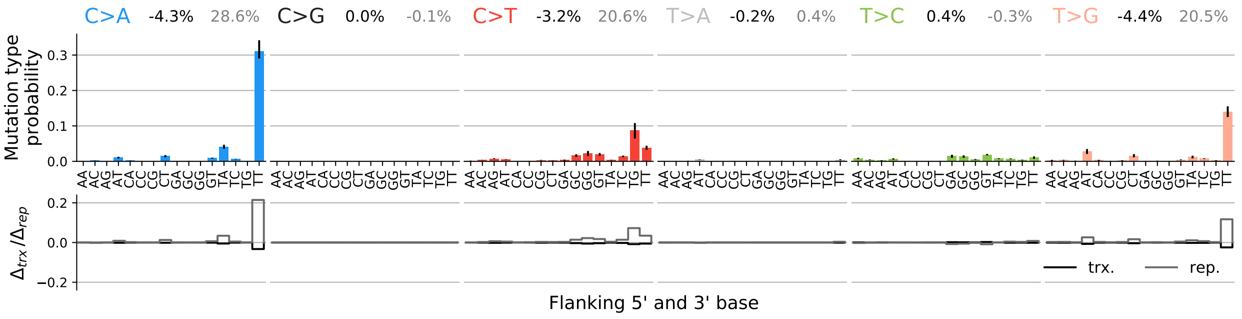

#### 17.2 SNV spectra for transcription and replication

Figure 82: Single base substitution spectra for template/coding and leading/lagging strand DNA.

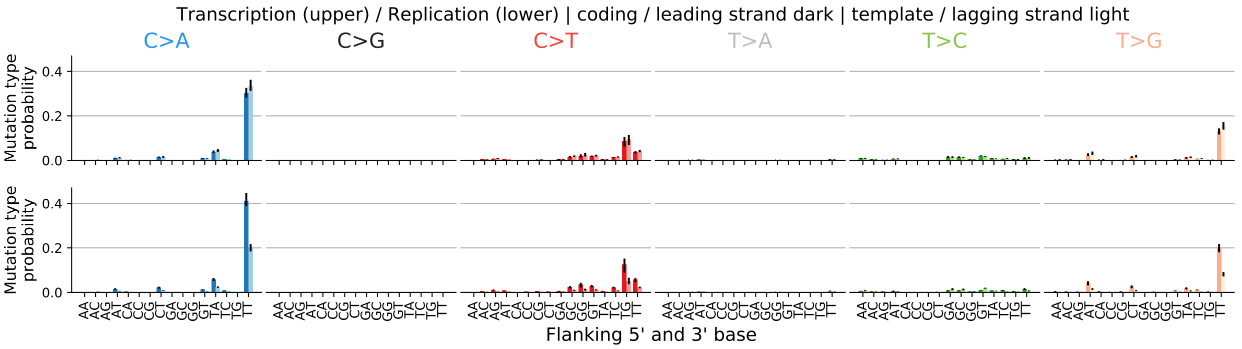

#### 17.3 Spectrum for other mutation types

Figure 83: Spectrum other mutation types.

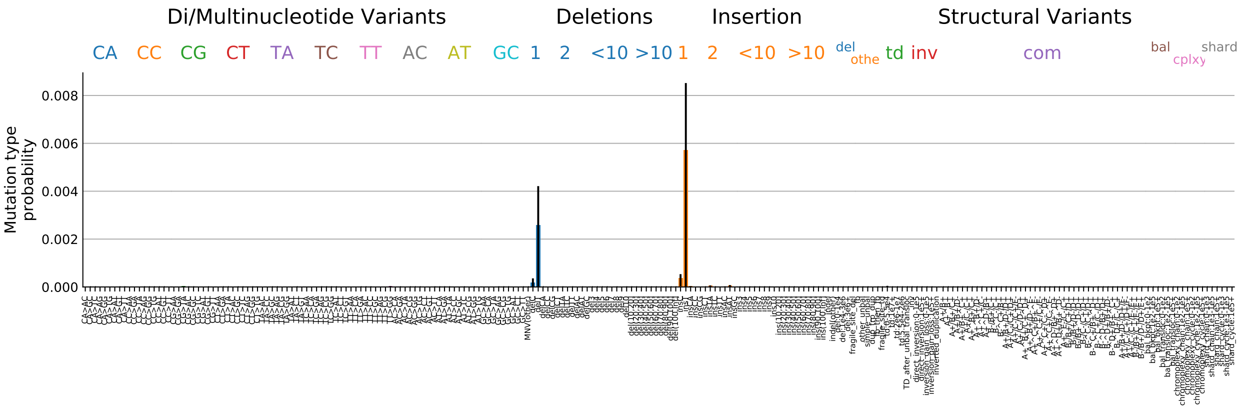

#### 17.4 Signature activity in different cancer types

Figure 84: Signature exposure in different cancer types.

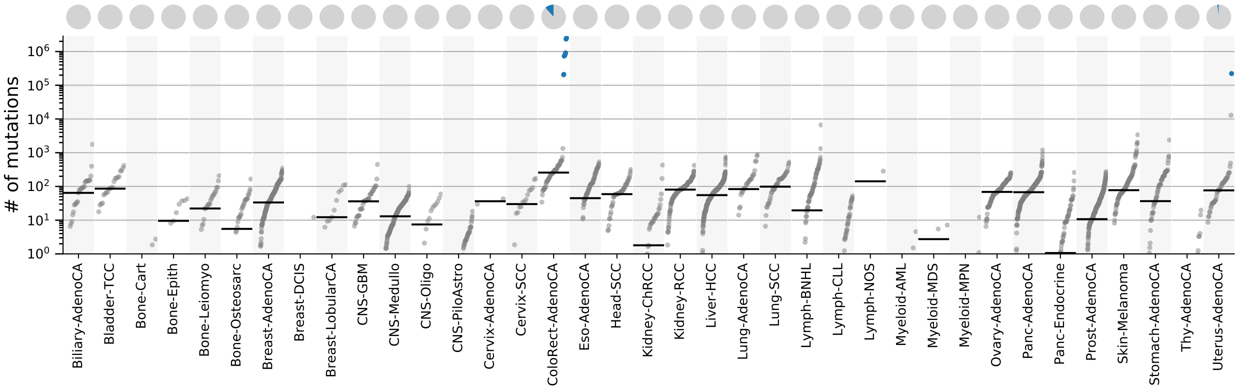

#### 17.5 Signature characterization

Figure 85: Signature specific tensor coefficients.

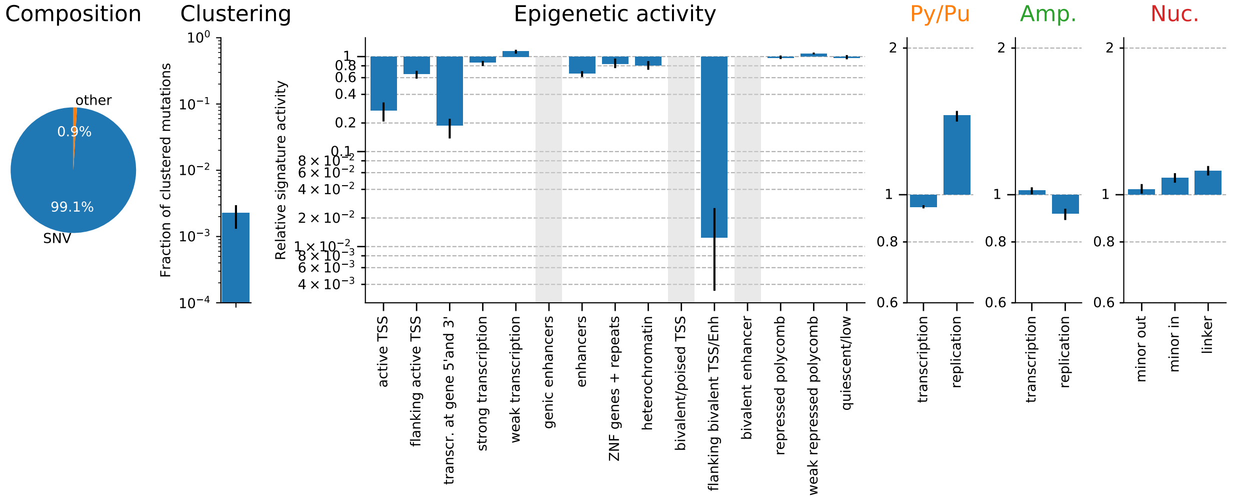

### 18 TS18-N[C>A]W (BERD/MUTYH)

#### 18.1 Overall SNV spectrum

Figure 86: Single base substitution spectrum.

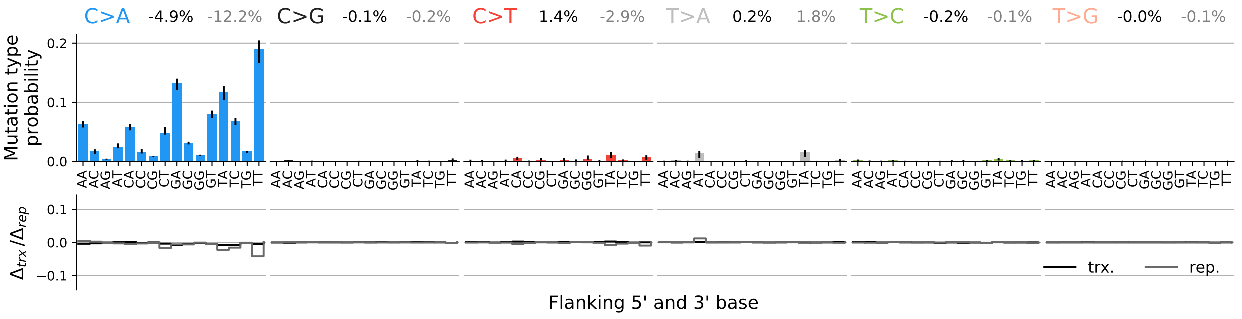

#### 18.2 SNV spectra for transcription and replication

Figure 87: Single base substitution spectra for template/coding and leading/lagging strand DNA.

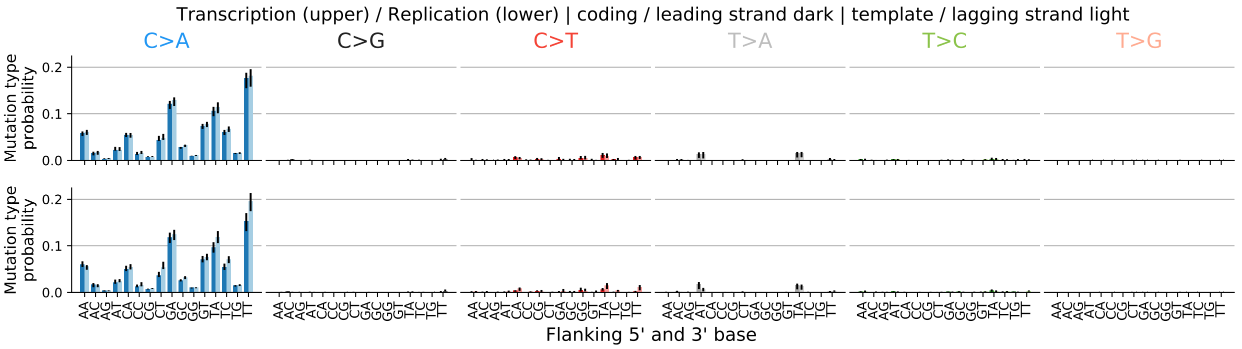

#### 18.3 Spectrum for other mutation types

Figure 88: Spectrum other mutation types.

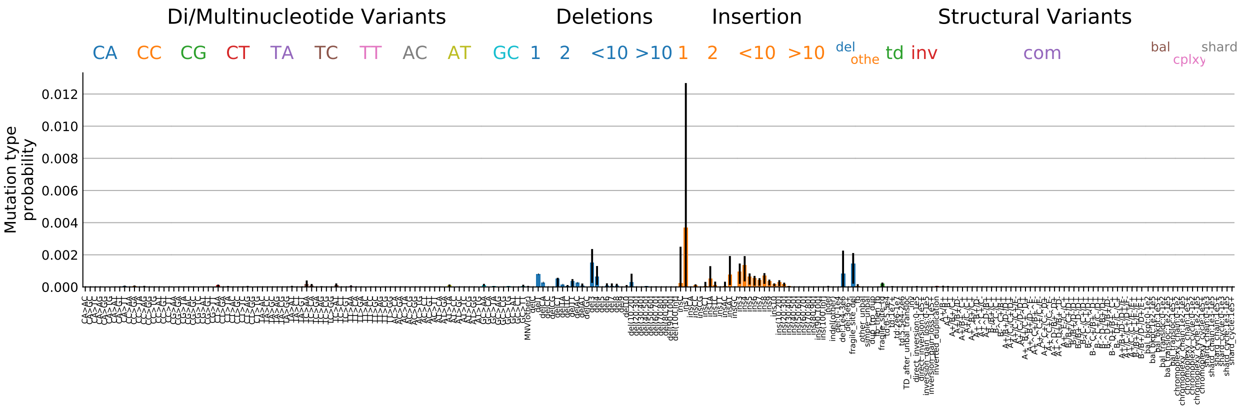

#### 18.4 Signature activity in different cancer types

Figure 89: Signature exposure in different cancer types.

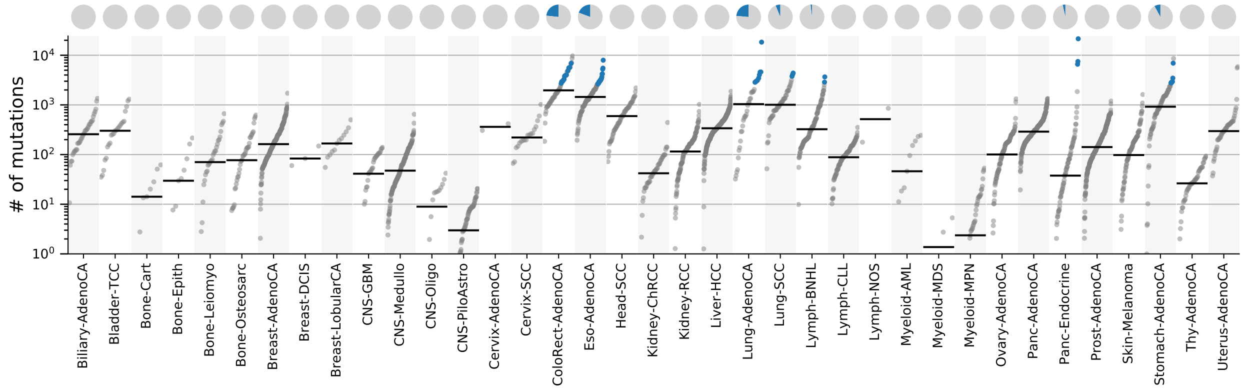

#### 18.5 Signature characterization

Figure 90: Signature specific tensor coefficients.

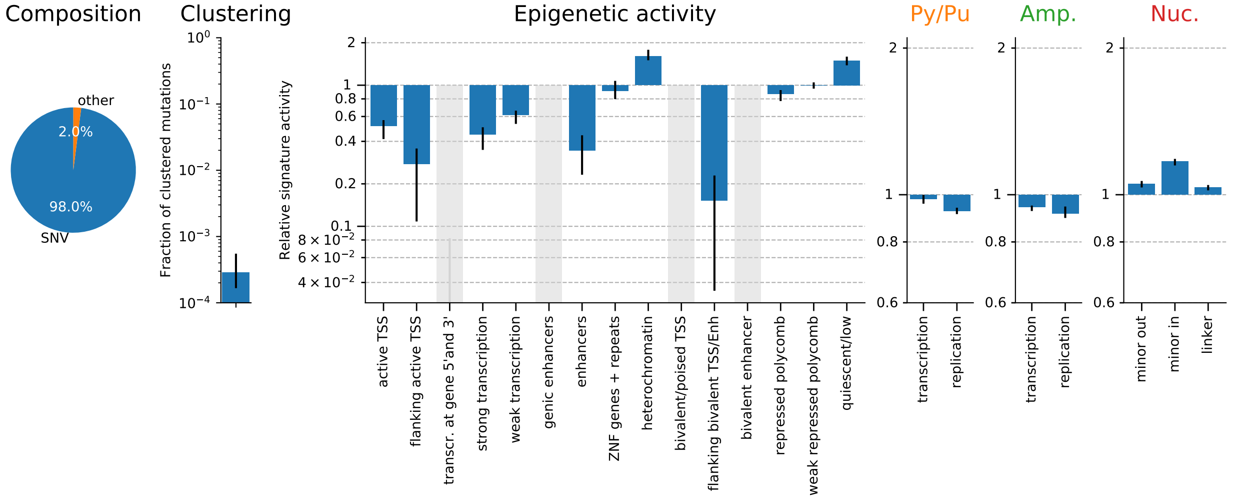

### 19 TS19-N[N>N]N;SV (HRD/BRCA)

#### 19.1 Overall SNV spectrum

Figure 91: Single base substitution spectrum.

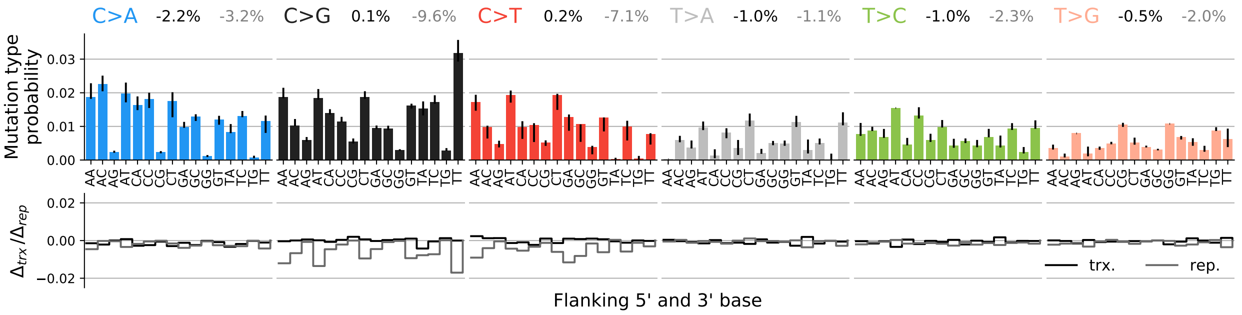

#### 19.2 SNV spectra for transcription and replication

Figure 92: Single base substitution spectra for template/coding and leading/lagging strand DNA.

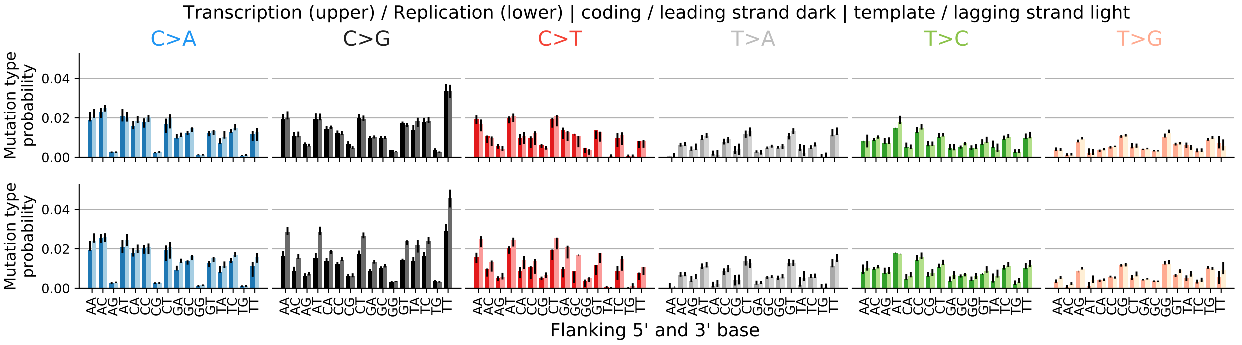

#### 19.3 Spectrum for other mutation types

Figure 93: Spectrum other mutation types.

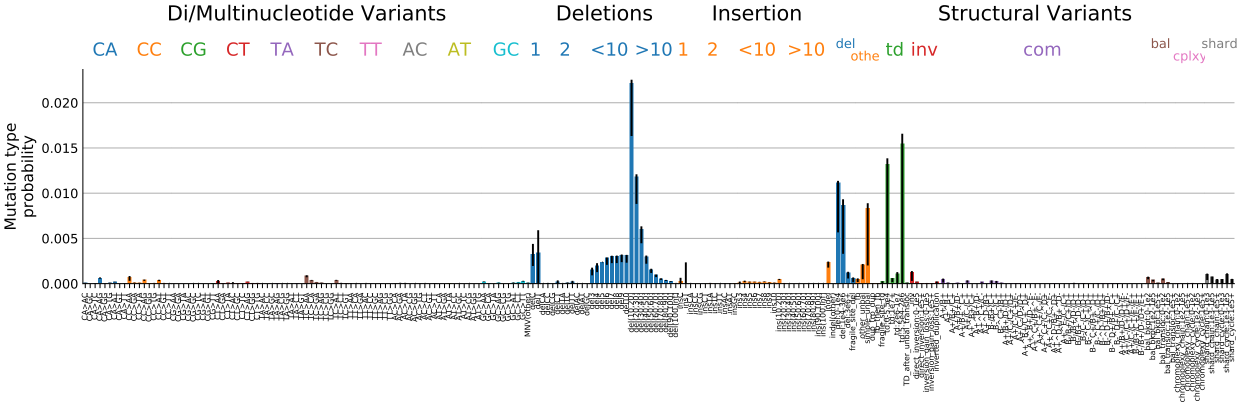

#### 19.4 Signature activity in different cancer types

Figure 94: Signature exposure in different cancer types.

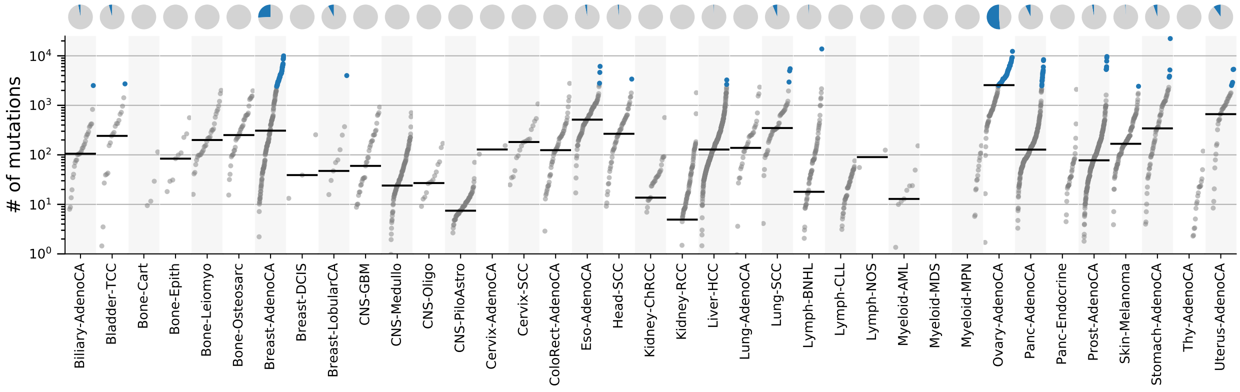

#### 19.5 Signature characterization

Figure 95: Signature specific tensor coefficients.

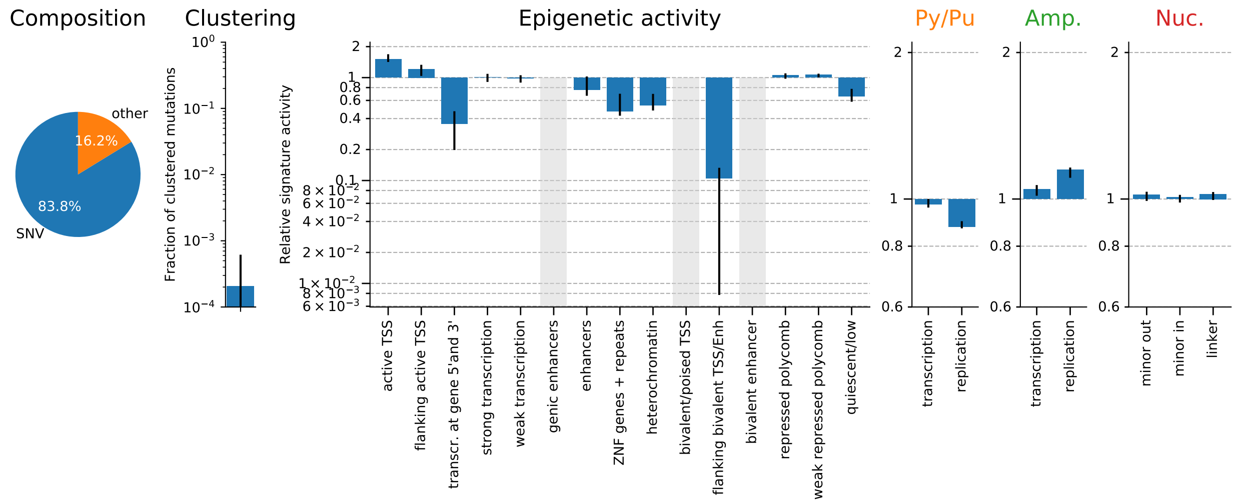

### 20 TS20-N[T>G]T (unknown/5FU)

#### 20.1 Overall SNV spectrum

Figure 96: Single base substitution spectrum.

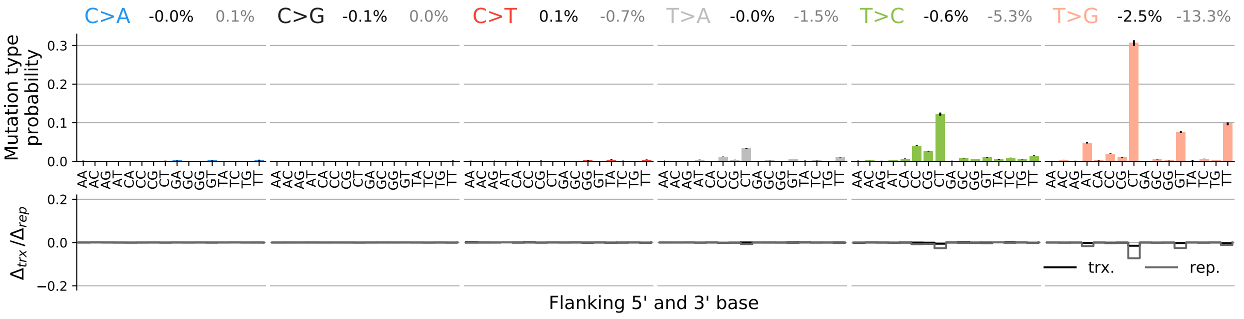

#### 20.2 SNV spectra for transcription and replication

Figure 97: Single base substitution spectra for template/coding and leading/lagging strand DNA.

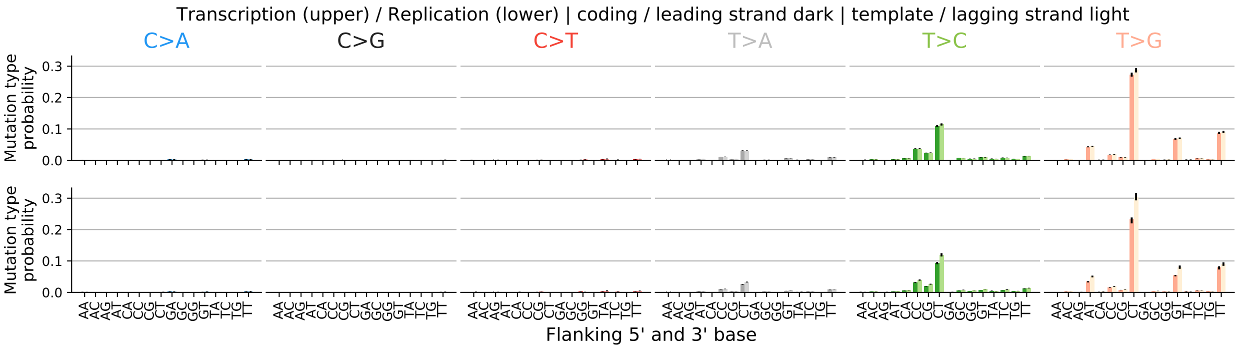

#### 20.3 Spectrum for other mutation types

Figure 98: Spectrum other mutation types.

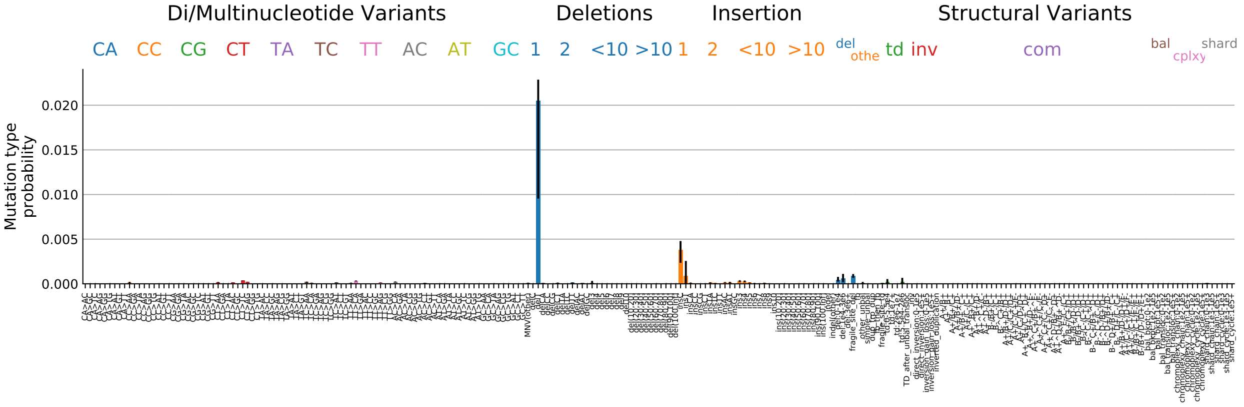

#### 20.4 Signature activity in different cancer types

Figure 99: Signature exposure in different cancer types.

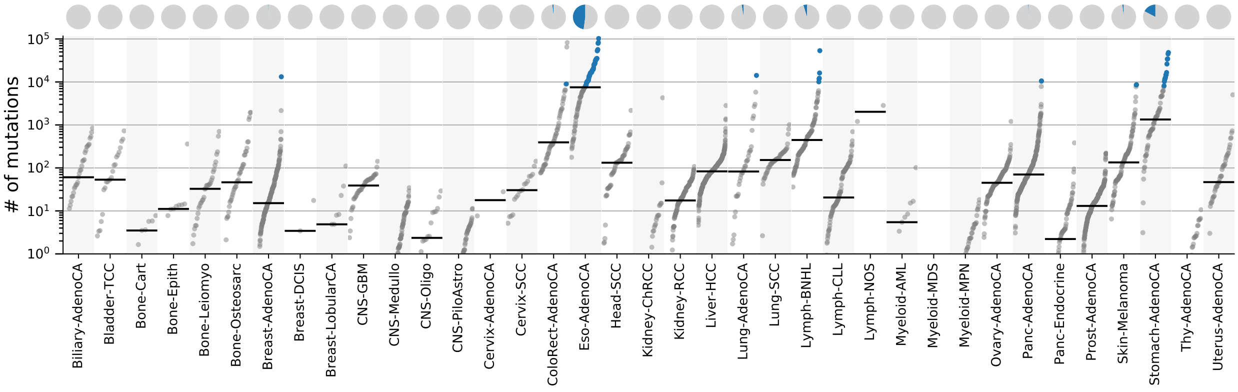

#### 20.5 Signature characterization

Figure 100: Signature specific tensor coefficients.

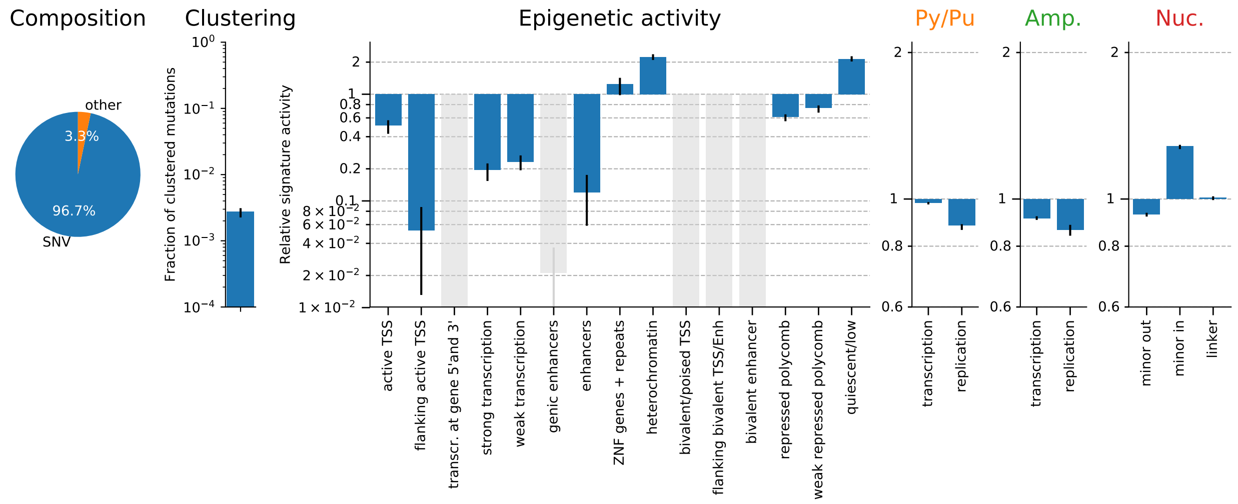

### 21 Tensor factors

#### 21.1 Transcriptional and replicational strand bias

Figure 101: Transcriptional and replicational biases of each signature.

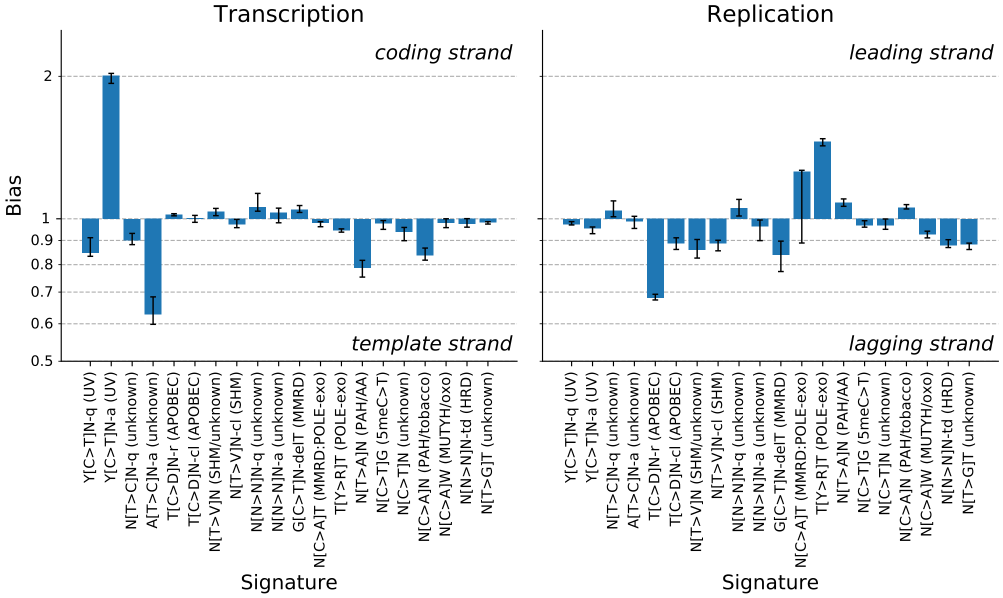

#### 21.2 Transcriptional and replicational amplitudes

Figure 102: Transcriptional and replicational biases of each signature.

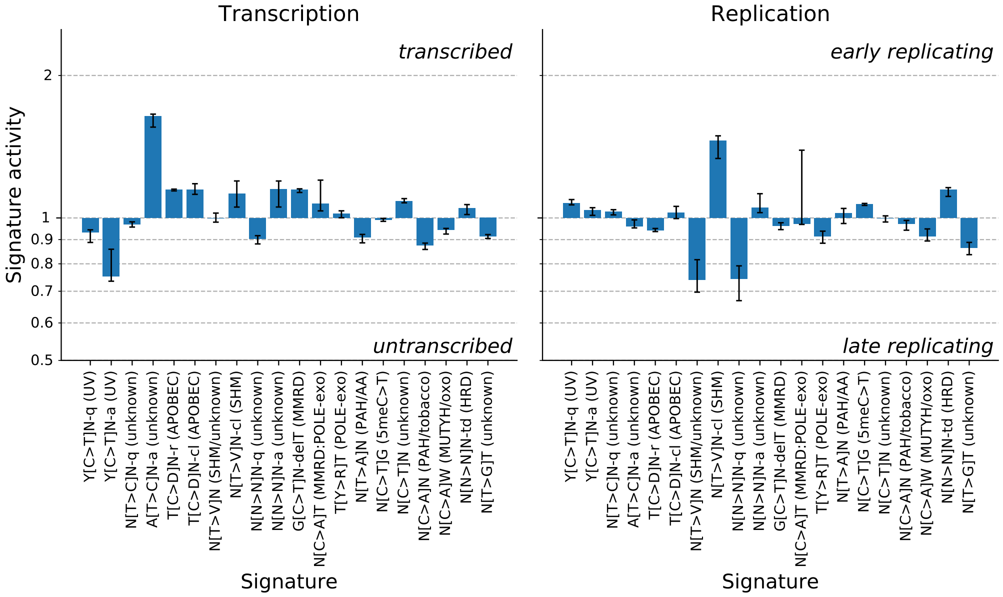

#### 21.3 Nucleosome activities

Figure 103: Nucleosome.

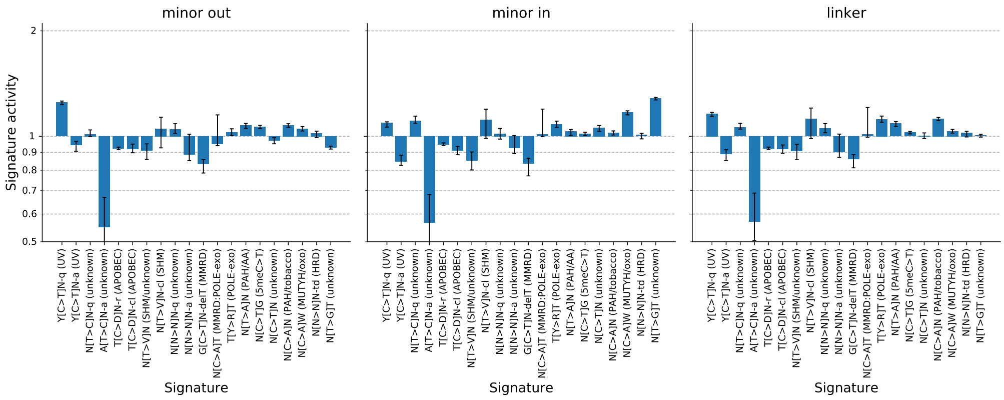

#### 21.4 ChromHMM states

Figure 104: Signature activities across chromHMM states.

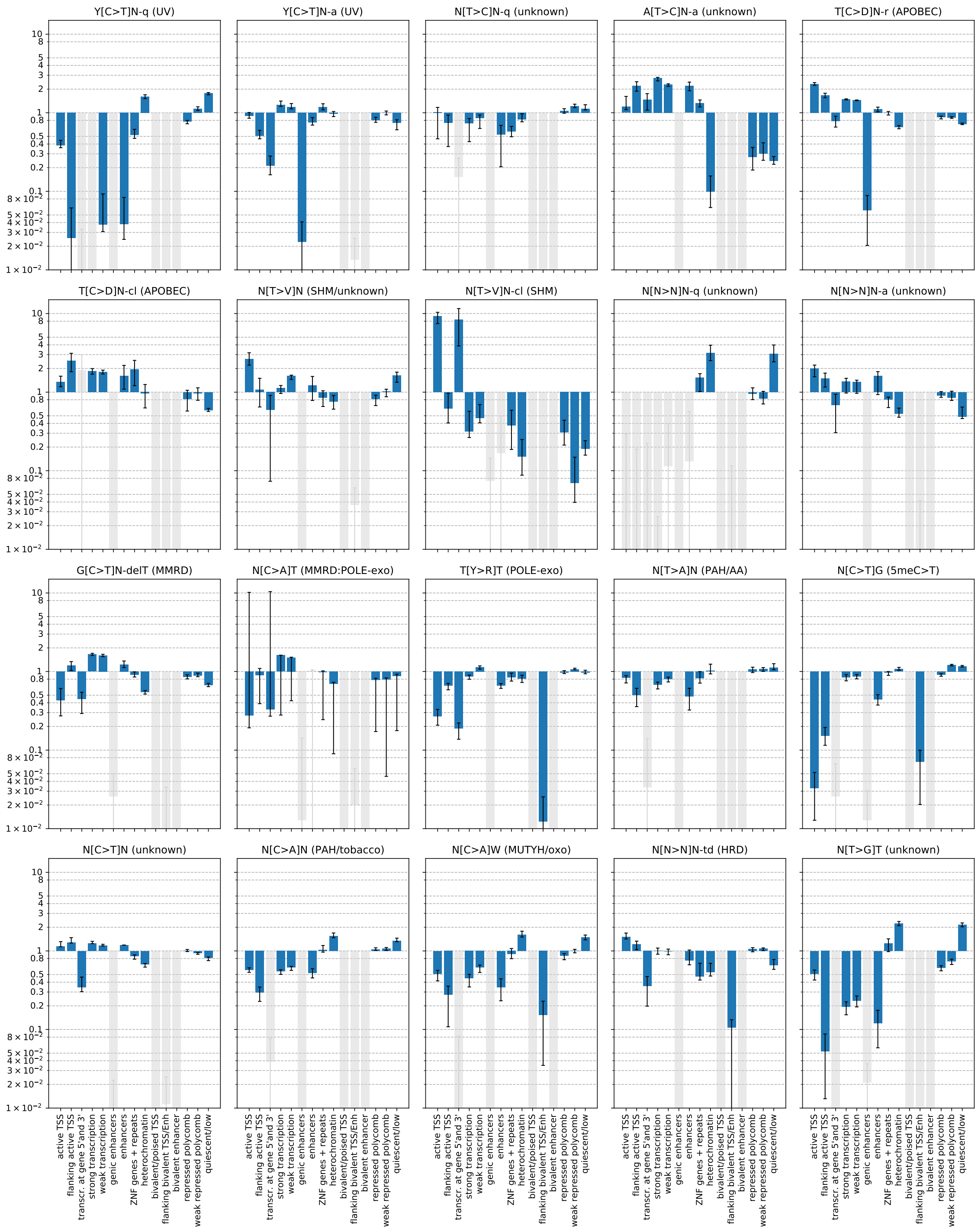
